# Taxogenomic resolution of the 16SrIII (X-disease) group: elevation of ‘*Candidatus* Phytoplasma pruni’-related strains to species rank and proposal of three novel *Candidatus* species ‘*Candidatus* Phytoplasma caryae’ sp. nov., ‘*Candidatus* Phytoplasma conii’ sp. nov. and ‘*Candidatus* Phytoplasma lactucae’ sp. nov

**DOI:** 10.64898/2026.11.16.766873

**Authors:** Usman Bello Rabiu, Sara Wallace, Chelsea L Murphy, Nathan R. Walker, Stephen Marek, Francisco M. Ochoa-Corona, Mustafa O. Jibrin

## Abstract

Alongside the proposal of the taxon *Candidatus* Phytoplasma pruni for 16SrIII (X-Disease) group, the designation “*Candidatus* Phytoplasma pruni-related strains” was proposed for strains that were closely related to the taxon based on multilocus sequence characterization. However, recent genome-based studies have created taxonomic ambiguity by either assigning related strains directly to the taxon *Ca.* P. pruni or separating them into more than one genomic cluster without formal taxonomic classification or proposing new taxon for new divergent strains. In this study, we sequenced genomes of the pecan bunchy top phytoplasma (PBT), previously designated a “*Ca.* P. pruni-related strain,” initially to confirm the pathogen in an Oklahoma pecan orchard. Phylogenomic comparisons with previously sequenced *Candidatus* Phytoplasma pruni strains revealed four well-supported taxonomic groups based on average nucleotide identity, digital DNA–DNA hybridization, and phylogenetic analyses, with different host ecologies consistent with International Research Programme on Comparative Mycoplasmology (IRPCM) guidelines for assigning new taxon. Accordingly, we propose a reclassification of *Ca.* P. pruni and related strains into four species: *Candidatus* Phytoplasma pruni, representing the taxon associated with X-disease phytoplasma; three proposed new *Candidatus* species including *Candidatus* Phytoplasma caryae sp. nov, *Candidatus* Phytoplasma conii sp. nov and *Candidatus* Phytoplasma lactucae sp. nov. The study provides comparative-genomic insights into these taxonomic groups that support their evolution.

## Introduction

Phytoplasmas are obligate intracellular parasitic bacteria (class Mollicutes) that inhabit the phloem tissue of infected plants and are spread between plants mostly by phloem-feeding leafhoppers, planthoppers and psyllids [1]. Phytoplasmas infect many plant species worldwide and have not been cultured in vitro which has historically limited study of their biology, virulence and management [1–4].

Traditionally, phytoplasma taxonomy relies primarily on 16S rRNA sequencing and RFLP profiles [5]. However, genome sequencing of phytoplasma has progressed concurrently with general advancement in genomics and are now included in species delineation requirements. According to IRPCM (International Research Programme on Comparative Mycoplasmology) guidelines [6] and recently updated recommendations, the major criteria required to define a ‘*Ca.* Phytoplasma’ taxa include: (1) a unique 16S rRNA gene sequence (≥1,200 bp) that defines the reference strain and distinguished from closely related strains distinguished by minor sequence differences; (2) if its 16S rRNA gene sequence shares <98.65% identity or its whole genome shares an ANI score <95% to that of any previously described ‘*Ca.* Phytoplasma’ species, or (3) if above this threshold, evidence that the organism represents an ecologically distinct population, demonstrated by differences in insect vectors, plant hosts and supporting molecular divergence [7–9]. Following broad recommendations for genome-based microbial species delineation, a genome coverage of at least 60% was also proposed although this is not always strictly applied [9, 10]. The revised IRPCM rule discourages the use of subspecies, however, there has been debate on the use of “member strain” or “related strain” for strains where only minimal differences exist in the 16S rRNA gene sequence compared to a closely related established taxon [9].

The taxon *Candidatus* Phytoplasma pruni was established using comparison of 16S rRNA gene to describe the strains associated with X disease in peach (*Prunus persica*) and other stone fruits in the 16S rRNA gene RFLP group 16SrIII, subgroup 16SrIII-A [ 6, 8, 11]. In the study formerly establishing the *Ca.* P. pruni taxon, multilocus sequence analysis was used to separate *Ca.* P. pruni strains from other related strains [11]. The descriptive epithet ‘*Ca.* P. pruni-related strains’ was proposed for other closely related 16SrIII strains that cause disease on diverse plant species, with the authors emphasizing that the taxonomic status of these strains could change with studies including more genomic regions [11]. The improved availability of published genomes therefore provides a resource to clarify the taxonomy of *Ca.* P. pruni and *Ca.* P. pruni-related strains [3, 12–15]. Such clarification is needed to provide guidance in assigning taxon for the scientific community, enhance phylo- and comparative genomics, pathogen surveillance and management guidelines for these groups of phytoplasma. In this study, we generated the first draft genomes for the PBT phytoplasma and used phylogenomic and comparative genomics methods to establish the distinction between *Ca.* P. pruni and *Ca.* P. pruni-related strains as well as laying a foundation for surveillance and management of PBT.

### Recent Taxogenomic progress and inconsistencies associated with the nomenclature of 16SrIII group strains

True to the prediction by Davis *et al.* [11], recent genome-based characterization has continued to demonstrate clear taxonomic differentiation of 16SrIII group strains classified as *Ca.* P. pruni and *Ca.* P. pruni-related strains. In a comparative genomic study of *Cicuta* witches’ broom phytoplasma, for example, the average nucleotide identity (ANI), digital DNA-DNA hybridization (dDDH) and maximum likelihood phylogeny shows that the *Cicuta* witches’ broom phytoplasma, a 16SrIII-J strain, belong to the same phylogenetic cluster as *Phytoplasma Vc33*, another 16SrIII-J strain and *China tree decline phytoplasma*, a 16SrIII-B strain, distinct from all other *Ca.* P. pruni and *Ca.* P. pruni-related strains [16]. Similar whole genome comparison demonstrated a clear demarcation of *Lettuce Witches’-Broom phytoplasma*, a 16SrIII-X phytoplasma strain, as a unique cluster that is distinct from all other *Ca.* P. pruni and *Ca.* P. pruni-related strains [14]. Recently, too, whole genome comparison was used to propose a new 16SrIII taxon, ‘*Candidatus* Phytoplasma manihotii’ sp. nov. for the phytoplasma associated with cassava oversprouting disease in Brazil which are 16SrIII-B strains [17]. ‘*Candidatus* Phytoplasma platensis’ was also proposed in a preprint for the 16SrIII-J strain BellVir from daisy virescence [80]. Additionally, the Genome Taxonomy Database (GTDB) [18] contains at least three different clusters for *Ca.* P. pruni and *Ca.* P. pruni-related strains, including clusters designated as *Phytoplasma sp000309485*, *Phytoplasma sp001623385* and *Phytoplasma pruni*, suggesting different groupings for strains in this taxon. However, some recent studies of *Ca.* P. pruni-related strains have defaulted to the use of *Ca.* P. pruni for both *Ca.* P. pruni and *Ca.* P. pruni-related strains, defeating the original aim of Davis *et al.* [11] and further compounding the confusion in classifying members of this *Ca.* P. pruni-related group [3, 11–15]. Our recent studies identified distinct genomic clusters among *Ca.* P. pruni and related strains based on ANI analyses that justify reclassification of *Ca.* P. pruni-related strains into new species [19, 20].

### Pecan bunchy top phytoplasma and the need for taxonomic clarity

Pecan bunchy top phytoplasma (PBT) disease is one of the earliest phytoplasma diseases reported in North America and can be destructive on pecans (*Carya illinoinensis* (Wangenh.) K. Koch) [21–24]. The disease is known to occur sporadically in the Southern US pecan belt and is characterized by branch dieback, witches’ broom, and indeterminate growth of normally dormant axillary buds on slender willowed stems [25–28]. The disease is usually very conspicuous in late spring and early summer [ 28]. No insect vector has been confirmed for PBT, but leafhoppers are widely speculated [26]. Current management guidelines advise removal of infected branches well below visible symptoms as well as destruction of heavily infected trees as severely infected trees rarely recover [26, 27].

PBT has been widely classified as *Candidatus* Phytoplasma pruni in extension notes and are usually grouped with other X-disease phytoplasma group (16SrIII) as 16SrIII-C, although the recommended classification is ‘*Candidatus* Phytoplasma pruni-related strains’ [11]. PBT was grouped with other related strains and classified as *Candidatus* Phytoplasma pruni-related strains based on sequence comparisons of the 16S rRNA, *secY* and ribosomal protein (rp) genes [11]. While describing the taxon, Davis *et al.* [11] noted that the described *Ca.* P. pruni-related strains may not fall into the described taxon, suggesting that sequencing more genetic regions would improve the classification of the *Ca.* P. pruni-related strains in general [11]. Since then, however, there has been limited study of PBT, and no genome-based classification studies exist for the *Ca.* P. pruni-related strains. Alongside other *Ca.* P. pruni-related strains, PBT represents a distinct ecology with its unique host different from *Pruni* species. Clarity of taxonomic status will improve the study of these unique pathosystems.

## Materials and Methods

### Sampling pecan orchard for pecan bunchy top disease

The pecan orchard where PBT symptoms were observed in Oklahoma contained more than 500 trees, with symptomatic trees concentrated in one section displaying irregular growth and canopy decline (Supplementary Figure S1.1). Pecan trees with typical pecan bunchy top symptoms in the orchard were identified and distinctively sampled, using sterilized cutting tool in between trees (Figure 1). While some trees have conspicuous bunch top symptoms that are widespread on tree canopies (Figure 1A), others have bunch symptoms on relatively newly sprouting shoots (Figure 1B). Samples were also collected from asymptomatic pecan trees in the orchard (Figure 1C). All samples were labelled with the prefix “PPS” with a unique number for each sampled tree. Samples were placed on ice, transported to the laboratory and stored at 4 ^0^C until they were processed.

**Figure 1.**
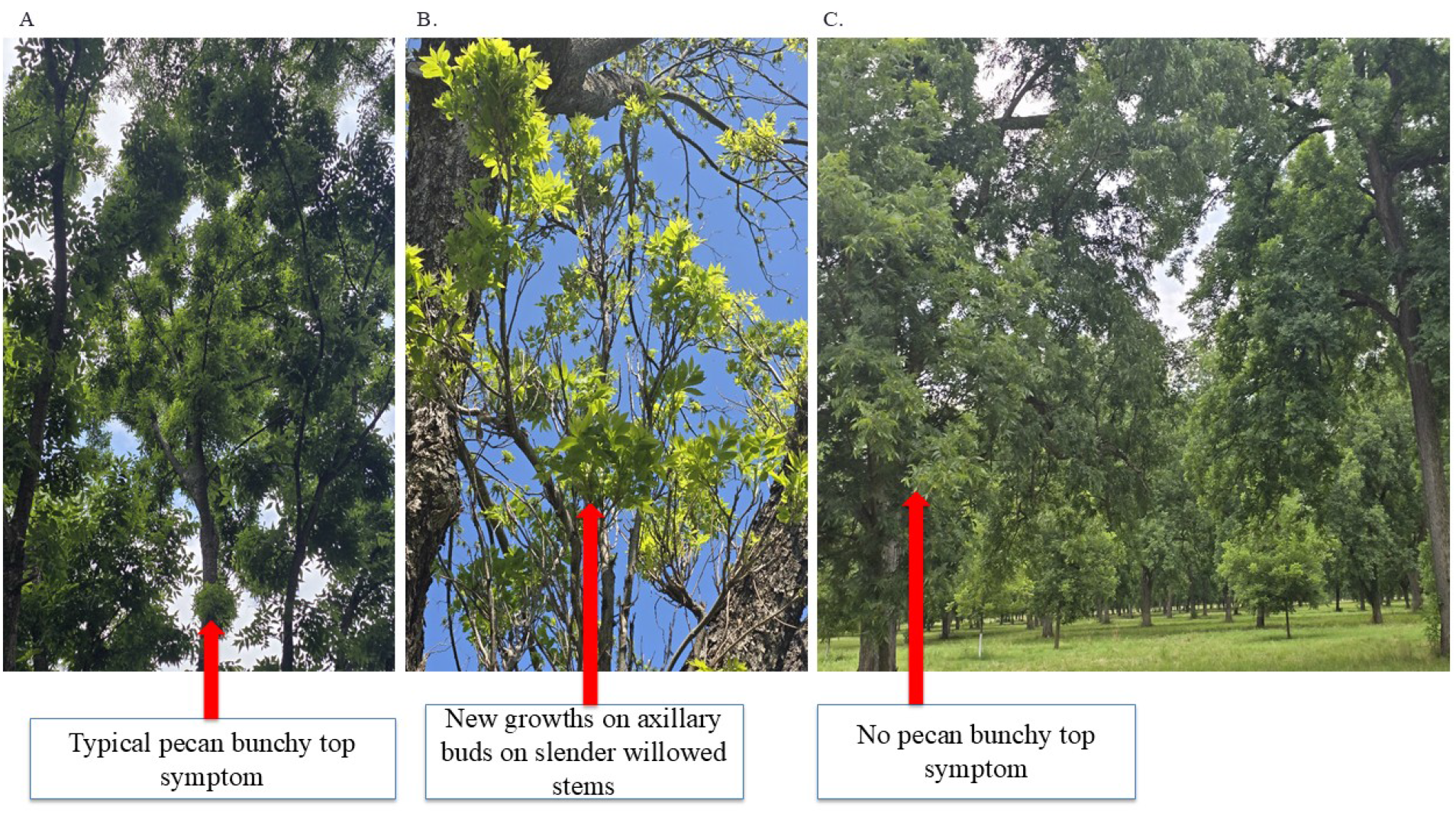
Pecan bunchy top symptoms in the sampled commercial pecan orchard in Oklahoma, USA. (A) Typical bunchy top symptoms on the tree canopy; (B) new growth from axillary buds on slender, willowy shoots; (C) an asymptomatic tree in the same orchard. Arrows indicate the features described.

### Phytoplasma detection and confirmation by PCR

The presence of phytoplasma in pecan samples was confirmed as described previously [29, 30]. Briefly, genomic DNA was extracted from batched samples of individual pecan tree using the Qiagen DNeasy Plant Mini Kit (Qiagen, MD, U.S.A.) following the manufacturer’s instruction. A qPCR assay was carried out using the 18S gene as an internal control and the 23S gene for phytoplasma detection to confirm the presence of phytoplasma [29, 30]. Additionally, a nested PCR of the 16S rRNA gene using primers P1/P7 in the first round [31, 32] and R16F2n/R16R2 in the nested round [33] was carried out as described previously [30]. Nested PCR amplicons were purified and Sanger sequenced at the DNA Sequencing Core Facility at OSU. Sequences were compared with GenBank records by BLASTn to confirm phytoplasma identity prior to selection for whole-genome sequencing.

### Genome sequencing, assembly and annotation

High-quality DNA of six representative strains from different trees were sent for genome sequencing at SeqCenter, LLC (Pittsburgh, PA, USA). Libraries were prepared with the Illumina DNA Prep kit with IDT 10 bp unique dual indices (target insert size 280 bp) and sequenced on an Illumina NovaSeq X Plus (2 x 151 bp, paired-end). Trimming of raw reads and genome assembly followed a combination of recent best practices for assembling phytoplasma genomes [3, 15, 34]. Briefly, the generated reads were demultiplexed and adapter-trimmed with bcl-convert v4.4.4 and filtered them with fastp v1.3.6 [35], discarding read pairs with more than 50 ambiguous bases or failing a base-quality threshold of Q15, and trimming poly-G artifacts of 10 bp or longer. Reads were further trimmed using Trim Galore v0.6.7 with default quality and adapter trimming parameters [36]. Host reads were removed by mapping quality-trimmed reads against the *Carya illinoinensis* cultivar Pawnee reference genome (assembly *C. illinoinensis* Pawnee_v1; NCBI accession GCF_018687715.1) [37] using Bowtie2 v2.4.4 [38]; unmapped reads were used for phytoplasma genome assembly. Quality-trimmed, host-depleted reads were assembled using SPAdes v3.15.5 [39] with the--meta flag invoking the metaSPAdes algorithm [40]. Mean sequencing depth and per-base coverage were calculated by mapping quality-trimmed reads to the assembled phytoplasma genomes using BWA-MEM v0.7.17 [41] and SAMtools v1.10 [42]. Assembly statistics were computed by QUAST v5.3.0 [43]. Annotation was performed using Prokka v1.13 [44]. Genome completeness and contamination were assessed using CheckM v1.2.0 [45].

### Additional Genomes Used in the Study

All 293 unique phytoplasma assemblies available from NCBI (downloaded from https://www.ncbi.nlm.nih.gov/datasets/genome/ by searching “*Candidatus* phytoplasma”), plus the six PBT genomes from this study (n = 299), were assessed with CheckM v1 (lineage_wf). Of the 299 genomes, 252 had a completeness of at least 90% and a contamination of at most 5%. A curated set of 83 genomes used for analyses comprised 62 genomes from a 67-genome base set (the 65 genomes curated by Pellegrinetti *et al.* [15] plus two additional genomes), the six PBT genomes (completeness 90.0 to 97.3%, contamination 0%), seven additional reported *Ca.* P. pruni and related genomes, and eight genomes added so that every organism name with a passing assembly was represented. Five genomes designated as *Ca.* P. pruni strains were excluded because they did not meet these criteria in our CheckM run: OY-V (completeness 89.5%), NJAY (89.2%), SA213 (81.7%), NCHU2019 (contamination 9.6%) and ChTYXIII-Mo (contamination 15.9%). Vc33 (completeness 83.6%) was excluded from the curated set but was retained in the ANI and dDDH comparisons of the 22 PBT, *Ca.* P. pruni and *Ca.* P. pruni-related genomes.

### Multilocus sequence comparison and 16S rRNA subgroup classification

Pairwise comparison for full-length genes was conducted to determine nucleotide differences in selected gene sequence including 16S rRNA, *rpoB*, *secA*, *secY*, *rps3*, *rps19* and *rpl22* of genomes of *Ca.* P. pruni and *Ca.* P. pruni-related strains using PPS1A as reference. Nucleotide identity was calculated for 16S rRNA, *rpoB*, *secA*, *secY*, *rps3*, *rps19* and *rpl22*. Protein alignments of the marker genes were converted to codon alignments using the nucleotide sequences. Identity was the number of matching nucleotides divided by the number of nucleotides compared, using alignment columns with A, C, G or T in both sequences, and positions are given in PPS1A gene coordinates. There was a frameshift in the CX *rpoB* which was masked, and the PPS10A *secY* sequence was shortened by a contig end (413 of 423 residues).

The 16Sr group and subgroups classification was assessed using iPhyClassifier [5] applied to the full length 16S rRNA gene sequences recovered from the assembled genomes in this study as well as other previously sequenced genomes in *Ca.* P. pruni to confirm their ribosomal group assignments.

### Average Nucleotide Identity, Digital DNA-DNA Hybridization (d4) and Genome Taxonomy Database (GTDB) Clustering

Average nucleotide identity (ANI) was computed at the following levels: PBT strains only (n = 6), PBT strains plus all *Ca.* P. pruni and *Ca.* P. pruni-related (n =22), a curated list of phytoplasma genomes with CheckM values above 90 (n=83), and all phytoplasma genomes downloaded from https://www.ncbi.nlm.nih.gov/datasets/genome/ by searching “*Candidatus* phytoplasma” (n= 299) using FastANI v1.34 [46]. Pairwise ANI among the 22 genomes of the PBT, *Ca.* P. pruni and *Ca.* P. pruni-related strains (231 pairs) were additionally computed with pyANI v0.2.12, using ANIm (MUMmer4) [47] and ANIb (BLAST+ v2.16.0) [48]. Genomes were grouped by average-linkage hierarchical clustering of the ANI matrices at 95% identity, and the sensitivity of the grouping was tested with single, average and complete linkage at 94%, 95% and 96% identity. *Acholeplasma laidlawii* PG-8A was included as an outgroup because it is the closest culturable relative of phytoplasmas and the conventional root for phytoplasma phylogenies [49]. FastANI heatmaps were generated in R v4.5.2 [50] using the ggplot2 package [51]. Digital DNA-DNA hybridization (dDDH) was calculated based on the Genome-to-Genome Distance Calculator (GGDC) Formula, d_4_, which uses High-Scoring Segment Pair (HSP) length and is independent of genome length and is recommended for incomplete genomes [52, 53]. A dDDH value above 70% was taken as the species boundary [52]. The dDDH was calculated with the GGDC web server using the BLAST+ option and default settings for all pairs among the 22 genomes, and formulas 1 and 3 were compared with formula 2.

Additional genome-based classification was conducted using the Genome Taxonomy Database Toolkit, GTDB-Tk v2.6.1, to identify the classification of the *Ca.* P. pruni and *Ca.* P. pruni-related in genome-based databases [18]. GTDB-Tk v2.6.1 (classify_wf) was run against GTDB Release r226 on 53 genomes: the 22 *Ca.* P. pruni and *Ca.* P. pruni-related genomes and 31 other phytoplasma genomes.

### Phylogenetic Reconstructions

Phylogenetic reconstructions using individual marker genes were conducted using 16S rRNA, *secY*, *rp*, *tuf gene* sequences as described previously [11]. Genes were extracted from each assembled genome by BLASTn (v2.13.0) homology search against a curated reference database of phytoplasma marker gene sequences representing multiple 16Sr groups [54]. Extracted sequences were aligned using MAFFT v7.525 [55]. Maximum likelihood trees were inferred independently for each marker gene using IQ-TREE2 v2.4.0 [56] with ModelFinder (-m MFP) selecting the best-fit substitution model under the Bayesian Information Criterion (BIC), with 1,000 ultrafast bootstrap replicates, rooted on *Acholeplasma laidlawii* PG-8A.

Core genome phylogeny was inferred for different groups of strains using core genome alignment output from pangenome analyses performed using Panaroo v1.7.0 [57]. Initially, pangenome analysis was performed using the PBT strains, *Ca.* P. pruni reference strains, and other strains outside *Ca.* P. pruni group III strains (Table S1.4). Maximum likelihood tree was inferred using IQ-TREE2 v2.4.0 with ModelFinder (-m MFP) and 1,000 ultrafast bootstrap replicates [56]. Phylogenetic trees were visualized and annotated using the Interactive Tree of Life (iTOL) v6 [58].

Subsequently, coding sequences were taken from the NCBI annotations (RefSeq or GenBank) of the 83 curated genomes and from Prokka annotations (genetic code 11) of the six PBT genomes and the two OVSP genomes, and were translated with NCBI translation table 11. Pseudogenes with internal stop codons and proteins shorter than 30 residues were excluded. A first pangenome analysis with Panaroo (identity threshold 0.7, sensitive mode) on the 83 genomes and the outgroup *A. laidlawii* PG-8A produced 12,616 gene families, of which only 4 were present in at least 95% of the genomes. This shows the divergence between 16Sr groups: the median identity of reciprocal best BLASTP hits between PPS1A and representatives of other groups was 55.4 to 61.3% (48.5% to *A. laidlawii*), below the clustering threshold. Orthology was therefore inferred with OrthoFinder v3.1.5 [59] using DIAMOND and MCL on all 84 proteomes, which produced 1,536 orthogroups with at least two genes. Of these, 103 were present in at least 95% of the phytoplasma genomes, had at most one copy per genome and contained one *A. laidlawii* copy. Each was aligned with MAFFT (--auto), alignment columns with more than 50% gaps were removed, and the 92 orthogroups that retained at least 80 columns were concatenated into a supermatrix of 26,970 amino acid sites. A maximum-likelihood tree was inferred with IQ-TREE v2.4.0 (ModelFinder restricted to the LG, WAG, JTT and Q.pfam families; best-fit model JTT+F+R6 by BIC) with 1,000 ultrafast bootstrap replicates.

### Comparative Genomics Analyses

Pangenome analyses for the three identified groups with multiple strains and another for the PBT strains were conducted using Panaroo v1.7.0 [57]. Pangenome openness was assessed using Heap’s Law, fitting the observed gene cluster count to the power law equation F = kN^alpha [60]. Mean pan-genome size over random genome orderings was regressed on the number of genomes on a log-log scale (pan-genome size = k × n^alpha). Recombination was inferred on the core genome of the three groups using ClonalFrameML [61].

Additional comparative genomics utilized all *Ca.* P. pruni and *Ca.* P. pruni-related strains. Comparative genomics was conducted to identify the Potential Mobile Unit (PMU) cluster, metabolic pathway, COG annotation, and secretome prediction. Contig identifiers, gene coordinates, strand orientation, and copy number were recorded for each matched gene to evaluate cluster integrity and gene order across genomes. Metabolic pathway gene content was assessed for seven curated pathways: nucleotide salvage, glycolysis, DNA replication, folate biosynthesis, glycerophospholipid biosynthesis, protein secretion, and transport. Gene presence or absence in each pathway was determined by parsing Prokka-generated GFF annotation files for gene name and product field matches against a curated list of 66 pathway-specific gene identifiers compiled from published phytoplasma metabolic reconstructions and BLAST [62]. The effectors were further compared using a curated set of effectors, including TENGU, SAP05, SAP06/48-like, SAP11, SAP54, SAP54-like, SVM_705, SVM_1019, SVM_1100, SVM_1102 and SVM_1103 [49]. Generally, genes not detected in the Prokka annotation were further searched using BLASTp v2.13.0 with protein sequences of the corresponding genes from annotated reference phytoplasma genomes as queries against the predicted proteomes of all PBT strains and all *Ca.* P. pruni and *Ca.* P. pruni-related strains at an e-value threshold of 1e-3. Genes returning no BLASTp hit at this threshold in any genome were scored as absent. Besides Prokka-based annotation and BLASTp, functional annotation of predicted proteins was performed using BlastKOALA [63] via the KEGG web server (https://www.kegg.jp/blastkoala/) to assign KEGG Orthology (KO) numbers and map genes to KEGG metabolic pathways. COG functional category annotation was performed by BLASTp comparison of predicted protein sequences against the COG2020 database [64] using DIAMOND v2.2.1 (e-value threshold 1×10⁻⁵) [65]. Comparative orthogroup analysis was performed using OrthoFinder v3.1.5 [59] on predicted protein sequences from all PBT strains and six *Ca.* P. pruni reference strains to identify PBT-exclusive and reference-exclusive orthogroups. Signal peptide prediction was performed on Prokka v1.13-predicted protein sequences using SignalP v5.0b [66] with organism type gram-positive (-org gram+); proteins with predicted signal peptide class SP(Sec/SPI) were classified as secreted candidates. Conservation of secreted protein candidates across *Ca.* P. pruni reference strains was assessed by BLASTp comparison against the proteomes of the reference genomes using DIAMOND v2.1.13 [65] with an e-value threshold of 1×10⁻⁵ and minimum sequence identity of 30%.

## Results and Discussion

### Confirmation of PBT, Genome assembly, annotation, and quality assessment

The presence of pecan bunchy top phytoplasma was confirmed in recovered samples by both qPCR and BLAST of nucleotide sequence products of the 16S rRNA gene in six samples from six different trees (PPS1A, PPS2B, PPS6A, PPS6B, PPS9B, PPS10A). The 16S rRNA gene sequences were deposited in the NCBI database under the following accession numbers: QB316898-QB316903 for strains PPS1A-PPS10A respectively. The 16Sr group and subgroups classification, assessed using iPhyClassifier [5] applied to the full length 16S rRNA gene sequences, classified all PBT strains as 16SrIII-C strains (Figure S1.2). Although PBT is widely inferred to be present in Oklahoma due to its conspicuous symptoms [26, 28], this represents the first confirmation of PBT in the state.

Six draft phytoplasma genomes were assembled from quality-filtered Illumina paired-end sequencing reads. Reads mapping to the phytoplasma genome constituted only 1.41% (PPS9B) to 4.50% (PPS6B) of the total libraries (Table 1). This low recovery is due to the high abundance of host (*Carya illinoinensis*) DNA inherent in culture-independent metagenomic sequencing of infected tissues performed without prior pathogen enrichment [67]. The resulting assemblies ranged in size from 510,648 bp (strain PPS10A) to 630,190 bp (strain PPS6A). Assembly contiguity varied among the strains. PPS6B yielded the most contiguous assembly (N50 = 40,330 bp; 41 contigs), followed by PPS1A (N50 = 29,166 bp; 46 contigs) and PPS6A (N50 = 26,154 bp; 65 contigs). PPS9B and PPS10A assemblies showed higher fragmentation, with N50 values of 4,036 bp (160 contigs) and 3,156 bp (188 contigs), respectively. This fragmentation correlates directly with their reduced mean sequencing depths of 14.6× and 12.6× for PPS9B and PPS10A respectively (Table 1). Strain PPS6B, with sequencing depth of 76.3x, has the highest genome coverage while genome completeness ranged from 90.01% (PPS10A) to 97.26% (PPS1A, PPS2B, PPS6A, PPS6B), with 95.21% for PPS9B (Table 1; Supplementary Figure S1.3). The GC content ranged from 26.96% to 27.65% across all six strains (Table 1), consistent with the low GC content typical of phytoplasma genomes [14, 62]. Whole genome synteny comparisons showed various degrees of collinear alignments among sequenced strains and between previously sequenced *Ca.* P. pruni and *Ca.* P. pruni-related strains (Fig. S1.4-1.7).

**Table 1.**
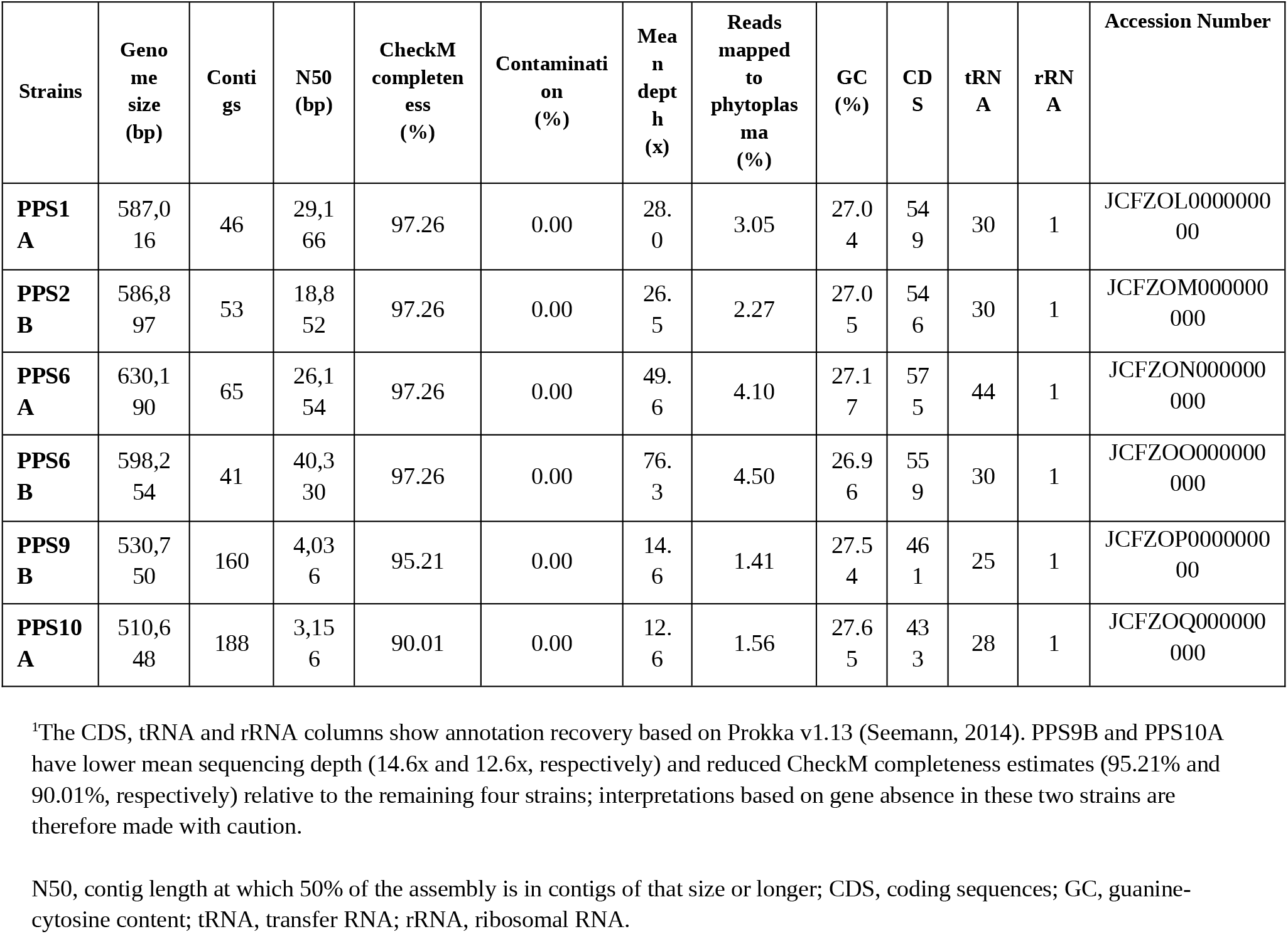
Assembly statistics, genome quality metrics and annotation for the six *Ca.* Phytoplasma caryae (16SrIII-C) strains causing pecan bunchy top disease in Oklahoma in 2025^1^.

### Single copy gene comparison

Pairwise comparison for full-length genes initially conducted to determine nucleotide differences in selected gene sequence including 16S rRNA, *rpoB*, *secA*, *secY*, *rps3*, *rps19* and *rpl22* of genomes of *Ca.* P. pruni and *Ca.* P. pruni-related strains using PPS1A produced varying results. Strains PDA15, Vc33 and ZT3-1 do not have any 16S rRNA gene while strains CanS4, PR2021, CicWB-2022, 6A1, OVSP-L01, OVSP-S01 and LWB have two copies of the gene (Tables S2.1–S2.5). Comparison using the 16S rRNA gene shows that most *Ca.* P. pruni and *Ca.* P. pruni-related strains are above the 98.65% similarity set by the IRPCM rule c guideline (Table 3; Figs. S2.1–S2.5). However, strain ChTDIII and one of the two 16S rRNA gene copies of OVSP-L01 have less than 98.65% identity in the 16S rRNA gene with other strains (Fig. S2.1; Tables S2.2–S2.5). So, based on the IRPCM rule c, the 16S rRNA gene alone does not differentiate these groups of strains. Compared with the 60 described ‘*Candidatus* Phytoplasma’ species that have a 16S rRNA gene sequence in GenBank, the reference 16S rRNA gene sequences of *Ca.* P. caryae, *Ca.* P. conii and *Ca.* P. lactucae are 99.0–99.3% identical to *Ca.* P. pruni (JQ044393) and 95.1% or less identical to every other species (Table S2.6). Species delimitation therefore relies on ANI, dDDH and phylogenomic analysis (Tables 3 and 4; Figs. 2, 3 and 5). However, significant differences were detected in the following genes: *rpoB, secA, secY,* and *rps3* (Table 3; Fig. S2.8).

**Figure 2.**
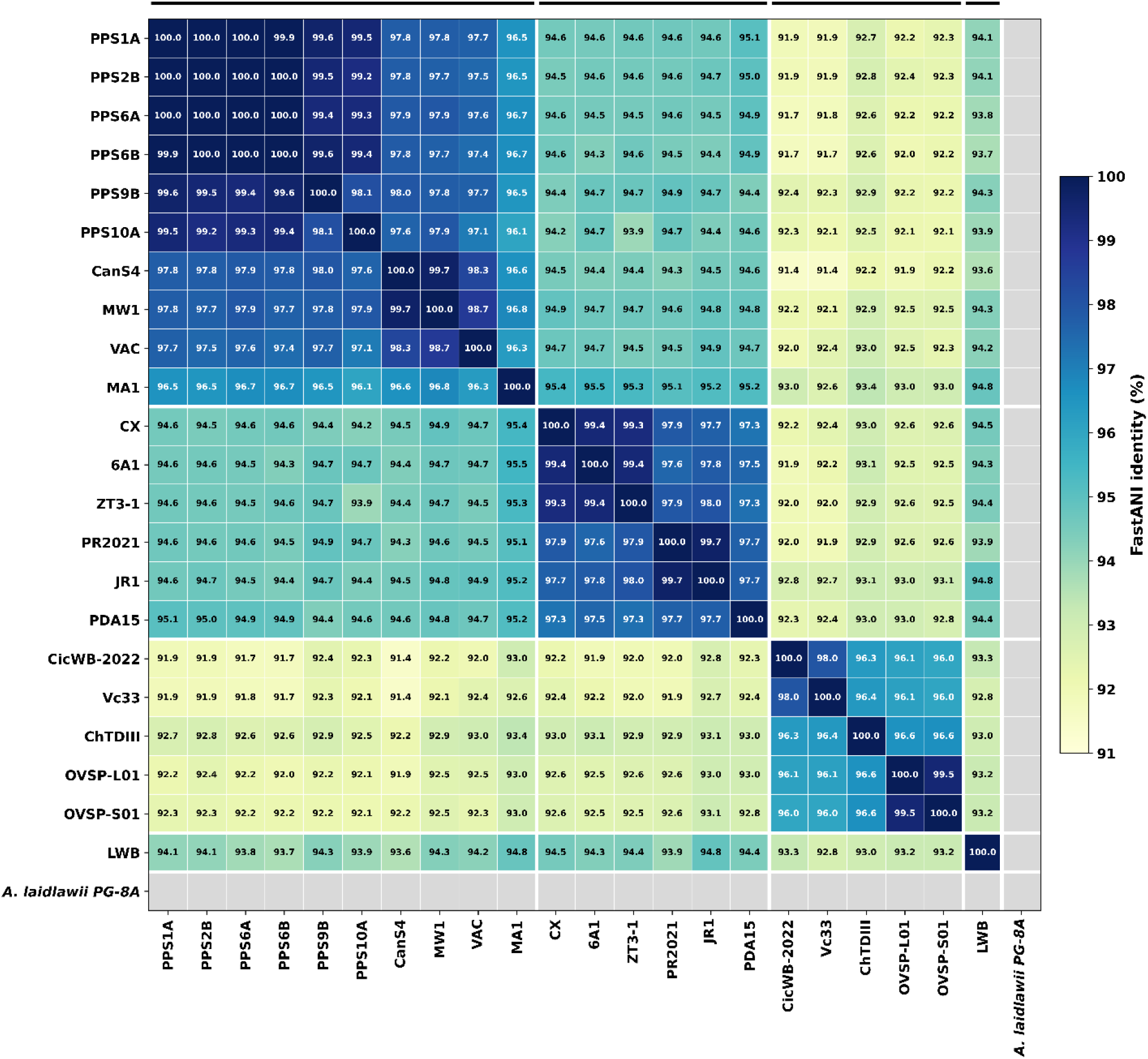
Pairwise average nucleotide identity (ANI) among the 22 *Ca.* P. pruni and *Ca.* P. pruni-related genomes, calculated with FastANI v1.34 [46]. Values are the mean of the two directional estimates and are shown in each cell. White lines separate the four ANI groups: *Ca.* P. caryae (Group 1), *Ca.* P. pruni (Group 2), *Ca.* P. conii (Group 3) and *Ca.* P. lactucae (Group 4). *Acholeplasma laidlawii* PG-8A was included as the outgroup; FastANI returned no values for it (grey cells).

**Figure 3.**
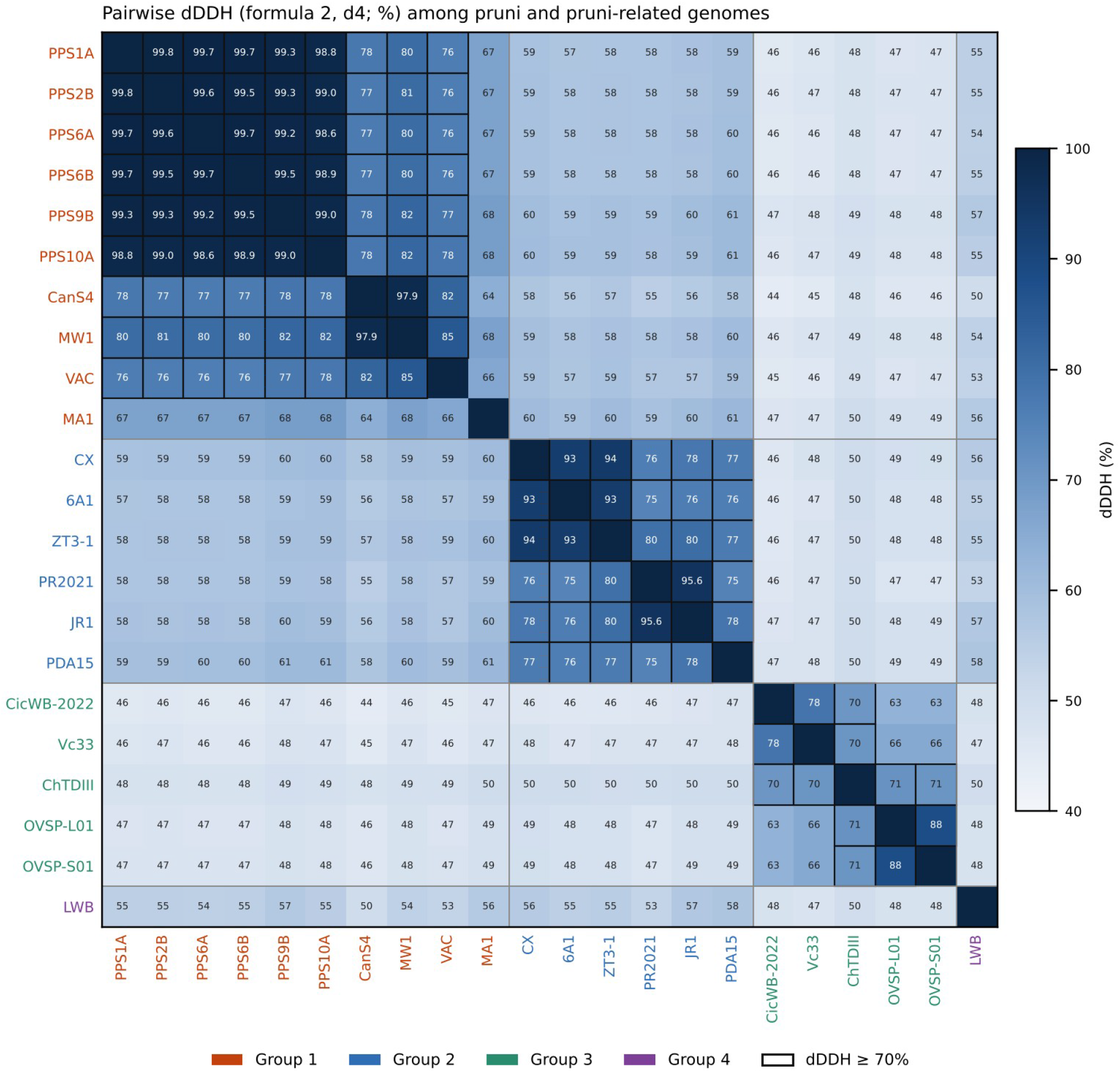
Pairwise digital DNA–DNA hybridization (dDDH) among the 22 genomes, calculated with the Genome-to-Genome Distance Calculator formula 2 (d4). Values are shown in each cell, and black outlines mark pairs with dDDH of 70% or more, the species threshold. Strain labels are coloured by ANI group (Group 1, orange-red; Group 2, blue; Group 3, green; Group 4, purple).

### Average Nucleotide Identity (ANI), digital DNA-DNA Hybridization (dDDH) Analyses and Genome Taxonomy Database (GTDB) Clustering

Pairwise FastANI values across the six PBT strains ranged from 98.11 to 99.96% (Fig. 2, S3.1). ANI analyses of genomes of strains within *Ca.* P. pruni and *Ca.* P. pruni*-*related strains showed clear separation into four groups of strains based on <95–96% species limit for phytoplasma (Fig. 2, S3.1; Tables S3.1–S3.5). Based on the ANI threshold, all *Ca.* P. pruni and *Ca.* P. pruni*-*related strains fall into four groups. These groups include group 1 (the PPS group) consisting of PPS1A-PPS10A (the PBT strains), CanS4, MW1, VAC and MA1; group 2 (the CX group) consisting of strains CX, JR1, 6A1, ZT3-1, PR2021 and PDA15; group 3 (the CicWB-2022 group) consisting of CicWB-2022, ChTDIII, Vc33, OVSP-L01 and OVSP-S01; and group 4 (the LWB group) which consist only the LWB strain (Fig. 2, S3.1; Table S1.3). Each of these groups share less than 96% with each other (Table 4). Two strains, PDA15 and MA1, have slight intermediate values. Strain PDA15 shares 95.1% ANI with PPS1A but shares less than 95% with all other strains in the PPS group. Similarly, strain MA1 is close to group 2 strains by sharing 95.1–95.5% ANI but shares above 96% ANI with all group 1 strains.

Digital DNA-DNA hybridization (dDDH) was calculated based on the Genome-to-Genome Distance Calculator (GGDC) Formula, d_4_, which uses High-Scoring Segment Pair (HSP) length and is independent of genome length and is recommended for incomplete genomes [52, 53]. A dDDH value above 70% was taken as the species boundary [52]. The dDDH was calculated with the GGDC web server using the BLAST+ option and default settings for all pairs among the 22 genomes, and formulas 1 and 3 were compared with formula 2. The result of the ANI is further supported by the dDDH analyses (Figs. 3, S3.2–S3.3; Tables S4.1–S4.7). Each of the four groups share less than 70% dDDH values with each other and above 70% with strains within the same group (Figs. 3, S3.2–S3.3; Tables S4.1–S4.7). The exceptions are strain MA1, which shares less than 70% dDDH with the other group 1 strains (64.2–68.5%) but is still closer to group 1 than to group 2 (58.9–61.4%), and group 3, in which CicWB-2022 and Vc33 share 63.3–65.7% dDDH with OVSP-L01 and OVSP-S01, while ChTDIII shares 70.4–71.2% with the other group 3 strains (Tables S4.3, S4.5). No other strains within the 83 curated strains nor the 299 genomes downloaded from the entire list of phytoplasma genomes from NCBI shares ANI or dDDH values with these *Ca.* P. pruni and *Ca.* P. pruni*-*related strains.

GTDB clustering was achieved by running GTDB-Tk v2.6.1 [18] against GTDB Release r226 on 53 genomes: the 22 *Ca.* P. pruni and *Ca.* P. pruni-related genomes and 31 other phytoplasma genomes. GTDB-Tk v2.6.1 placed all *Ca.* P. pruni and *Ca.* P. pruni*-*related strains into three groups, with no cluster for strain LWB (Table S5.1). The groups 1, 2 and 3 strains were assigned to *Phytoplasma sp000309485*, *Phytoplasma pruni* and *Phytoplasma sp001623385* respectively, further supporting the ANI and dDDH results of different taxonomic groupings for these strains (Tables S5.1–S5.3).

### Phylogenetic reconstructions

Maximum likelihood phylogenetic analysis based on the 16S rRNA gene placed all *Ca.* P. pruni and *Ca.* P. pruni*-*related strains within different clusters, but the bootstrap support value (57) suggests that the 16S rRNA gene is not sufficient to separate them from other *Candidatus* phytoplasma species (Fig. 4). Maximum likelihood phylogenetic analyses based on *rp, rpoB, secY*, *tuf* and concatenation of *rpoB, secA, secY* and 20 ribosomal protein genes placed all *Ca.* P. pruni and *Ca.* P. pruni*-*related strains within well-supported four monophyletic clades for *Ca.* Phytoplasma pruni (16SrIII group) with high bootstrap support across all four phylogenies (Figs. S4.1–S4.5).

**Figure 4.**
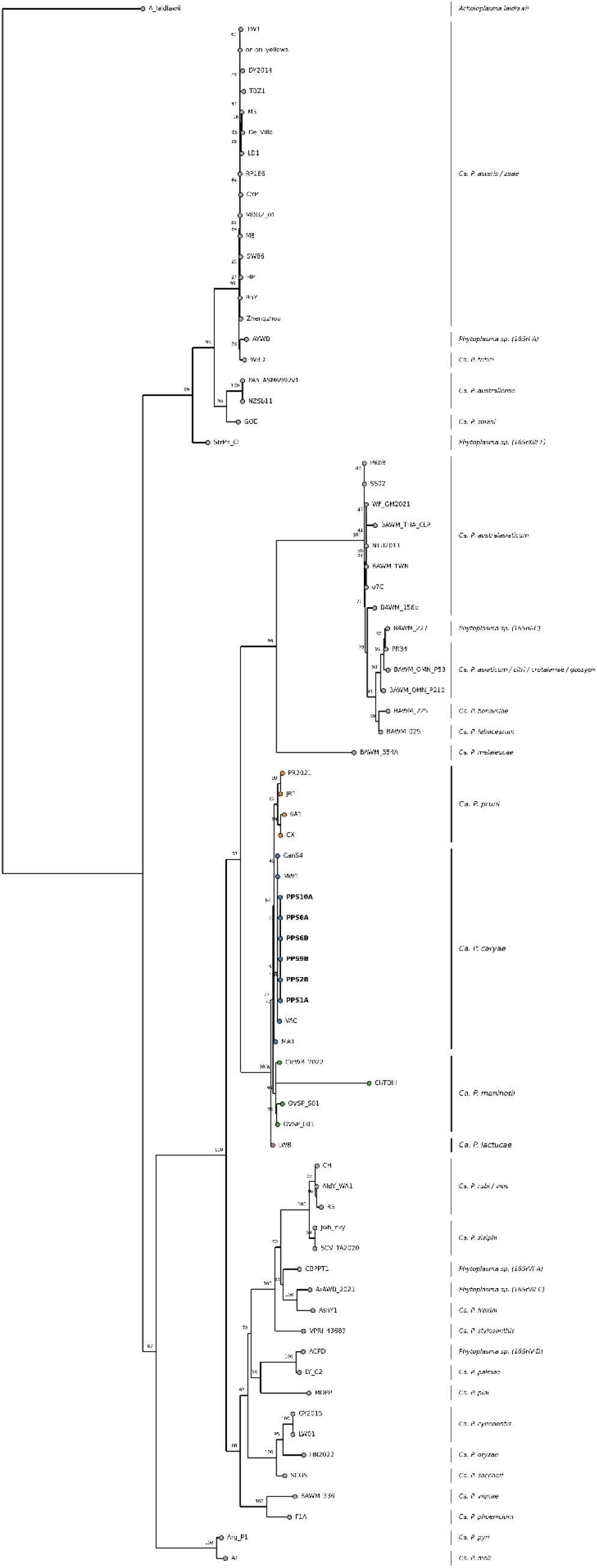
Maximum-likelihood phylogeny of the 16S rRNA gene of the *Ca.* P. pruni and *Ca.* P. pruni-related strains and representative ‘*Candidatus* Phytoplasma’ species, inferred with IQ-TREE2 v2.4.0 with 1,000 ultrafast bootstrap replicates [79] and rooted on *Acholeplasma laidlawii*. Tip colours indicate the ANI groups (Group 1, blue; Group 2, orange; Group 3, green; Group 4, purple).

Core genome phylogeny was inferred for the 83 curated genomes and *A. laidlawii* PG-8A as an outgroup using 3300-nucleotide core gene alignment output from pangenome analyses performed using Panaroo v1.7.0 [57]. Maximum likelihood phylogeny based on alignments of the concatenated core genes separated the *Ca.* P. pruni and *Ca.* P. pruni*-*related strains into four well-supported clades (Fig. 5, S4.6). Accordingly, we propose the reclassification of *Ca.* P. pruni and *Ca.* P. pruni*-*related strains into four species as *Candidatus* Phytoplasma caryae, *Candidatus* Phytoplasma pruni, *Candidatus* Phytoplasma conii and *Candidatus* Phytoplasma lactucae representing ANI groups 1, 2, 3 and 4 respectively.

**Figure 5.**
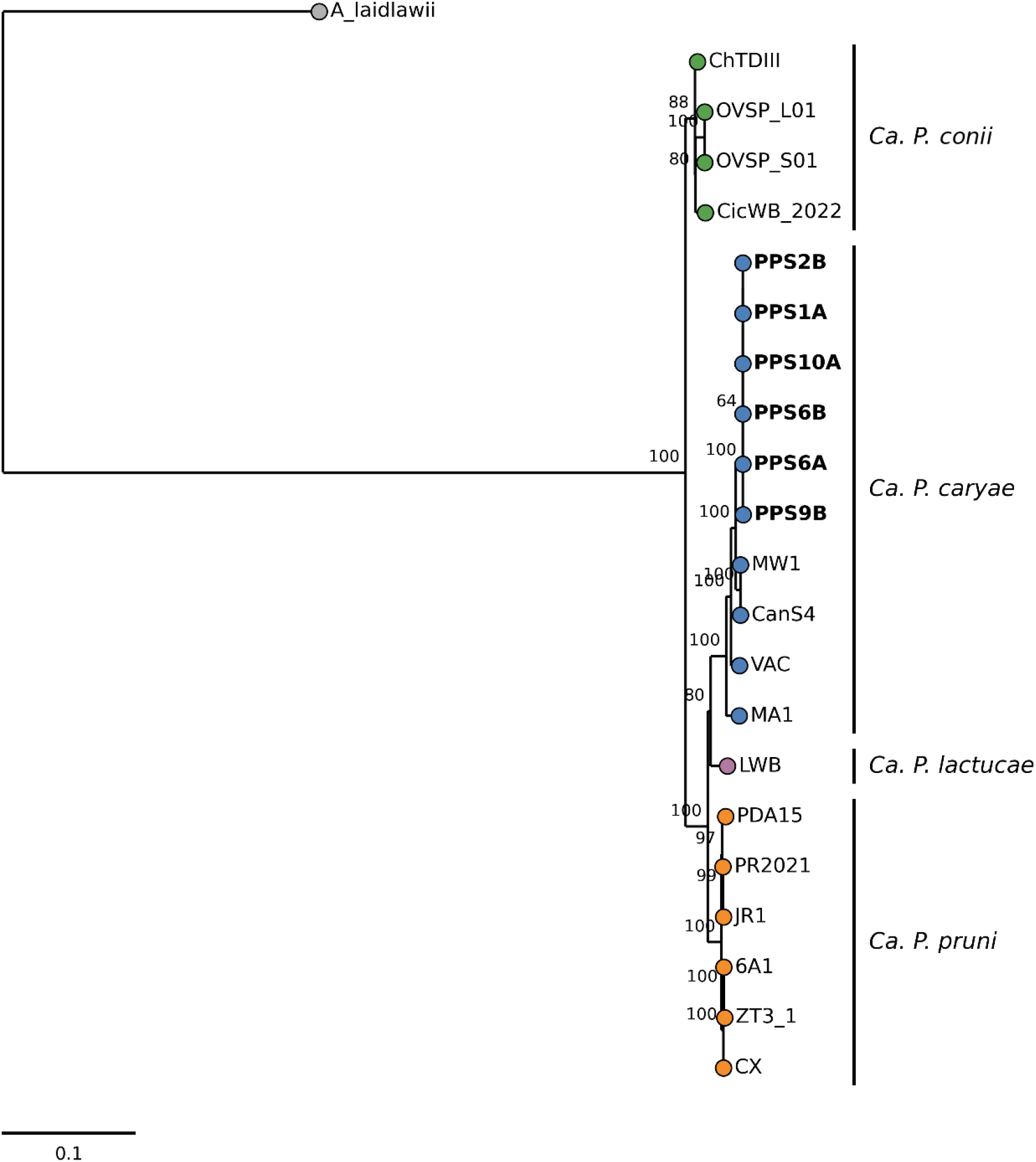
Maximum-likelihood phylogeny of the core genome of 21 *Ca.* P. pruni and *Ca.* P. pruni-related genomes, inferred from the Panaroo core gene alignment with IQ-TREE2 v2.4.0 and 1,000 ultrafast bootstrap replicates, and rooted on *A. laidlawii* PG-8A. Bars indicate the four species, *Ca.* P. caryae, *Ca.* P. pruni, *Ca.* P. conii and *Ca.* P. lactucae; the six PPS strains are shown in bold.

### Comparative genomics

The summary of the inferred values of the pangenome evolution and recombination analyses for the three species *Ca.* P. caryae, *Ca.* P. pruni, and *Ca.* P. conii are shown in Table 4. The Heap’s Law alpha diversity value shows that all three species have open pangenomes (Table 4, Figs. 6, S5.1-S5.3). However, recombination has more impact compared to mutation in *Ca.* P. pruni than the other species (Table 4, Figs. 6; S5.1-S5.6). Although these phytoplasmas have different hosts and ecologies, recombination may have taken place in their vectors and other intermediate pathways [68, 69]. Analyses of the PBT strains show that the inferred recombination can be influenced by the number of genomes compared, since this impact on the number of core genes and the quality of genome sequencing. When the four strains with higher genome completeness were compared, the PBT strains have no detected recombination events in their core genome, indicating that they are clonal strains (Fig. S5.4). However, inclusion of the PPS9B and PPS10A with lower genome completeness distorts the inferred recombination events (Fig. S5.5). Similar conclusions are made from *Ca.* P. pruni, and *Ca.* P. conii strains (Figs. S5.5-S5.6). Interpretation of recombination events should therefore be made with caution, especially for phytoplasma with differing hosts and vectors.

**Figure 6.**
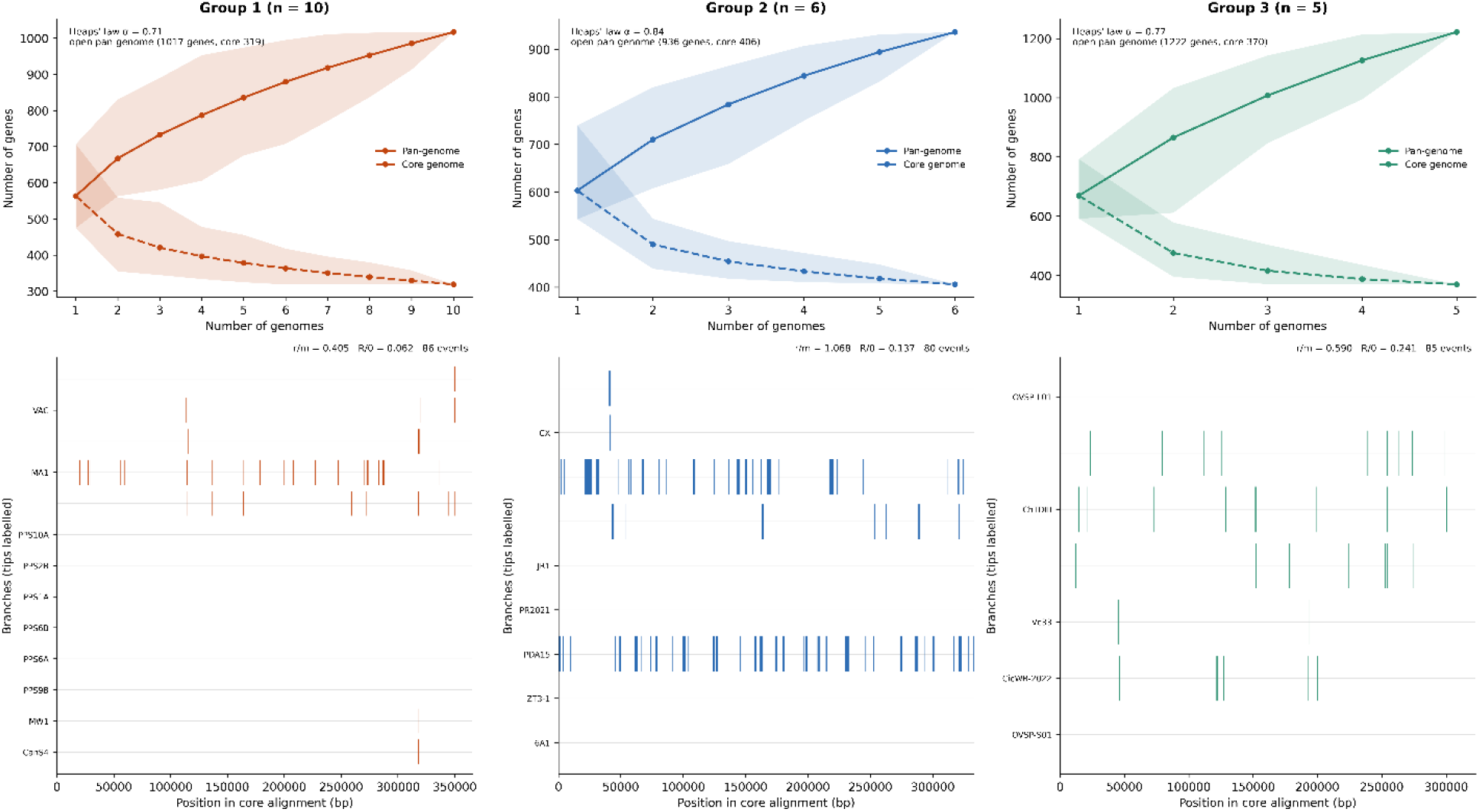
Pangenome and recombination in *Ca.* P. caryae (Group 1, n = 10), *Ca.* P. pruni (Group 2, n = 6) and *Ca.* P. conii (Group 3, n = 5). Top, pangenome (solid line) and core genome (dashed line) accumulation curves, with ranges shaded, Heaps’ law α and pangenome and core genome sizes. Bottom, recombination events inferred with ClonalFrameML along the core genome alignment for each branch, with r/m and R/θ. Values are summarized in Table 4.

### Potential mobile unit

Potential Mobile Units (PMUs) are composite, transposon-like mobile genetic elements that constitute a major driver of genome plasticity, dynamic genome size variation, and host-adaptation in phytoplasmas [70, 71]. A genomic window (or region) containing all five PMU genes (*sigF*, *ssb*, *hupB*, *ftsH* and *smc*) on one contig was found in 12 of the 22 genomes: five of the 10 genomes of group 1; three of six genomes of group 2, three of five genomes of group 3 and the group 4 strain, LWB (Figure 7; Tables S6.1–S6.5). In group 1, genomic windows with PMU genes were found in PPS1A, PPS2B, PPS6A and PPS6B (6,200 to 6,201 bp) and in CanS4 (6,240 bp). In PPS1A and PPS2B strains, the gene order was *smc*-*ftsH*-*hupB*-*ssb*-*sigF* on the reverse strand. In PPS6A and PPS6B it was *sigF*-*ssb*-*hupB*-*ftsH*-*smc* on the forward strand, an inverted but otherwise conserved arrangement. Diversity in trends in gene order and strand orientation are found in genomes of other strains (Figure 7). In PR2021, the window spans 37,435 bp and sigF is represented only by a 36 bp TBLASTN hit without an annotated coding sequence (Tables S6.2, S6.5).

**Figure 7.**
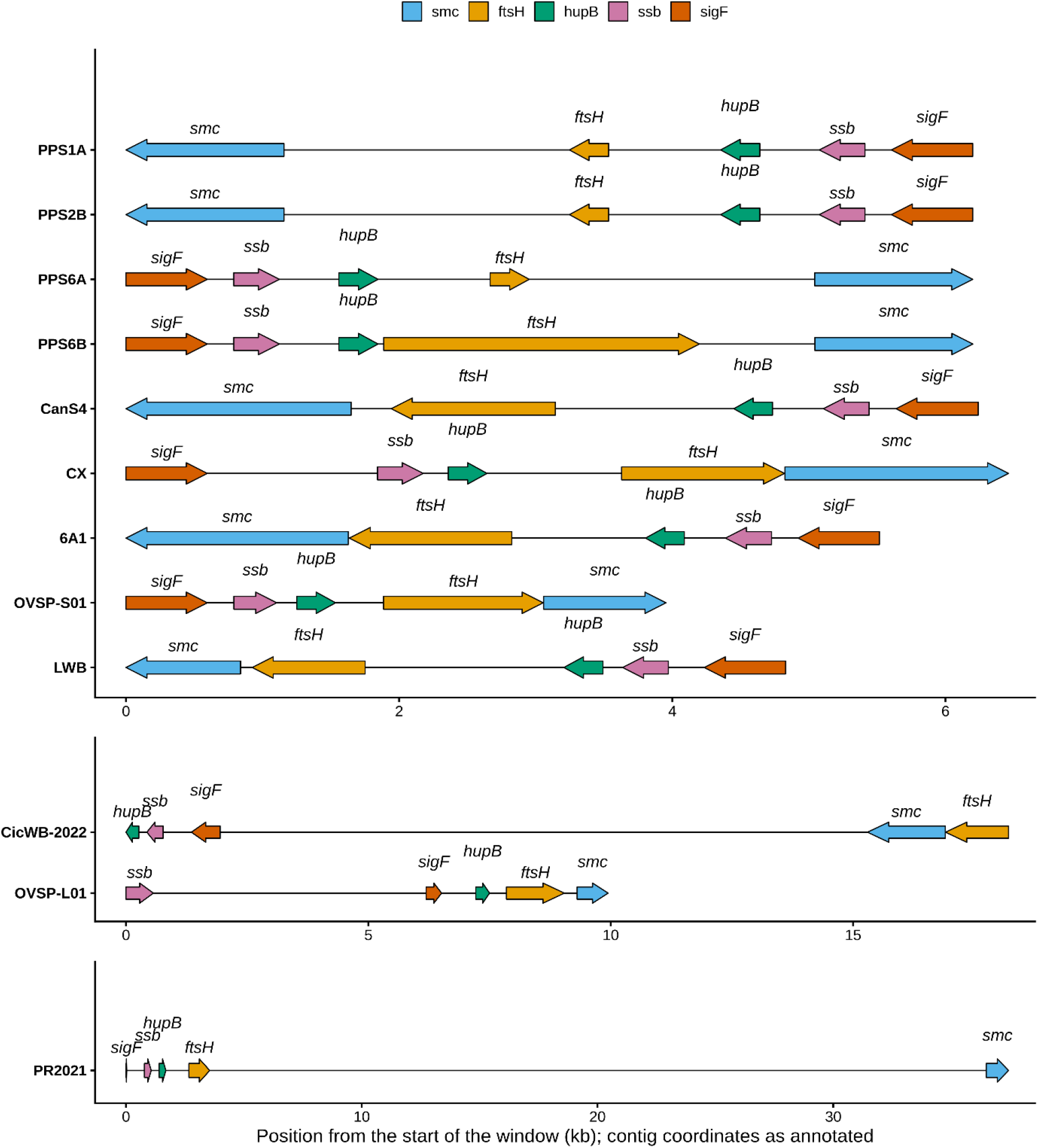
Gene order and coding strand orientation of the five potential mobile unit (PMU) genes (*sigF*, *ssb*, *hupB*, *ftsH* and *smc*) in the 12 genomes with all five genes on one contig: PPS1A, PPS2B, PPS6A, PPS6B and CanS4 (*Ca.* P. caryae); CX, 6A1 and PR2021 (*Ca.* P. pruni); CicWB-2022, OVSP-L01 and OVSP-S01 (*Ca.* P. conii); and LWB (*Ca.* P. lactucae). Arrows show gene length and direction; positions are measured from the start of each window. Panels use different scales to accommodate windows of about 4 to 37 kb (Tables S6.1 and S6.2).

### Metabolic pathways

Metabolic pathway gene content was assessed using 66 curated genes for seven curated pathways. A heatmap of the gene content presence and absence across the nucleotide salvage, glycolysis, folate biosynthesis, glycerophospholipid, DNA replication, protein secretion and transport pathways are shown in Figure 8 and Tables S7.1-S7.11. In general, unless when absent in the four other PBT genomes, gene absences in strains PPS9B and PPS10A were interpreted with caution due to lower genome completeness compared to the other PBT strains. For example, *pyrG*, *glyA*, and *dnaA* were not detected in PPS9B while *dut*, *pgi*, *tpiA*, *gapA*, *gpmI*, *gpsA*, *dnaN*, *secY*, *potD*, and *dnaA* were not detected in PPS10A. None of these absences were shared with the four complete strains, and these may be attributed to the lower qualities of PPS9B and PPS10A assemblies. This same principle was also applied to genomes of OVSP-L01, OVSP-S01, Vc33.

**Figure 8.**
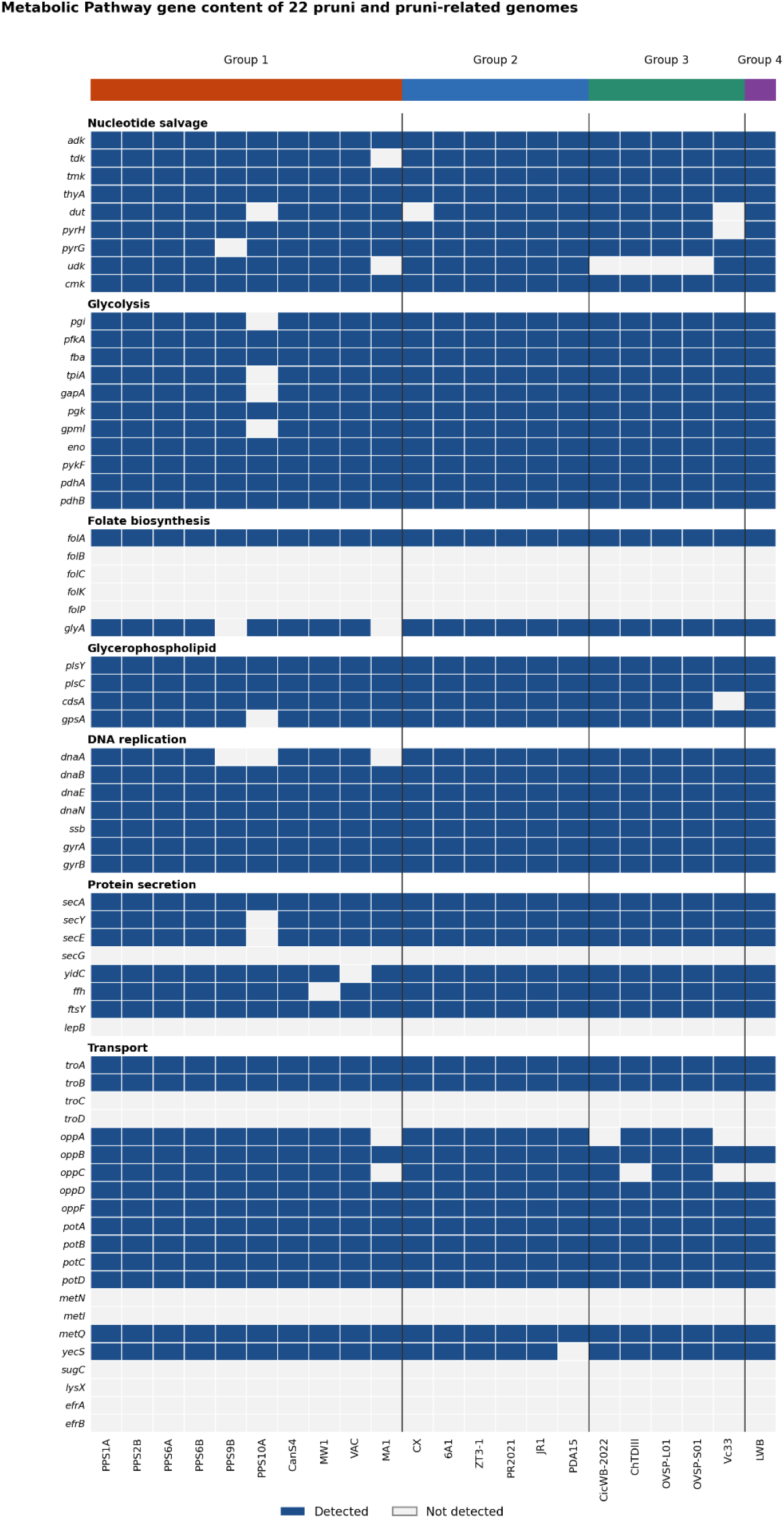
Presence and absence of 66 metabolic genes in seven pathways (nucleotide salvage, glycolysis, folate biosynthesis, glycerophospholipid biosynthesis, DNA replication, protein secretion and transport) in the 22 genomes. Genes were detected by Prokka annotation or BLASTp (Table S7.2). Dark blue, detected; grey, not detected. The colour bar above the columns indicates the ANI group.

Across the 22 genomes, 32 of the 66 genes were present in every genome, 14 genes (*folB*, *folC*, *folK*, *folP*, *secG*, *lepB*, *troC*, *troD*, *metN*, *metI*, *sugC*, *lysX*, *efrA* and *efrB*) were absent from every genome, and 20 varied among genomes. No gene was specific to any of the four species nor missing from any given species. However, in strain LWB which is the only strain in *Candidatus* Phytoplasma lactucae, genes for OppA and OppC which are core components of the Oligopeptide Permease (Opp) system [72], were absent (*oppA* is also missing in strains MA1 and CicWB-2022) (Figure 8, Tables S7.2-S7.3; S7.5-S7.6).

A complete nucleotide salvage pathway gene set is present in PPS1A, PPS2B, PPS6A, PPS6B, CanS4, MW1 and VAC of *Ca.* P. caryae (9/9 genes). This gene set is also present in all strains of *Ca.* P. pruni (except strain CX) and *Ca.* P. lactucae. However, multiple strains of *Ca.* P. conii lack *udk* gene, which encodes cytidine/uridine kinase, a catalyst within the pyrimidine salvage pathway. Loss of *udk* genes have also been similarly documented in ‘*Candidatus* Phytoplasma mali’ (16SrX group) and ‘*Candidatus* Phytoplasma australiense’ (16SrXII group), suggesting that this is not unique to *Ca.* P. conii strains [73]. Genes for glycolysis, glycerophospholipid biosynthesis and DNA replication were present in majority of the genomes (Fig. 8, Tables S7.2, S7.3 and S7.5). Folate biosynthesis showed the most pronounced reduction, with only 2 (*folA* and *glyA*) of 6 genes detected across genomes (Fig. 8, Tables S7.2, S7.3 and S7.5). Phytoplasma genomes are known to selectively retain downstream salvage and reduction enzymes while surviving as metabolic parasites that depend strictly on scavenging preformed folates from their environments including plant phloem sap, vector hemolymph and salivary glands and transport systems [62, 70, 74].

### Secretome prediction and effector candidates

SignalP v5.0b [66] predicted between 11 and 25 secreted proteins per strain, representing 2.5% to 4.6% of total coding sequences (Table S8.1; Figure S6.2). Across all 22 genomes, SignalP predicted 11 (PPS10A) to 37 (CicWB-2022, Vc33 and ChTDIII) secreted proteins per genome (Tables S8.1 and S8.3). Most predicted secreted proteins were annotated as hypothetical proteins. TroA (periplasmic zinc-binding protein) and DppB (dipeptide transport system permease) were detected across all six strains. YecS was present among the predicted secreted proteins of PPS1A, PPS2B, PPS6A, and PPS6B. The total predicted secretome across all six strains comprised 127 candidates, all but one of which (PPS10A_00365) had a homologue in at least one of the other 16 genomes (Tables S9.1–S9.5). The effectors were further compared using a curated set of effectors, including TENGU, SAP05, SAP06/48-like, SAP11, SAP54, SAP54-like, SVM_705, SVM_1019, SVM_1100, SVM_1102 and SVM_1103 [49]. The variation in effector gene content in all 22 strains is shown in Table S10.1. Among these candidates, a homolog of the AY-WB effector SAP11 (93.4% amino acid identity; PPS1A_00098, PPS2B_00459, PPS6A_00114, and PPS6B_00526) was identified in PPS1A, PPS2B, PPS6A, and PPS6B, and a fragment of SAP54 (77.5% identity over 32% of the query) was found in PPS1A, PPS2B, PPS6A and PPS6B (Table S10.5).

### Conclusions

This study provides first genome-based confirmation of pecan bunchy top phytoplasma and a comprehensive genome based analyses of strains known as ‘*Candidatus* Phytoplasma pruni’ and ‘*Candidatus* Phytoplasma pruni’-related strains. None of the reclassified ‘*Candidatus* Phytoplasma pruni’-related strains have a *Pruni* species as a host, implying different and diverse ecologies. These genome comparisons led to the recommendation for the reclassification of these strains into four *Candidatus* species, including two novel species. The description for each species is provided below.

**Updates on the description of ‘*Candidatus* Phytoplasma pruni’ and description of three novel *Candidatus* Phytoplasma Species ‘*Candidatus* Phytoplasma caryae’, ‘*Candidatus* Phytoplasma conii’** and **‘*Candidatus* Phytoplasma lactucae’.**

**‘*Candidatus* Phytoplasma pruni’**: Species description is as provided previously [11]. The genome of the previously designated reference strain, PX11CT1R, has not been sequenced. Strains that cluster under this epithet include CX, 6A1, ZT3-1, PR2021, JR1 and PDA15 (GenBank Accession numbers are provided in Tables 1 and S1.2). Multilocus sequence analysis of ‘*Ca.* P. pruni’ strains from *Prunus* hosts in North America resolved five lineages on commercial *Prunus* species and two on wild *Prunus virginiana*, each with an average 16S rRNA gene identity of 99.75% or more to strain CX; the CX and ZT3 genomes represent the Eastern-X and Northwestern-X2 lineages, respectively [75]. The vector includes various leafhopper species.

[(Mollicutes) NC; NA; O, wall-less; NAS (16S rRNA GenBank accession number for PX11CT1R: JQ044393); P (*Prunus,* phloem)]; ex-Davis *et al. [11]*; Rabiu *et al.* this study. Unique regions of the 16S rRNA gene compared to strains of all three other species are: C197T, C830T, C1264A. Other variations in the 16S rRNA are provided in Figs. S2.1–S2.6, Table 3 and Table S2.3. Unique nucleotide sequences *rpoB*, *rps19*, *rps3*, *rpl22*, *secY*, and *secA* are shown in Figure S2.8 and Table S3.

**‘*Candidatus* Phytoplasma caryae’** [ca.ry’ae. N.L. gen. fem. n. caryae, of *Carya*, a botanical genus name, referring to *Carya illinoinensis* (pecan), the natural plant host of the reference strain]. Since pecan bunchy top phytoplasma is one of the oldest described phytoplasma and falls into this *Candidatus* species, strain PPS6B with highest coverage of 76x and completeness of 97.26% is the designated reference strain for this taxon. Other strains in this taxon include other pecan bunchy top phytoplasma strains PPS1A, PPS2B, PPS6A, PPS9B, PPS10A; *Milkweed yellows phytoplasma* (MYp-CanS4 or simply CanS4, MW1)), *Vaccinium* witches’ broom phytoplasma (VAC)), *Italian clover phyllody* (MA1). GenBank accession numbers are provided in Tables 1, S1.1-S1.2. Leafhopper vectors are only confirmed for *Vaccinium* witches’ broom and *Italian clover phyllody* phytoplasmas.

[(Mollicutes) NC; NA; O, wall-less; NAS (Genome GenBank accession number for PPS6B: JCFZOO000000000 (PPS6B); 16S rRNA GenBank accession number for PPS6B: QB316901); P (*Carya illinoinensis*, phloem)]; Rabiu *et al.*; this study. Unique regions of the 16S rRNA gene compared to strains of all three other *Candidatus* species are: A171G, C1332T and G1346A. This nucleotide change is conserved in other strains of the species except strain MA1. Other variations in the 16S rRNA gene compared to the three other *Candidatus* species are provided in Figs. S2.1–S2.6, Table 3 and Table S2.3. Unique nucleotide sequences *rpoB*, *rps19*, *rps3*, *rpl22*, *secY*, and *secA* are shown in Figure S2.8 and Table S3.

**‘*Candidatus* Phytoplasma conii’** [co’ni.i. N.L. gen. neut. n. conii, of *Conium*, a botanical genus name, referring to *Conium maculatum* (poison hemlock), the natural plant host of the reference strain]. The reference strain for this taxon is CicWB-2022. Other strains in this taxon based on ANI, dDDH and phylogenetic reconstruction include OVSP-L01, X-disease group phytoplasma of *Catharanthus roseus* (Vc33), *China tree decline phytoplasma* (ChTDIII), and OVSP-S01. Various leafhopper vectors have been reported as vectors for members of this taxon. The nomenclatures ‘*Candidatus* Phytoplasma platensis’ [80] and ‘*Candidatus* Phytoplasma manihotii’ [17] were proposed earlier in preprints and are not validly published; ‘*Candidatus* Phytoplasma conii’ maintains the host-based naming system of this taxon. The Genbank accession numbers are present in Tables 1 and S1.2.

[(Mollicutes) NC; NA; O, wall-less; NAS (Genome GenBank accession number for CicWB-2022: GCA_035853675.1); P (*Conium maculatum*, phloem)]; Rabiu *et al.*; this study. Unique regions of the 16S rRNA gene compared to strains of all three other *Candidatus* species are: A647T. Other variations in the 16S rRNA gene compared to the three other *Candidatus* species are provided in Figs. S2.1–S2.6, Table 3 and Table S2.3. Unique nucleotide sequences *rpoB*, *rps19*, *rps3*, *rpl22*, *secY*, and *secA* are shown in Figure S2.8 and Table S3.

**‘*Candidatus* Phytoplasma lactucae’** [lac.tu’cae. L. gen. fem. n. *lactucae*, of *Lactuca*, a botanical genus name, referring to *Lactuca sativa* (lettuce), the plant host from which the reference strain was characterized]. This taxon has only one strain, LWB, causal agent of Lettuce witches’ broom and is the reference strain. Various leafhopper vectors have been reported as vectors for members of this taxon. The GenBank accession number is GCA_051397825.1 (Table 2).

**Table 2.** Genomes of other *Ca.* P. pruni and *Ca.* P. pruni-related strains used in this study.

| Phytoplasma | Strain | 16SrIII group | Host | Year of isolation | Country of isolation | Accession | Reference |
|---|---|---|---|---|---|---|---|
| *Candidatus* Phytoplasma pruni | JR1 | 16SrIII-A | *Catharanthus roseus* (periwinkle)<sup>a</sup> | NA | NA | GCF_000309465.1 | [76] |
| *Candidatus* Phytoplasma pruni | PR2021 | 16SrIII-A | *Euphorbia pulcherrima* (poinsettia) | 2021 | Taiwan (Taipei) | GCA_029746895.1 | [3] |
| *Candidatus* Phytoplasma pruni | CX | 16SrIII-A | *Prunus persica* (peach) | 1991 | Canada (eastern Canada) | GCF_001277135.1 | [12] |
| *Candidatus* Phytoplasma pruni | 6A1 (2A1) | 16SrIII-A | *Syringa vulgaris* (lilac) | 2020 | Canada (Victoria) | GCA_033391615.1 | [15] |
| China tree decline | ChTDIII | 16SrIII-B | *Melia azedarach* (China tree) | 2019 | Argentina | GCF_013391955.1 | [13] |
| Italian clover phyllody | MA1 | 16SrIII-B | *Catharanthus roseus* (periwinkle)<sup>a</sup> | NA | NA | GCF_000300695.1 | [76] |
| Milkweed yellows phytoplasma | MW1 | 16SrIII-F | *Catharanthus roseus* (periwinkle)<sup>a</sup> | NA | NA | GCF_000309485.1 | [76] |
| Milkweed yellows phytoplasma | MYp-CanS4 | 16SrIII-F | *Asclepias syriaca* (common milkweed) | 2024 | Canada (Ontario) | GCA_050286905.1 | [15] |
| Vaccinium witches’ broom | VAC | 16SrIII-F | *Catharanthus roseus* (periwinkle)<sup>a</sup> | NA | NA | GCF_000309405.1 | [76] |
| Cicuta witches’ broom | CicWB-2022 | 16SrIII-J | *Conium maculatum* (poison hemlock) | 2022 | Argentina (Villa Dolores, Córdoba) | GCA_035853675.1 | [16] |
| Phytoplasma Vc33 | Vc33 | 16SrIII-J | NA | NA | NA | GCA_001623385.2 | [77] |
| Lettuce witches’ broom phytoplasma | LWB | 16SrIII-X | *Lactuca sativa* (lettuce) | 2023 | Argentina (Córdoba) | GCA_051397825.1 | [14] |
| *Candidatus* Phytoplasma pruni | ZT3-1 | 16SrIII | *Prunus avium* (sweet cherry) | 2019 | USA (Zillah, Washington) | GCA_024586395.1 | [78] |
| *Candidatus* Phytoplasma pruni | PDA15 | 16SrIII | *Vitis vinifera* (grapevine) | 2016 | USA (Pennsylvania) | GCA_041984945.1 | NA |
| *Phytoplasma* sp. | OVSP-L01 | 16SrIII-B | *Manihot esculenta* (cassava) | 2023 | Brazil (Salitre, Ceará) | GCA_059597165.1 | [17] |
| *Phytoplasma* sp. | OVSP-S01 | 16SrIII-B | *Manihot esculenta* (cassava) | 2023 | Brazil (Salitre, Ceará) | GCA_059597155.1 | [17] |

NA, not available in the NCBI record or cited reference; to be completed. Host, year and location are as recorded in the NCBI BioSample. <sup>a</sup> The BioSample lists periwinkle, in which the strain is maintained; the original host, year and country are to be added from the cited reference.

[(Mollicutes) NC; NA; O, wall-less; NAS (Genome GenBank accession number for LWB: GCA_051397825.1; 16S rRNA GenBank accession numbers); P (*Lactuca sativa*, phloem)]; ex-Fernandez *et al. [14]*; Rabiu *et al.*; this study. Unique regions of the 16S rRNA gene compared to strains of all three other *Candidatus* species are: T182C, C412T, C413T and G617A (numbering of the PPS1A 16S rRNA gene). The published 16SrIII-X sequences KC412026 and MH497013 carry the same bases at these positions. Other variations in the 16S rRNA gene compared to the three other *Candidatus* species are provided in Figs. S2.1–S2.6, Table 3 and Table S2.3. Unique nucleotide sequences *rpoB*, *rps19*, *rps3*, *rpl22*, *secY*, and *secA* are shown in Figure S2.8 and Table S3.

**Table 3.**
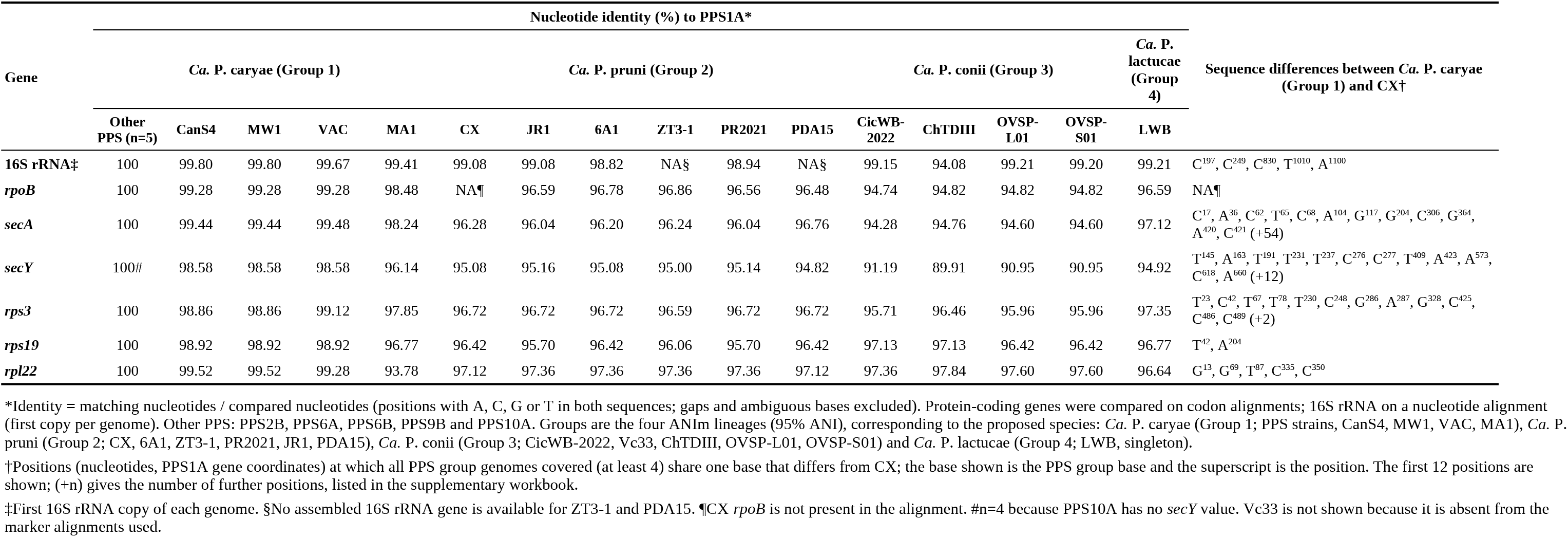
Nucleotide identity and PPS group-specific sequence differences in PPS1A, the reference strain of the PPS group, in comparison with all other pruni and pruni-related genomes.

| Gene | Nucleotide identity (%) to PPS1A* |  |  |  |  |  |  |  |  |  |  |  |  |  |  |  | Sequence differences between *Ca.* P. caryae (Group 1) and CX† |
|---|---|---|---|---|---|---|---|---|---|---|---|---|---|---|---|---|---|
|  | *Ca.* P. caryae (Group 1) |  |  |  |  | *Ca.* P. pruni (Group 2) |  |  |  |  |  | *Ca.* P. conii (Group 3) |  |  |  | *Ca.* P. lactucae (Group 4) |  |
|  | Other PPS (n=5) | CanS4 | MW1 | VAC | MA1 | CX | JR1 | 6A1 | ZT3-1 | PR2021 | PDA15 | CicWB-2022 | ChTDIII | OVSP-L01 | OVSP-S01 | LWB |  |
| **16S rRNA‡** | 100 | 99.80 | 99.80 | 99.67 | 99.41 | 99.08 | 99.08 | 98.82 | NA§ | 98.94 | NA§ | 99.15 | 94.08 | 99.21 | 99.20 | 99.21 | C<sup>197</sup>, C<sup>249</sup>, C<sup>830</sup>, T<sup>1010</sup>, A<sup>1100</sup> |
| ***rpoB*** | 100 | 99.28 | 99.28 | 99.28 | 98.48 | NA¶ | 96.59 | 96.78 | 96.86 | 96.56 | 96.48 | 94.74 | 94.82 | 94.82 | 94.82 | 96.59 | NA¶ |
| ***secA*** | 100 | 99.44 | 99.44 | 99.48 | 98.24 | 96.28 | 96.04 | 96.20 | 96.24 | 96.04 | 96.76 | 94.28 | 94.76 | 94.60 | 94.60 | 97.12 | C<sup>17</sup>, A<sup>36</sup>, C<sup>62</sup>, T<sup>65</sup>, C<sup>68</sup>, A<sup>104</sup>, G<sup>117</sup>, G<sup>204</sup>, C<sup>306</sup>, G<sup>364</sup>, A<sup>420</sup>, C<sup>421</sup> (+54) |
| ***secY*** | 100# | 98.58 | 98.58 | 98.58 | 96.14 | 95.08 | 95.16 | 95.08 | 95.00 | 95.14 | 94.82 | 91.19 | 89.91 | 90.95 | 90.95 | 94.92 | T<sup>145</sup>, A<sup>163</sup>, T<sup>191</sup>, T<sup>231</sup>, T<sup>237</sup>, C<sup>276</sup>, C<sup>277</sup>, T<sup>409</sup>, A<sup>423</sup>, A<sup>573</sup>, C<sup>618</sup>, A<sup>660</sup> (+12) |
| ***rps3*** | 100 | 98.86 | 98.86 | 99.12 | 97.85 | 96.72 | 96.72 | 96.72 | 96.59 | 96.72 | 96.72 | 95.71 | 96.46 | 95.96 | 95.96 | 97.35 | T<sup>23</sup>, C<sup>42</sup>, T<sup>67</sup>, T<sup>78</sup>, T<sup>230</sup>, C<sup>248</sup>, G<sup>286</sup>, A<sup>287</sup>, G<sup>328</sup>, C<sup>425</sup>, C<sup>486</sup>, C<sup>489</sup> (+2) |
| ***rps19*** | 100 | 98.92 | 98.92 | 98.92 | 96.77 | 96.42 | 95.70 | 96.42 | 96.06 | 95.70 | 96.42 | 97.13 | 97.13 | 96.42 | 96.42 | 96.77 | T<sup>42</sup>, A<sup>204</sup> |
| ***rpl22*** | 100 | 99.52 | 99.52 | 99.28 | 93.78 | 97.12 | 97.36 | 97.36 | 97.36 | 97.36 | 97.12 | 97.36 | 97.84 | 97.60 | 97.60 | 96.64 | G<sup>13</sup>, G<sup>69</sup>, T<sup>87</sup>, C<sup>335</sup>, C<sup>350</sup> |

\*Identity = matching nucleotides / compared nucleotides (positions with A, C, G or T in both sequences; gaps and ambiguous bases excluded). Protein-coding genes were compared on codon alignments; 16S rRNA on a nucleotide alignment (first copy per genome). Other PPS: PPS2B, PPS6A, PPS6B, PPS9B and PPS10A. Groups are the four ANIm lineages (95% ANI), corresponding to the proposed species: *Ca.* P. caryae (Group 1; PPS strains, CanS4, MW1, VAC, MA1), *Ca.* P. pruni (Group 2; CX, 6A1, ZT3-1, PR2021, JR1, PDA15), *Ca.* P. conii (Group 3; CicWB-2022, Vc33, ChTDIII, OVSP-L01, OVSP-S01) and *Ca.* P. lactucae (Group 4; LWB, singleton).
†Positions (nucleotides, PPS1A gene coordinates) at which all PPS group genomes covered (at least 4) share one base that differs from CX; the base shown is the PPS group base and the superscript is the position. The first 12 positions are shown; (+n) gives the number of further positions, listed in the supplementary workbook.
‡First 16S rRNA copy of each genome. §No assembled 16S rRNA gene is available for ZT3-1 and PDA15. ¶CX *rpoB* is not present in the alignment. #n=4 because PPS10A has no *secY* value. Vc33 is not shown because it is absent from the marker alignments used.

**Table 4.** Average nucleotide identity (ANI), pangenome and recombination summary for the four ANI groups (*Ca.* P. caryae, *Ca.* P. pruni, *Ca.* P. conii and *Ca.* P. lactucae).

| ANI group<sup>*</sup> | *Ca*. P. caryae (Group 1, n = 10) | *Ca*. P. pruni (Group 2, n = 6) | *Ca*. P. conii (Group 3, n = 5) | *Ca*. P. lactucae (Group 4, n = 1) |
|---|---|---|---|---|
| *Ca*. P. caryae (Group 1, n = 10) | 96.06–99.96 | 93.91–95.47 | 91.42–93.39 | 93.59–94.78 |
| *Ca*. P. pruni (Group 2, n = 6) | 93.91–95.47 | 97.25–99.71 | 91.89–93.09 | 93.87–94.85 |
| *Ca*. P. conii (Group 3, n = 5) | 91.42–93.39 | 91.89–93.09 | 95.99–99.47 | 92.81–93.27 |
| *Ca*. P. lactucae (Group 4, n = 1) | 93.59–94.78 | 93.87–94.85 | 92.81–93.27 | NA<sup>†</sup> |
| Pangenome comparison |  |  |  |  |
| ANI group | Number of genes in pangenome | Number of genes in core genome | Heaps’ law α<sup>‡</sup> |  |
| *Ca*. P. caryae (Group 1, n = 10) | 1017 | 319 | 0.71 |  |
| *Ca*. P. pruni (Group 2, n = 6) | 936 | 406 | 0.84 |  |
| *Ca*. P. conii (Group 3, n = 5) | 1222 | 370 | 0.77 |  |
| *Ca*. P. lactucae (Group 4, n = 1) | NA<sup>†</sup> | NA<sup>†</sup> | NA<sup>†</sup> |  |
| Recombination |  |  |  |  |
| ANI group | R/θ<sup>§</sup> | r/m<sup>§</sup> | Number of recombination events |  |
| *Ca*. P. caryae (Group 1, n = 10) | 0.062 | 0.405 | 86 |  |
| *Ca*. P. pruni (Group 2, n = 6) | 0.137 | 1.068 | 80 |  |
| *Ca*. P. conii (Group 3, n = 5) | 0.241 | 0.590 | 85 |  |
| *Ca*. P. lactucae (Group 4, n = 1) | NA<sup>†</sup> | NA<sup>†</sup> | NA<sup>†</sup> |  |

<sup>\*</sup>Values are FastANI average nucleotide identity (%), the mean of both directions, given as the lowest and highest value among pairs of different genomes within a group (diagonal) and between two groups.
<sup>†</sup>NA, not applicable; *Ca*. P. lactucae is represented by a single genome.
<sup>‡</sup>Heaps’ law α < 1 indicates an open pangenome.
<sup>§</sup>R/θ, ratio of recombination to mutation rates; r/m, ratio of the effects of recombination to mutation on nucleotide diversity (ClonalFrameML).

## Supporting information

Rabiu et al., Supplementary figures and Tables

## Acknowledgements

This study was funded in part by Oklahoma State University Agricultural Experimental Station HATCH grant to Dr. Mustafa O. Jibrin. A portion of the computing for this project was performed at the High Performance Computing Center at Oklahoma State University supported in part through the National Science Foundation grant OAC-1531128. We also thank members of Oklahoma Pecan Growers Association for discussions at pecan workshops that led to this project and the grower who guided us in sampling his pecan orchard.

## Author contributions

Conceptualization, funding acquisition and study design: MOJ; Survey and sample collection: URB and MOJ; qPCR, Sanger Sequencing and interpretation: SW and MOJ; Genome sequencing, assembly and analyses: URB, CLM, MOJ, paper draft: URB and MOJ; Supervision: MOJ, NRW, SM, FOC. All authors approved the final version of the manuscript.

## Author Notes on Data Availability

The GenBank accession numbers of the 16S rRNA genes for strains PPS1A, PPS2B, PPS6A, PPS6B, PPS9B and PPS10A are QB316898, QB316899, QB316900, QB316901, QB316902, QB316903 respectively. The genomes of the six pecan bunchy top phytoplasma (PBT) strains in this study have been deposited in the NCBI database under the BioProject accession number PRJNA1541623. The whole genome assemblies have been deposited in GenBank under the accession numbers JCFZOL000000000 (PPS1A), JCFZOM000000000 (PPS2B), JCFZON000000000 (PPS6A), JCFZOO000000000 (PPS6B), JCFZOP000000000 (PPS9B) and JCFZOQ000000000 (PPS10A). Specimen of pecan leaves and petioles from which the six strains were derived have been deposited in the Oklahoma State University Herbarium (OKLA). OKLA is part of the Texas-Oklahoma Regional Consortium of Herbaria (TORCH). The curator at the time when the samples were deposited is Dr. Mark Fishbein.

## List of Supplementary Figures

Supplementary Figure S1.1. Schematic of the sampled orchard (>500 trees) showing the distribution of symptomatic trees in one section of the orchard displaying typical pecan bunch top symptoms. Six symptomatic pecan trees were selected for sampling.

Supplementary Figure S1.2. iPhyClassifier [5] virtual RFLP analysis confirming 16Sr group and subgroup classification for all six *Ca.* Phytoplasma pruni (16SrIII-C) PPS strains. All six PPS strains were classified as subgroup 16SrIII-C (input: full-length 1,523 bp 16S rRNA gene).

Supplementary Figure S1.3. Genome quality metrics for the six *Ca.* Phytoplasma pruni (16SrIII-C) PPS strains assessed by CheckM v1.2.5. (A) Genome completeness (%) per strain; dashed reference lines at 90% and 95%. (B) Mean sequencing depth (x) per strain. (C) Genome completeness versus mean sequencing depth; bubble size proportional to genome size (bp).

Supplementary Figure S1.4. Whole-genome synteny dot plots of the six *Ca.* P. pruni PPS strains (PPS1A, PPS2B, PPS6A, PPS6B, PPS9B and PPS10A; y-axis, Mb) against the genome of strain CanS4 (x-axis, Mb). Blue lines are forward-strand alignments and red lines are reverse-strand alignments. A continuous diagonal indicates collinear sequence. Left, all six strains; right, PPS1A, PPS2B, PPS6A and PPS6B enlarged.

Supplementary Figure S1.5. Whole-genome synteny dot plots of the six PPS strains against the genome of strain CicWB-2022.

Supplementary Figure S1.6. Whole-genome synteny dot plots of the six PPS strains against the genome of strain CX.

Supplementary Figure S1.7. Whole-genome synteny dot plots of the six PPS strains against the genome of strain LWB.

Supplementary Figure S2.1. Pairwise 16S rRNA gene identity (%) among the 16S rRNA gene copies of the PPS strains and the *Ca.* P. pruni and *Ca.* P. pruni-related reference strains. The suffixes c1 and c2 label the 16S rRNA gene copies of a genome. Sequence labels are colored by group (Group 1, green; Group 2, orange; Group 3, purple; Group 4, pink). Identity ranges from 92% to 100%.

Supplementary Figure S2.2. Group-specific nucleotide positions in the 16S rRNA gene. The matrix shows the base at each of 11 positions (171, 1332, 1346, 197, 830, 1264, 647, 182, 412, 413 and 617) in every 16S rRNA gene copy. The bar above the matrix marks the group that each position distinguishes (Group 1, positions 171, 1332 and 1346; Group 2, positions 197, 830 and 1264; Group 3, position 647; Group 4, positions 182, 412, 413 and 617). Black outlines mark the bases that define the group. MA1 does not carry the Group 1 bases at positions 171, 1332 and 1346, and VAC carries T at position 1264.

Supplementary Figure S2.3. Sequence context of the group-specific 16S rRNA gene positions 171, 1332 and 1346 (Group 1). Each panel shows the aligned sequence around one position, with the position boxed and numbered below the panel. Bases are colored: A, green; C, blue; G, orange; T, red.

Supplementary Figure S2.4. Sequence context of the group-specific 16S rRNA gene positions 197, 830 and 1264 (Group 2). Each panel shows the aligned sequence around one position, with the position boxed and numbered below the panel. Bases are colored: A, green; C, blue; G, orange; T, red.

Supplementary Figure S2.5. Sequence context of the group-specific 16S rRNA gene position 647 (Group 3). The aligned sequence around the position is shown, with the position boxed and numbered below the panel. Bases are colored: A, green; C, blue; G, orange; T, red.

Supplementary Figure S2.6. Sequence context of the group-specific 16S rRNA gene positions 182, 412, 413 and 617 (Group 4). Each panel shows the aligned sequence around one position, with the position boxed and numbered below the panel. Bases are colored: A, green; C, blue; G, orange; T, red.

Supplementary Figure S2.7. Number of group-specific positions in the 16S rRNA gene for each group: Group 1, 3; Group 2, 3; Group 3, 1; Group 4, 4.

Supplementary Figure S2.8. Codon-level differences from PPS1A in *rpoB* (1,247 codons), *secY* (423 codons), *secA* (836 codons), *rps3* (279 codons) and *rpl22* (139 codons). Each row is one strain and each column is one codon. Codons are identical to PPS1A (light grey), synonymous changes (orange), non-synonymous changes (blue) or gaps or unresolved codons (grey). The first row in each panel combines the PPS strains with no differences to PPS1A, and the number of strains is shown in parentheses. Numbers at the right show the nucleotide differences from PPS1A and the nucleotide identity (%). Strains are ordered from the most to the least similar to PPS1A.

Supplementary Figure S3.1. Pairwise average nucleotide identity (ANI) heatmaps of the 22 *Ca.* P. pruni and *Ca.* P. pruni-related genomes, with *Acholeplasma laidlawii* PG-8A as the outgroup, calculated with pyANI v0.2.12 [48]. (A) ANIb. (B) ANIm.

Supplementary Figure S3.2. Digital DNA-DNA hybridization (dDDH) dendrogram of the 22 genomes, calculated with the Genome-to-Genome Distance Calculator formula 2 (d4) and clustered by UPGMA (average linkage). The dashed line marks the 70% dDDH species threshold. Strain labels are colored by group (Group 1, red; Group 2, blue; Group 3, green; Group 4, purple).

Supplementary Figure S3.3. Pairwise dDDH heatmaps of the 22 genomes calculated with the three Genome-to-Genome Distance Calculator formulas: formula 1 (d0, HSP length over total length), formula 2 (d4, identities over HSP length) and formula 3 (d6, identities over total length). Darker blue indicates higher dDDH. Black outlines mark pairs with dDDH of 70% or more. Strain labels are colored by group.

Supplementary Figure S4.1. Maximum-likelihood tree of the concatenated ribosomal protein genes of the rp operon (20 genes). The tree was generated using IQ-TREE2 v2.4.0 with 1,000 ultrafast bootstrap replicates and is rooted on *Acholeplasma laidlawii*.

Supplementary Figure S4.2. Maximum-likelihood tree of the *rpoB* gene (RNA polymerase beta subunit). The tree was generated using IQ-TREE2 v2.4.0 with 1,000 ultrafast bootstrap replicates and is rooted on *Acholeplasma laidlawii*.

Supplementary Figure S4.3. Maximum-likelihood tree of the *secY* gene (protein translocase subunit). The tree was generated using IQ-TREE2 v2.4.0 with 1,000 ultrafast bootstrap replicates and is rooted on *Acholeplasma laidlawii*.

Supplementary Figure S4.4. Maximum-likelihood tree of the *tuf* gene (elongation factor Tu). The tree was generated using IQ-TREE2 v2.4.0 with 1,000 ultrafast bootstrap replicates and is rooted on *Acholeplasma laidlawii*.

Supplementary Figure S4.5. Maximum-likelihood tree of the concatenated alignment of 23 marker genes (5,425 amino acid sites): *rpoB*, *secA*, *secY* and 20 ribosomal protein genes (*rps10*, *rpl3*, *rpl4*, *rpl2*, *rps19*, *rpl22*, *rps3*, *rpl16*, *rpl29*, *rps17*, *rpl14*, *rpl24*, *rpl5*, *rps14*, *rps8*, *rpl6*, *rpl18*, *rps5*, *rpl30*, *rpl15*). The tree was generated using IQ-TREE2 v2.4.0 with 1,000 ultrafast bootstrap replicates and is rooted on *Acholeplasma laidlawii*.

Supplementary Figure S4.6. Maximum-likelihood tree of the core genome of 84 genomes, inferred from a 3,300-nucleotide core gene alignment (Panaroo) under the GTR+F+I+R4 model. The tree was generated using IQ-TREE2 v2.4.0 with 1,000 ultrafast bootstrap replicates and is rooted on *Acholeplasma laidlawii*.

Supplementary Figure S5.1. Pan-genome and recombination in Group 1 (*Ca.* P. caryae, 10 genomes). Left, number of genes in the pan-genome (solid line) and the core genome (dashed line) as genomes are added; shaded areas show the variation among genome orders. Heaps’ law exponent 0.71; 1,017 pan-genome genes and 319 core genes. Right, recombination events per branch along the core alignment (ClonalFrameML): r/m = 0.405, R/theta = 0.062, 86 events.

Supplementary Figure S5.2. Pan-genome and recombination in Group 2 (*Ca.* P. pruni, 6 genomes). Display as in Figure S5.1. Heaps’ law exponent 0.84; 936 pan-genome genes and 406 core genes. r/m = 1.068, R/theta = 0.137, 80 events.

Supplementary Figure S5.3. Pan-genome and recombination in Group 3 (*Ca.* P. conii, 5 genomes). Display as in Figure S5.1. Heaps’ law exponent 0.77; 1,222 pan-genome genes and 370 core genes. r/m = 0.590, R/theta = 0.241, 85 events.

Supplementary Figure S5.4. Recombination in Group 1 (*Candidatus* Phytoplasma caryae, 10 genomes). Left, maximum likelihood phylogeny of the core gene alignment (scale bar, substitutions per site). Right, recombination along the core alignment inferred with ClonalFrameML. Blue is the clonal frame and red marks regions inferred to be imported by recombination. Each row is one tip or internal branch of the tree.

Supplementary Figure S5.5. Recombination in Group 2 (*Candidatus* Phytoplasma pruni, 6 genomes). Display as in Figure S5.4.

Supplementary Figure S5.6. Recombination in Group 3 (*Candidatus* Phytoplasma conii, 5 genomes). Display as in Figure S5.4.

Supplementary Figure S6.1. Pathway genes detected in each genome. Number of the 66 curated pathway genes (nucleotide salvage, glycolysis, DNA replication, folate biosynthesis, glycerophospholipid biosynthesis, protein secretion and transport) detected in each of the 22 genomes. Blue, genes detected in the Prokka v1.13 annotation; orange, additional genes detected only by BLASTp.

Supplementary Figure S6.2. Predicted secreted proteins in the 22 genomes. Top, number of proteins with a predicted Sec/SPI signal peptide (SignalP v5.0b, gram-positive setting) in each genome; the percentage above each bar is the share of all predicted proteins. Bottom, the same secreted candidates grouped by BLASTp (DIAMOND v2.1.13, e-value 1e-5, minimum identity 30%) comparison with the reference proteomes: light blue, a homolog was found in at least one reference; orange, no homolog was found.

Supplementary Figure S6.3. Orthogroups per genome in 20 genomes. Number of orthogroups (OrthoFinder v3.1.5) that contain each genome. Dark blue, orthogroups present in all genomes; light blue, orthogroups shared between the PPS strains and the reference strains; orange, orthogroups exclusive to the set (PPS or reference) to which the genome belongs.

Supplementary Figure S6.4. Orthogroups per genome in 22 genomes. Number of orthogroups (OrthoFinder v3.1.5) that contain each genome.

Supplementary Figure S6.5. Orthogroups per genome in 22 genomes, second analysis.

## List of Supplementary Tables

Supplementary Table S1.1. Assembly statistics of the six PPS genomes: genome size, contigs, N50, GC content, coding sequences, tRNA genes, sequencing depth, CheckM estimates and reads mapped.

Supplementary Table S1.2. Assembly statistics and source information (accession, strain, organism name in NCBI, host, location, collection date and isolation source) of the 22 genomes.

Supplementary Table S1.3. Genomes used in this study: the 22 *Ca.* Phytoplasma pruni and *Ca.* P. pruni-related genomes with proposed species, host, origin, accession numbers, reference, assembly statistics and CheckM v1 (lineage_wf) completeness and contamination. PPS strains were sequenced in this study; the other genomes are public.

Supplementary Table S1.4. The 83 curated genomes (CheckM completeness of at least 90% and contamination of at most 5%), including the six PPS genomes sequenced in this study: CheckM values, assembly details, organism name in NCBI and in this study, nearest neighbour genome by ANIm, ANI clusters at the 95% and 96% thresholds, and 16S rRNA gene copies.

Supplementary Table S1.5. *Ca.* P. pruni and *Ca.* P. pruni-related genomes excluded from the curated set because they did not meet the CheckM rule (completeness of at least 90% and contamination of at most 5%), with their CheckM values and the reason for exclusion.

Supplementary Table S1.6. CheckM v1 (lineage_wf) completeness and contamination of all 299 genomes scored (293 genomes downloaded from NCBI, one row per assembly, and the six PPS genomes), with assembly details and organism name in NCBI and in this study. Genomes that meet the CheckM rule (252) were considered for the curated set.

Supplementary Table S2.1. 16S rRNA gene reference sequence of each ANI group, with its length, GC content and the sequences in the group.

Supplementary Table S2.2. Mean, minimum and maximum 16S rRNA gene sequence identity (%) within and between the four ANI groups.

Supplementary Table S2.3. 16S rRNA gene positions at which the sequences of one ANI group share a base that differs from the other groups (diagnostic positions).

Supplementary Table S2.4. Base at each diagnostic position of the 16S rRNA gene in every 16S rRNA gene sequence.

Supplementary Table S2.5. Pairwise identity (%) between all 16S rRNA gene sequences. Supplementary Table S2.6. 16S rRNA gene identity (%) of the reference sequences of *Ca.* P. caryae (PPS1A), *Ca.* P. conii (OVSP-L01 and CicWB-2022), *Ca.* P. lactucae (LWB) and *Ca.* P. pruni (CX) to the 16S rRNA gene sequences of the 60 described ‘*Candidatus* Phytoplasma’ species that have a sequence in GenBank.

Supplementary Table S3.1. FastANI (%) between the 22 genomes. Supplementary Table S3.2. ANIb (%) between the 22 genomes.

Supplementary Table S3.3. ANIm (%) between the 22 genomes.

Supplementary Table S3.4. ANIm, ANIb and FastANI between *Acholeplasma laidlawii* PG-8A and each of the 22 genomes.

Supplementary Table S3.5. Range of pairwise ANI (%) within and between the four ANI groups, by method.

Supplementary Table S4.1. Digital DNA-DNA hybridization (dDDH) between all pairs of the 22 genomes by GGDC formulas 1, 2 and 3, with confidence intervals, intergenomic distances and the probability that dDDH is at least 70%.

Supplementary Table S4.2. dDDH (%) between the 22 genomes, GGDC formula 1. Supplementary Table S4.3. dDDH (%) between the 22 genomes, GGDC formula 2. Supplementary Table S4.4. dDDH (%) between the 22 genomes, GGDC formula 3.

Supplementary Table S4.5. F2 dDDH (%), minimum to maximum between lineages (within a lineage on the diagonal).

Supplementary Table S4.6. Genome clusters at the 70% dDDH threshold for each GGDC formula.

Supplementary Table S4.7. Genome pairs for which GGDC formula 2 and formulas 1 or 3 fall on opposite sides of the 70% dDDH threshold.

Supplementary Table S5.1. GTDB-Tk classification of the 22 pruni and pruni-related genomes.

Supplementary Table S5.2. GTDB-Tk classification of 53 genomes: the 22 pruni and pruni-related genomes and 31 representative genomes of the other ANIm clusters.

Supplementary Table S5.3. The 53 genomes classified by GTDB-Tk with their ANIm cluster (95% ANI threshold) and GTDB species.

Supplementary Table S6.1. Location of the five potential mobile unit (PMU) genes (*sigF*, *ssb*, *hupB*, *ftsH* and *smc*) in each genome: whether they lie in one window on one contig, the span and gene order of the window, and the number of loci per gene.

Supplementary Table S6.2. Contig, position and strand of the five PMU genes inside the window of each genome.

Supplementary Table S6.3. PMU genes identified by gene name in the Prokka annotation of each genome.

Supplementary Table S6.4. Number of copies of each PMU gene in each genome, from detection by gene name.

Supplementary Table S6.5. BLASTp identity of each PMU window protein to the PMU window protein of PPS1A with the same name.

Supplementary Table S7.1. Metabolic genes searched, with pathway, EC numbers, gene names and product keywords used.

Supplementary Table S7.2. Detection call (P, B, x or -) for each metabolic gene in each of the 22 genomes; the codes are defined in the Legend sheet.

Supplementary Table S7.3. Detection call for each metabolic gene in PPS1A, PPS2B, PPS6A and PPS6B, with the number of strains detected by Prokka (P), by BLASTp only (B), or by either.

Supplementary Table S7.4. Number of metabolic genes detected in each genome.

Supplementary Table S7.5. Presence (1) or absence (0) of each metabolic gene in each of the 22 genomes, with the class of the gene across the genomes.

Supplementary Table S7.6. Metabolic genes that are specific to, or absent from, an ANI group relative to the other genomes.

Supplementary Table S7.7. Number of metabolic genes in each class (specific to or absent from a group) for each ANI group.

Supplementary Table S7.8. Prokka annotation evidence (locus tag, gene name and matching criterion) for each detected metabolic gene.

Supplementary Table S7.9. BLASTp result and final call for every metabolic gene in every genome.

Supplementary Table S7.10. BLASTp result and final call for the metabolic genes that Prokka did not detect but that have a BLASTp hit.

Supplementary Table S8.1. SignalP classification of the proteins of each of the 22 genomes: number of proteins in each class and share of secreted proteins.

Supplementary Table S8.2. SignalP classification of the proteins of PPS1A, PPS2B, PPS6A and PPS6B: number of proteins in each class and share of secreted proteins.

Supplementary Table S8.3. Secreted proteins (SP, Sec/SPI) and lipoproteins (LIPO, Sec/SPII) predicted by SignalP in the 22 genomes with SignalP probabilities, cleavage site position, protein length and annotated product.

Supplementary Table S8.4. Secreted proteins (SP, Sec/SPI) and lipoproteins (LIPO, Sec/SPII) predicted by SignalP in PPS1A, PPS2B, PPS6A and PPS6B with SignalP probabilities, cleavage site position, protein length and annotated product.

Supplementary Table S9.1. Number of secreted protein candidates of each genome with a hit in the other genomes and in the reference genomes.

Supplementary Table S9.2. Number of secreted protein candidates of PPS1A, PPS2B, PPS6A and PPS6B with a hit in the other genomes and in the reference genomes.

Supplementary Table S9.3. Secreted protein candidates of each genome, with the number of other and reference genomes with a hit, the best identity and the genomes with a hit.

Supplementary Table S9.4. Secreted protein candidates of PPS1A, PPS2B, PPS6A and PPS6B, with the number of other and reference genomes with a hit, the best identity and the genomes with a hit.

Supplementary Table S9.5. Secreted protein candidates of the six PPS genomes with no hit in any reference genome.

Supplementary Table S10.1. Call for each effector gene (present, partial, paralog, contig-break, stop codon or absent) in each of the 22 genomes.

Supplementary Table S10.2. Call for each effector gene in each of the 22 genomes, with identity and the number of additional loci of any call type.

Supplementary Table S10.3. Call for each effector gene in the four PPS strains PPS1A, PPS2B, PPS6A and PPS6B.

Supplementary Table S10.4. Call for each effector gene in PPS1A, PPS2B, PPS6A and PPS6B, with identity and the number of additional loci of any call type.

Supplementary Table S10.5. Search result for every effector in every genome: call, TBLASTN identity and query coverage, position on the contig, and BLASTp identity and coverage.

Supplementary Table S10.6. Search result for every effector in PPS1A, PPS2B, PPS6A and PPS6B: call, TBLASTN identity and query coverage, position on the contig, and BLASTp identity and coverage.

## List of Abbreviations

*Ca.* P.: *Candidatus* Phytoplasma
IRPCM: International Research Programme on Comparative Mycoplasmology
NCBI: National Center for Biotechnology Information
PBT: Pecan bunchy top phytoplasma

