## Supplementary material for "Taxogenomic resolution of the 16SrIII (X-disease) group: elevation of ‘*Candidatus* Phytoplasma pruni’-related strains to species rank and proposal of three novel *Candidatus* species ‘*Candidatus* Phytoplasma caryae’ sp. nov., ‘*Candidatus* Phytoplasma conii’ sp. nov. and ‘*Candidatus* Phytoplasma": Rabiu et al., Supplementary figures and Tables: Rabiu_et_al_Supplementary_File_1.pptx

### Slide 1
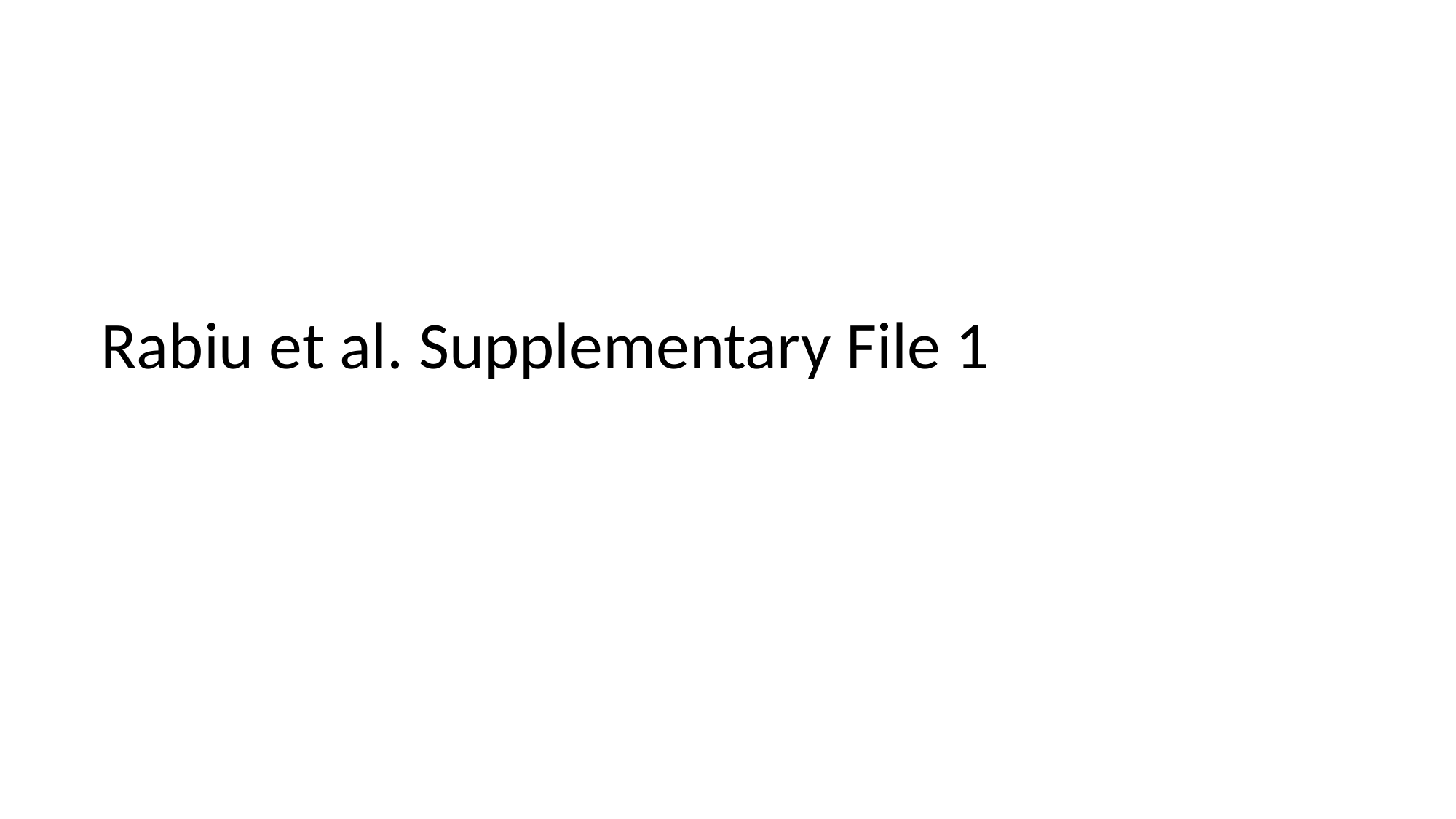

#### Rabiu et al. Supplementary File 1

### Slide 2
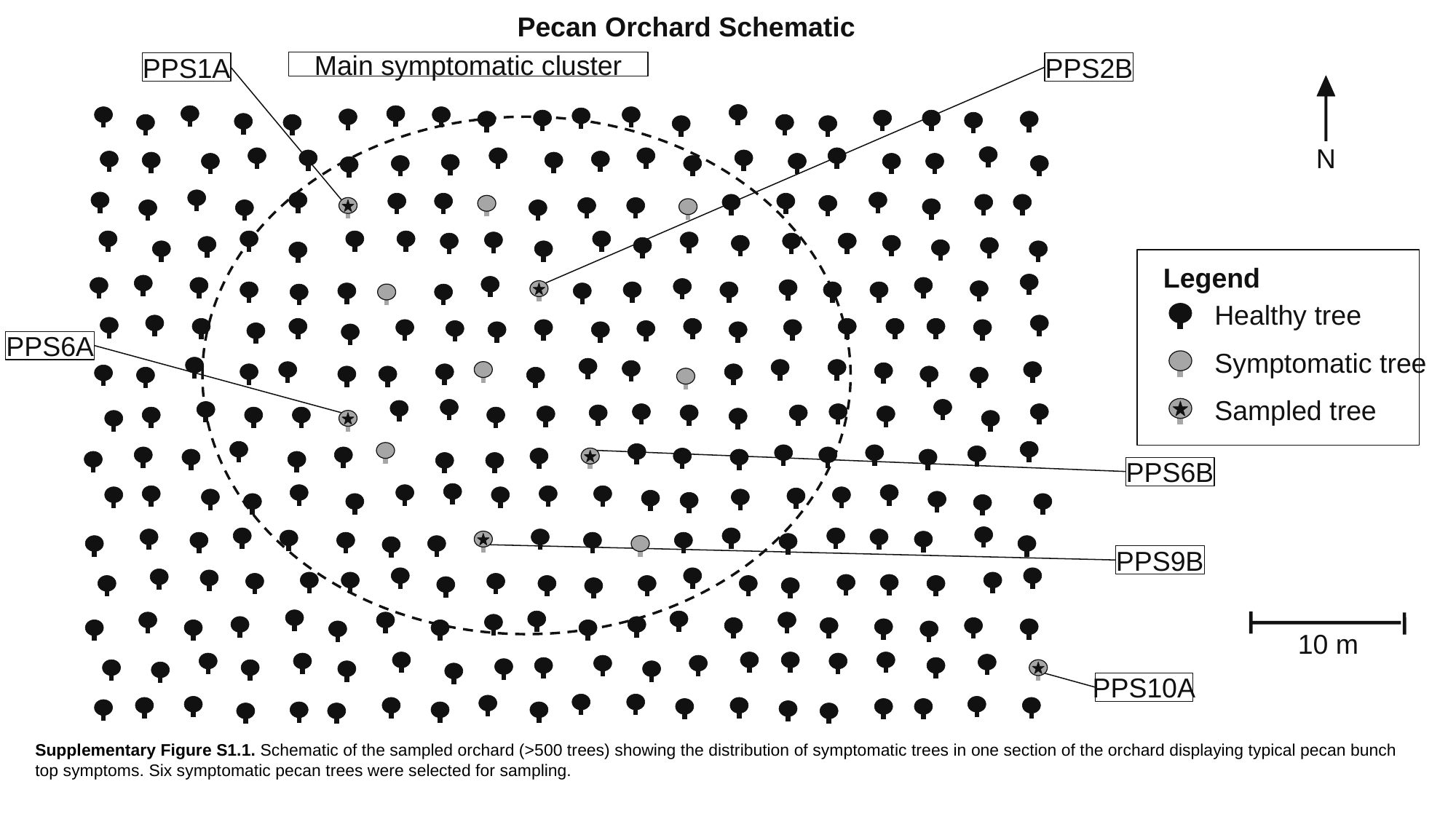

Pecan Orchard Schematic
Main symptomatic cluster
PPS1A
PPS2B
N
Legend
Healthy tree
PPS6A
Symptomatic tree
Sampled tree
PPS6B
PPS9B
10 m
PPS10A
Supplementary Figure S1.1. Schematic of the sampled orchard (>500 trees) showing the distribution of symptomatic trees in one section of the orchard displaying typical pecan bunch top symptoms. Six symptomatic pecan trees were selected for sampling.

### Slide 3
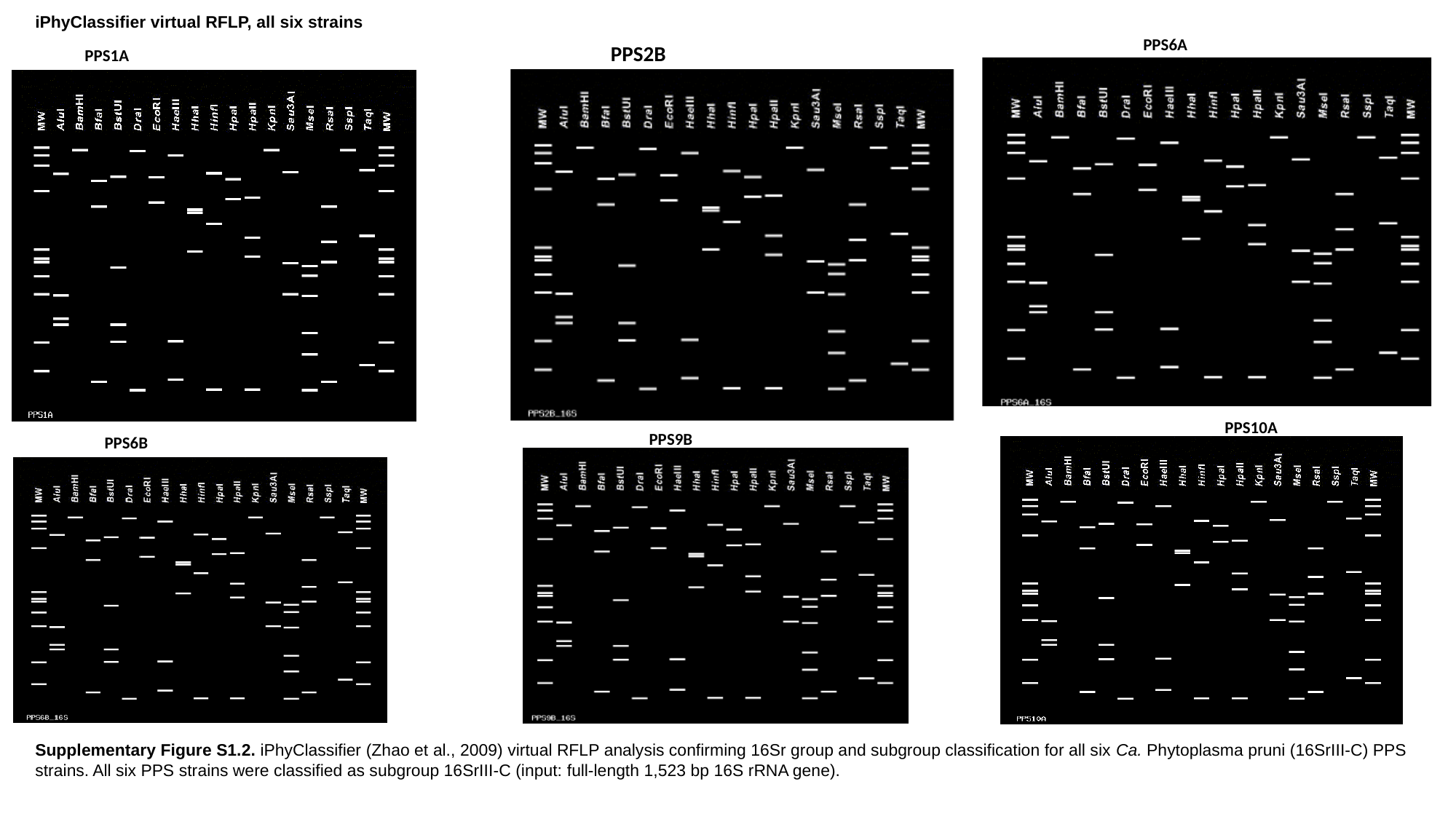

iPhyClassifier virtual RFLP, all six strains
PPS6A
#### PPS2B
PPS1A
PPS10A
PPS9B
PPS6B
Supplementary Figure S1.2. iPhyClassifier (Zhao et al., 2009) virtual RFLP analysis confirming 16Sr group and subgroup classification for all six Ca. Phytoplasma pruni (16SrIII-C) PPS strains. All six PPS strains were classified as subgroup 16SrIII-C (input: full-length 1,523 bp 16S rRNA gene).

### Slide 4
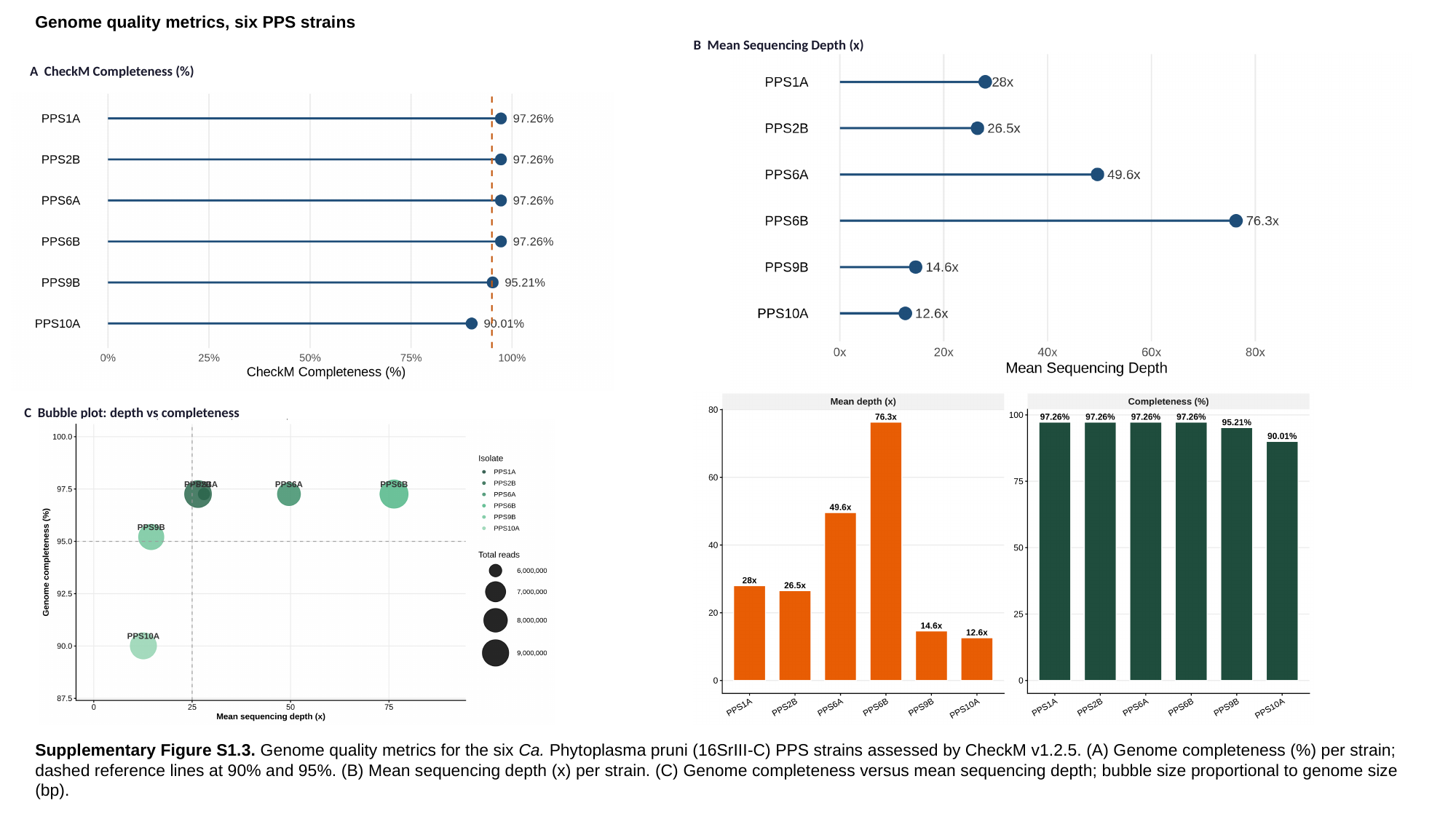

Genome quality metrics, six PPS strains
B Mean Sequencing Depth (x)
A CheckM Completeness (%)
C Bubble plot: depth vs completeness
Supplementary Figure S1.3. Genome quality metrics for the six Ca. Phytoplasma pruni (16SrIII-C) PPS strains assessed by CheckM v1.2.5. (A) Genome completeness (%) per strain; dashed reference lines at 90% and 95%. (B) Mean sequencing depth (x) per strain. (C) Genome completeness versus mean sequencing depth; bubble size proportional to genome size (bp).

### Slide 5
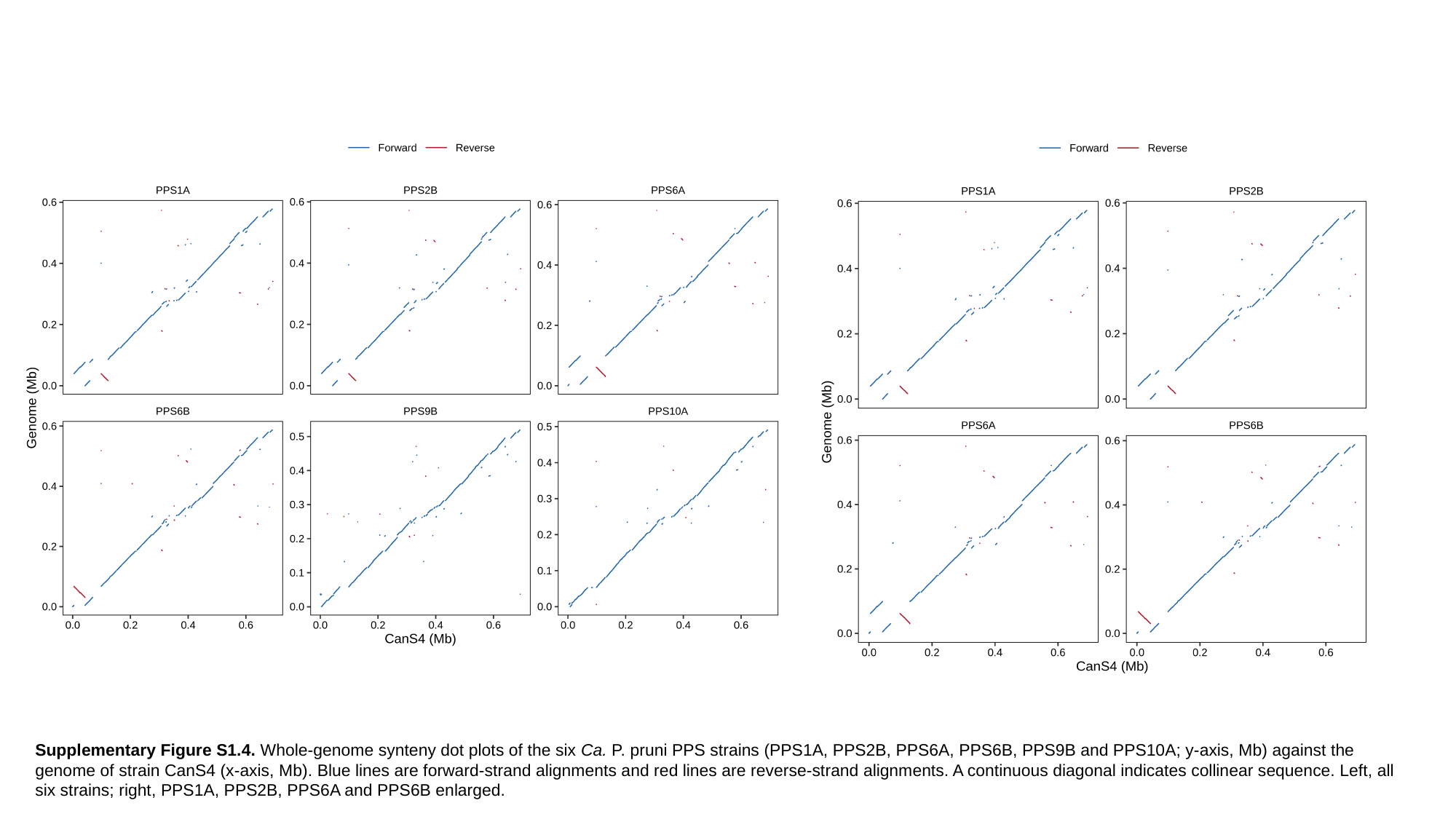

Supplementary Figure S1.4. Whole-genome synteny dot plots of the six Ca. P. pruni PPS strains (PPS1A, PPS2B, PPS6A, PPS6B, PPS9B and PPS10A; y-axis, Mb) against the genome of strain CanS4 (x-axis, Mb). Blue lines are forward-strand alignments and red lines are reverse-strand alignments. A continuous diagonal indicates collinear sequence. Left, all six strains; right, PPS1A, PPS2B, PPS6A and PPS6B enlarged.

### Slide 6
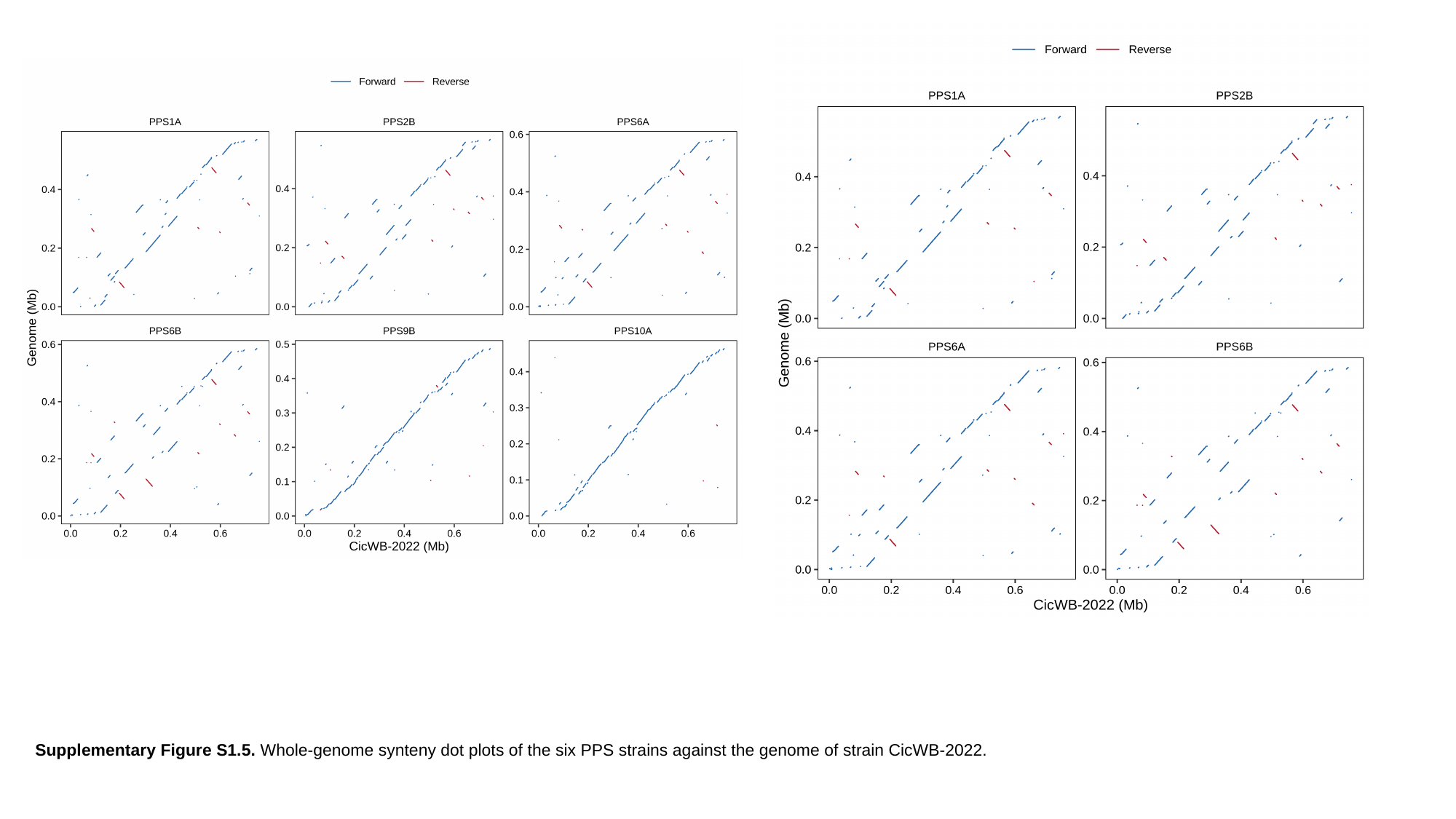

Supplementary Figure S1.5. Whole-genome synteny dot plots of the six PPS strains against the genome of strain CicWB-2022.

### Slide 7
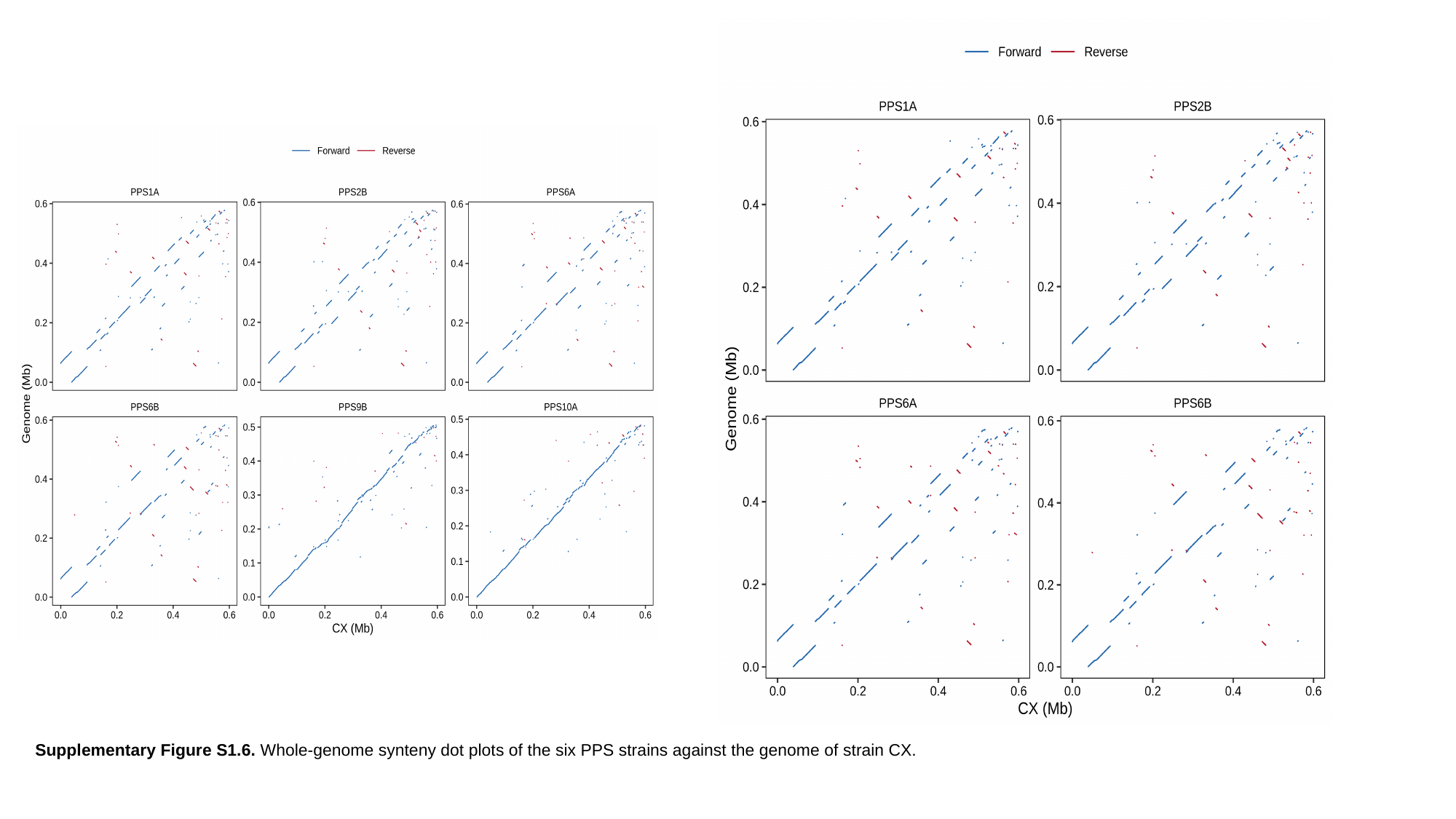

Supplementary Figure S1.6. Whole-genome synteny dot plots of the six PPS strains against the genome of strain CX.

### Slide 8
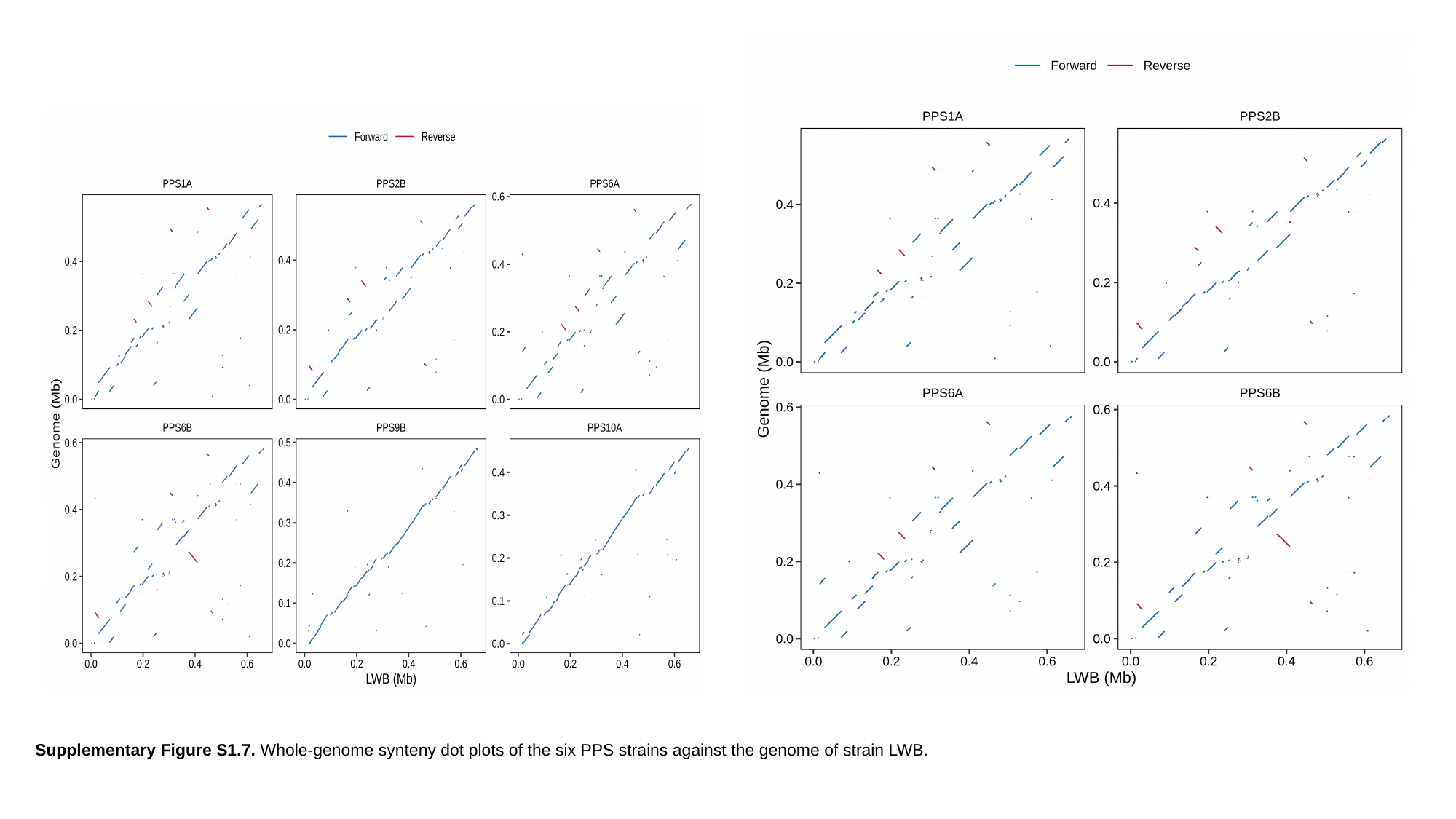

Supplementary Figure S1.7. Whole-genome synteny dot plots of the six PPS strains against the genome of strain LWB.
