## Supplementary material for "Taxogenomic resolution of the 16SrIII (X-disease) group: elevation of ‘*Candidatus* Phytoplasma pruni’-related strains to species rank and proposal of three novel *Candidatus* species ‘*Candidatus* Phytoplasma caryae’ sp. nov., ‘*Candidatus* Phytoplasma conii’ sp. nov. and ‘*Candidatus* Phytoplasma": Rabiu et al., Supplementary figures and Tables: Rabiu_et_al_Supplementary_File_2.pptx

### Slide 1
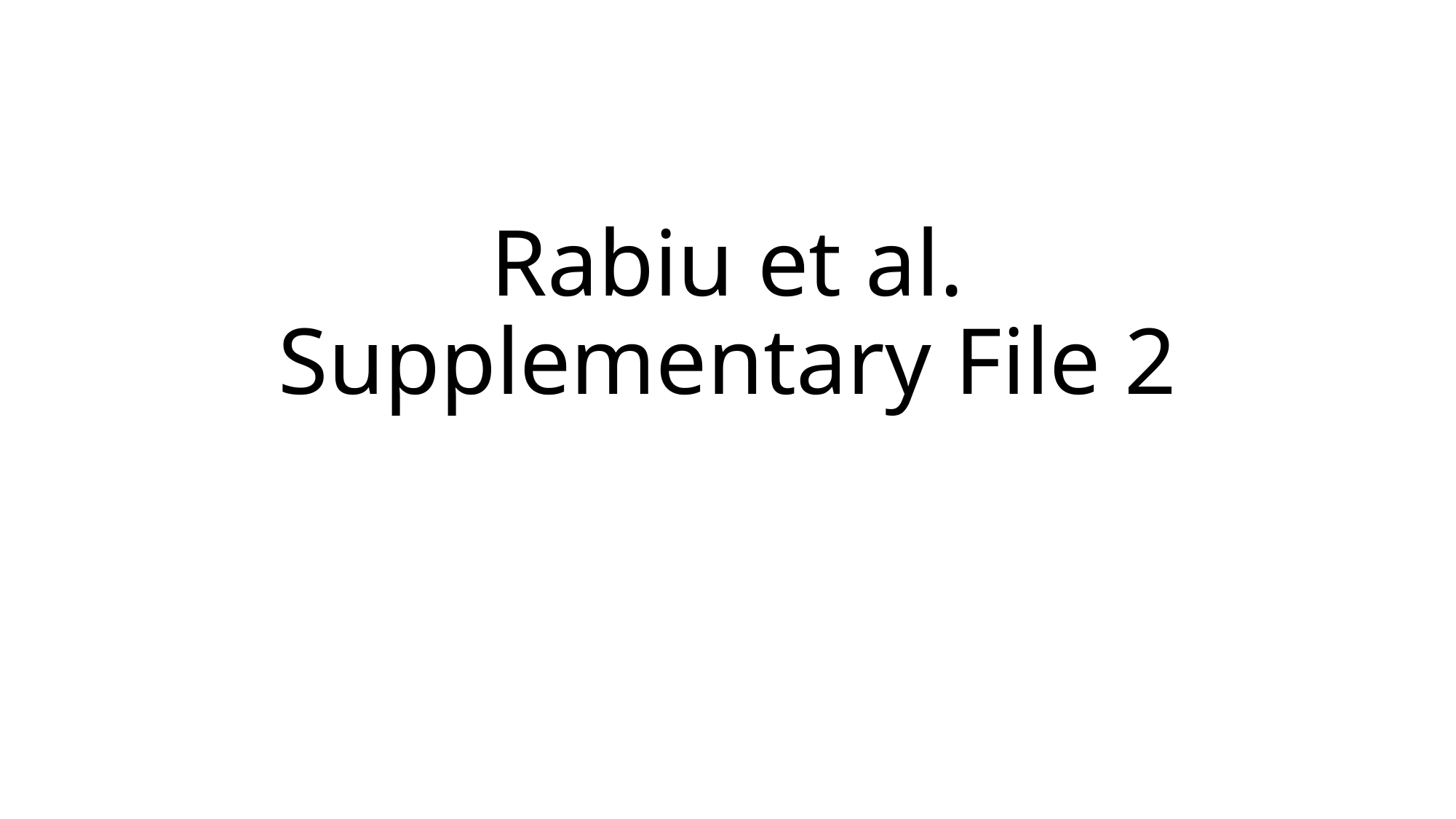

#### Rabiu et al. Supplementary File 2

### Slide 2
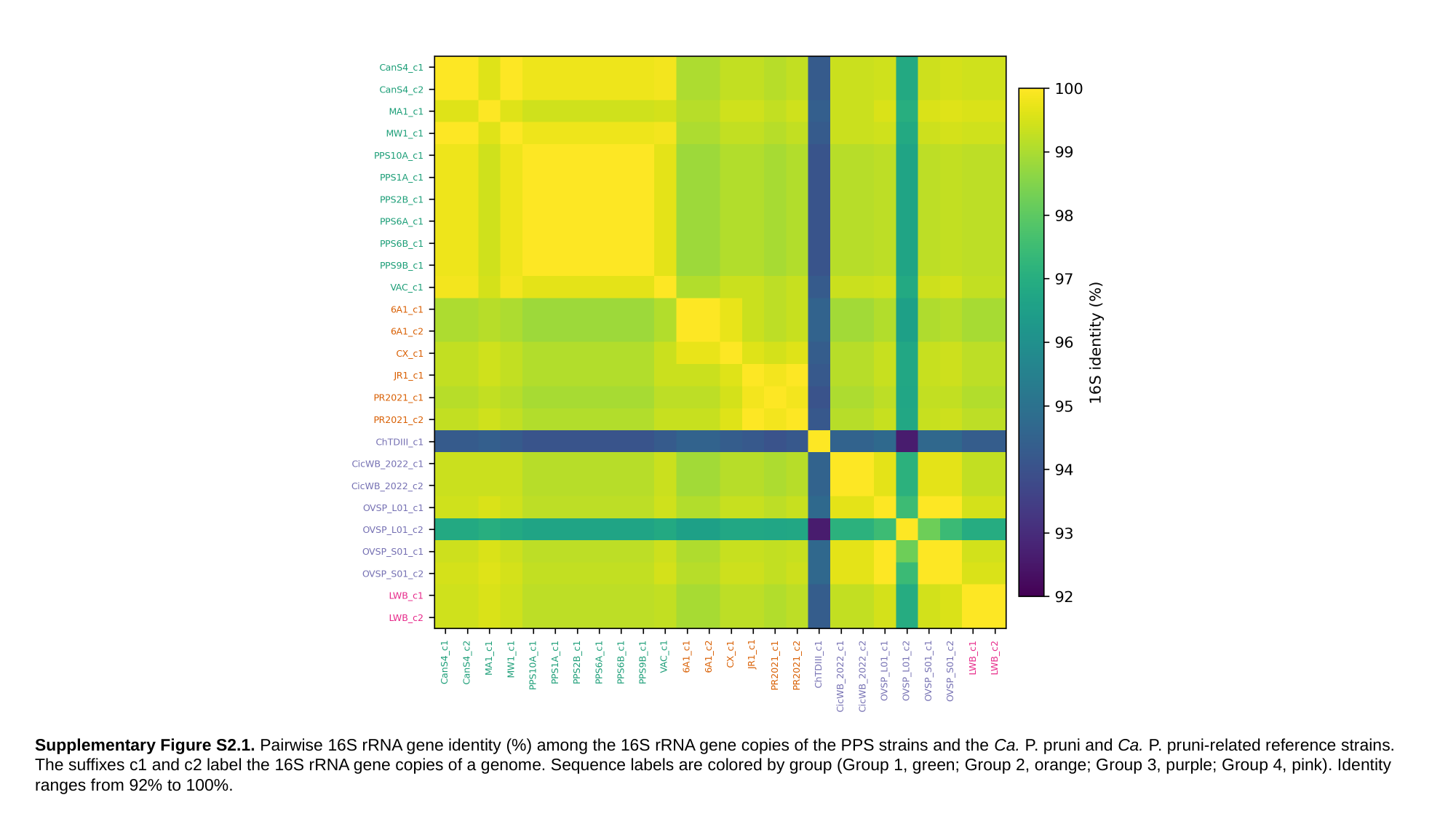

Supplementary Figure S2.1. Pairwise 16S rRNA gene identity (%) among the 16S rRNA gene copies of the PPS strains and the Ca. P. pruni and Ca. P. pruni-related reference strains. The suffixes c1 and c2 label the 16S rRNA gene copies of a genome. Sequence labels are colored by group (Group 1, green; Group 2, orange; Group 3, purple; Group 4, pink). Identity ranges from 92% to 100%.

### Slide 3
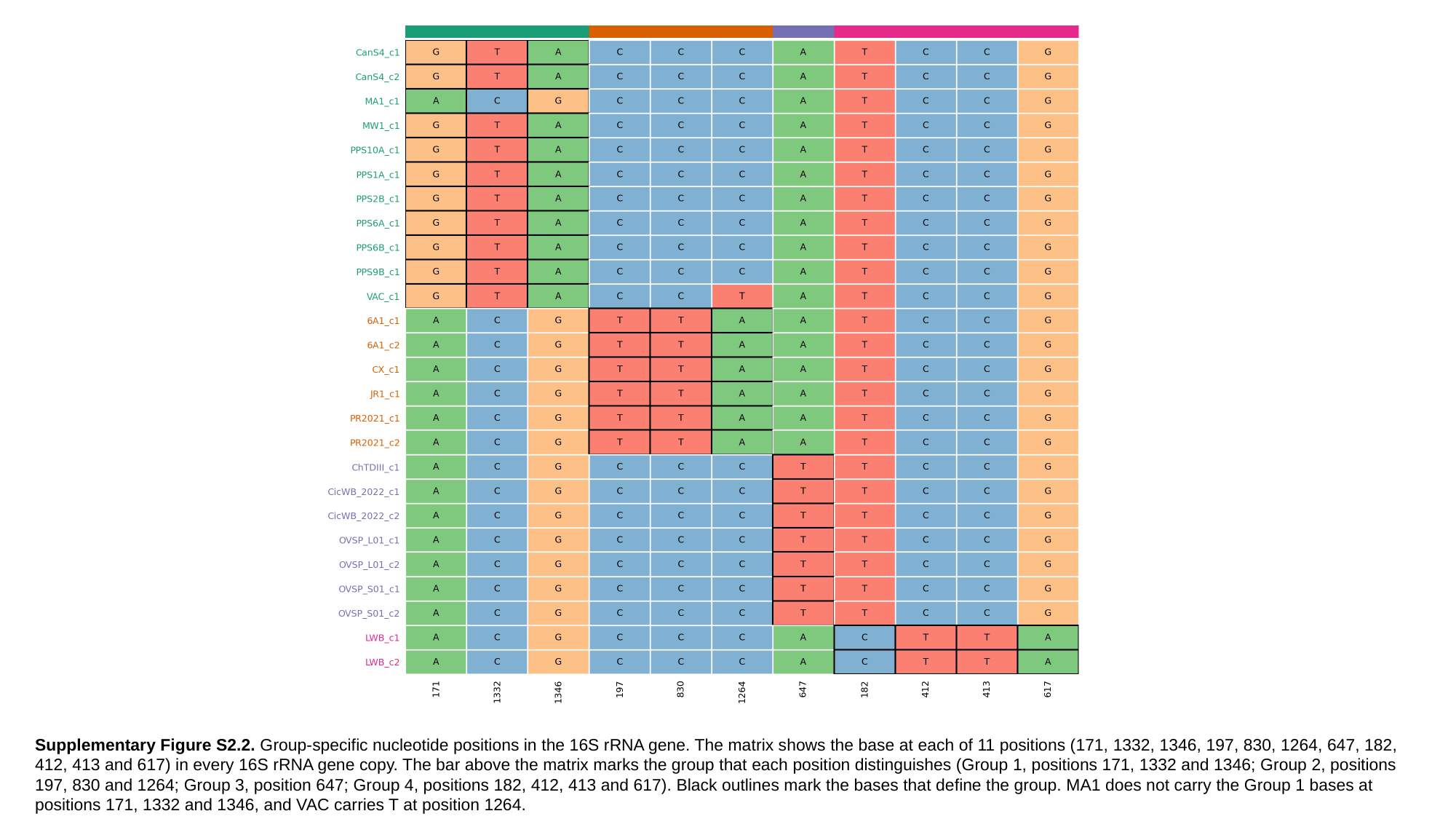

Supplementary Figure S2.2. Group-specific nucleotide positions in the 16S rRNA gene. The matrix shows the base at each of 11 positions (171, 1332, 1346, 197, 830, 1264, 647, 182, 412, 413 and 617) in every 16S rRNA gene copy. The bar above the matrix marks the group that each position distinguishes (Group 1, positions 171, 1332 and 1346; Group 2, positions 197, 830 and 1264; Group 3, position 647; Group 4, positions 182, 412, 413 and 617). Black outlines mark the bases that define the group. MA1 does not carry the Group 1 bases at positions 171, 1332 and 1346, and VAC carries T at position 1264.

### Slide 4
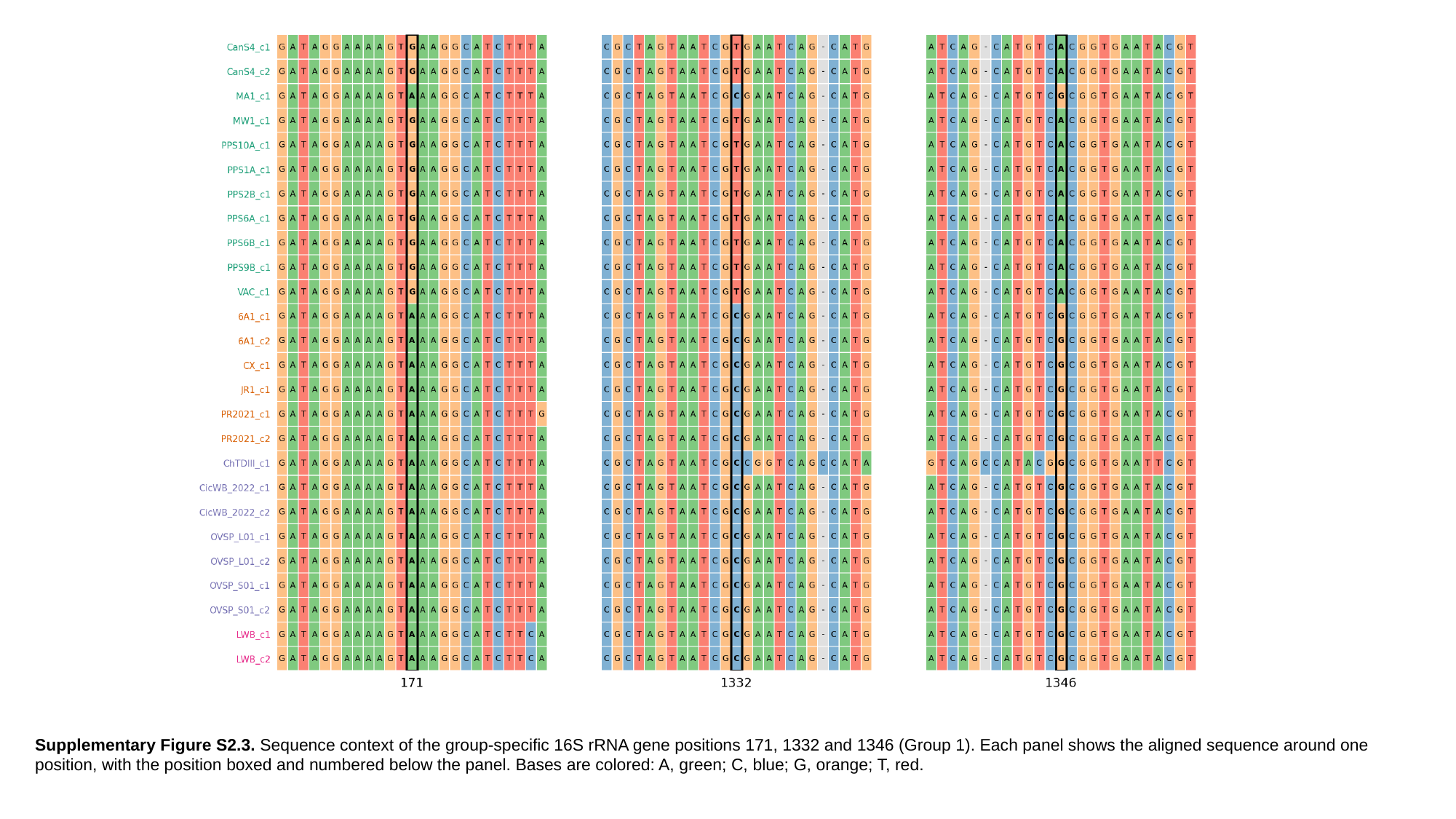

Supplementary Figure S2.3. Sequence context of the group-specific 16S rRNA gene positions 171, 1332 and 1346 (Group 1). Each panel shows the aligned sequence around one position, with the position boxed and numbered below the panel. Bases are colored: A, green; C, blue; G, orange; T, red.

### Slide 5
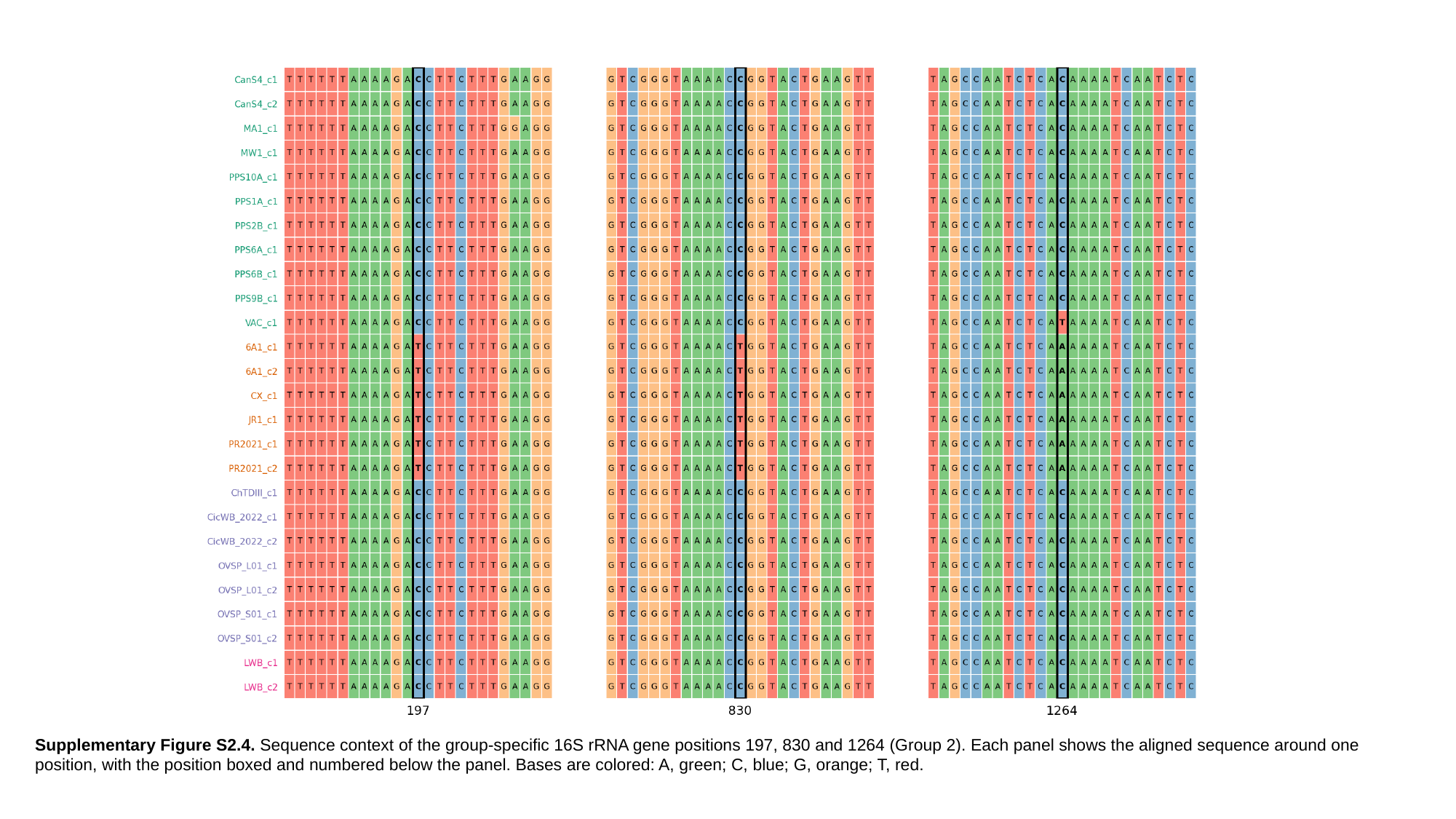

Supplementary Figure S2.4. Sequence context of the group-specific 16S rRNA gene positions 197, 830 and 1264 (Group 2). Each panel shows the aligned sequence around one position, with the position boxed and numbered below the panel. Bases are colored: A, green; C, blue; G, orange; T, red.

### Slide 6
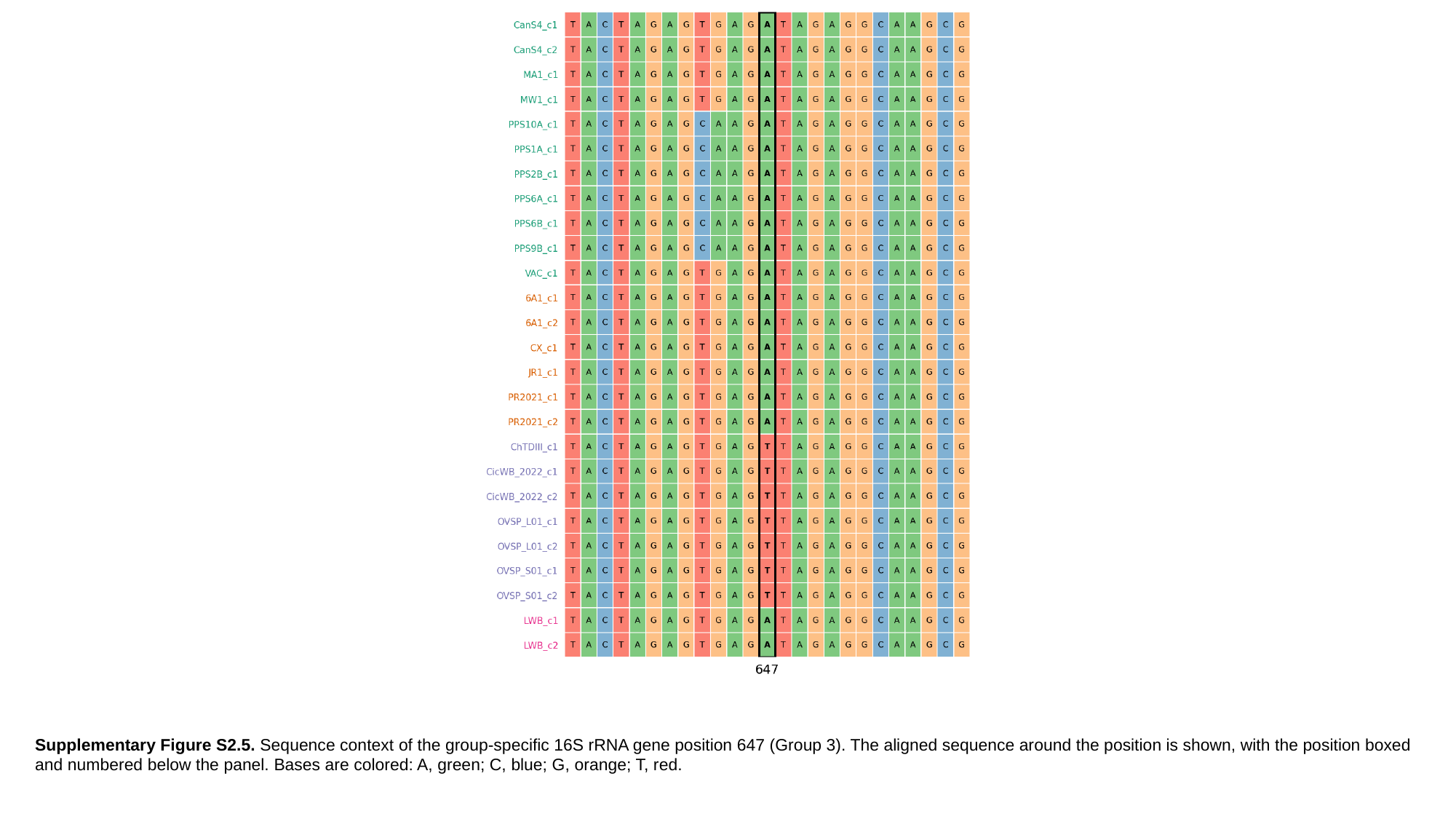

Supplementary Figure S2.5. Sequence context of the group-specific 16S rRNA gene position 647 (Group 3). The aligned sequence around the position is shown, with the position boxed and numbered below the panel. Bases are colored: A, green; C, blue; G, orange; T, red.

### Slide 7
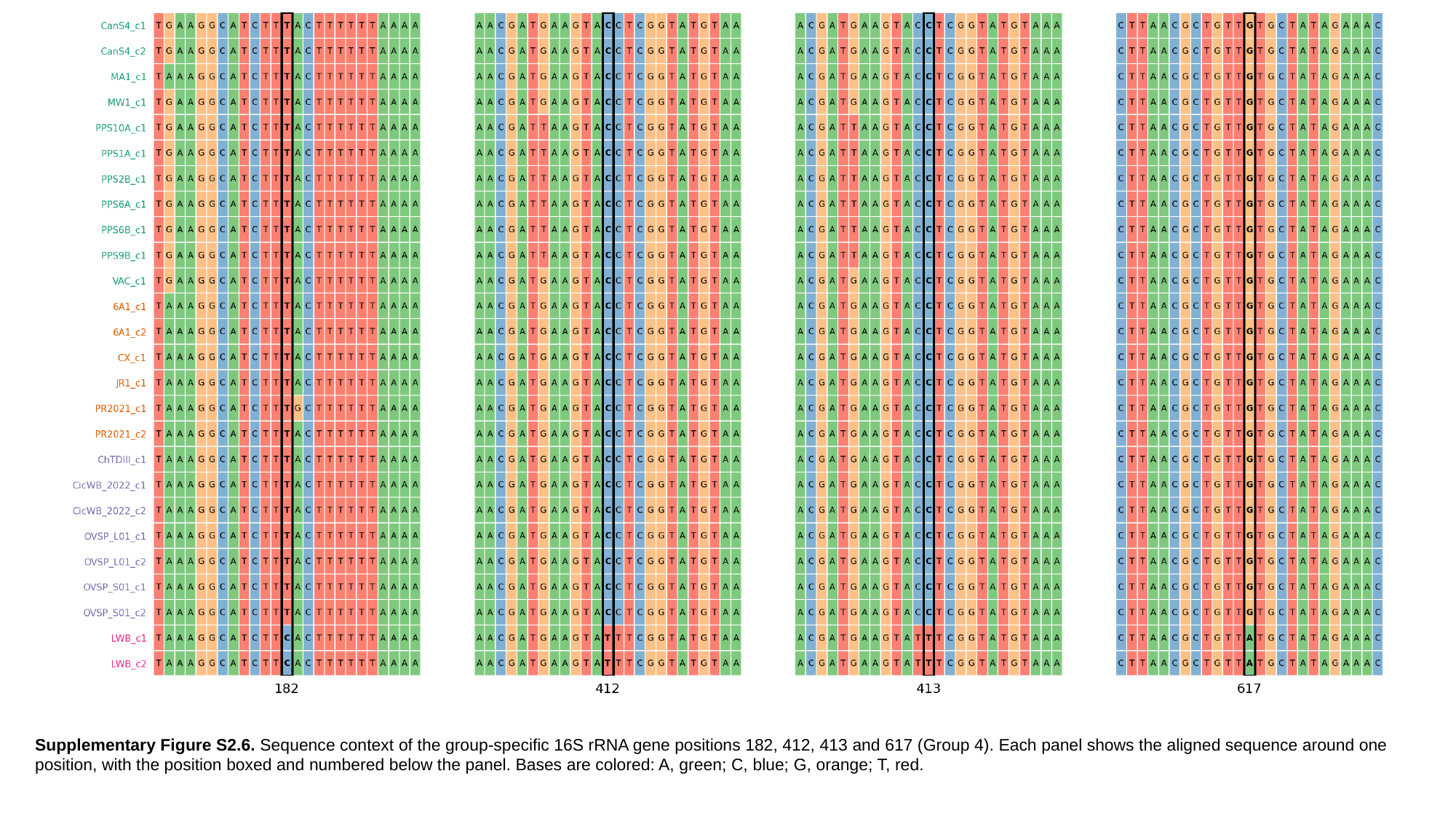

Supplementary Figure S2.6. Sequence context of the group-specific 16S rRNA gene positions 182, 412, 413 and 617 (Group 4). Each panel shows the aligned sequence around one position, with the position boxed and numbered below the panel. Bases are colored: A, green; C, blue; G, orange; T, red.

### Slide 8
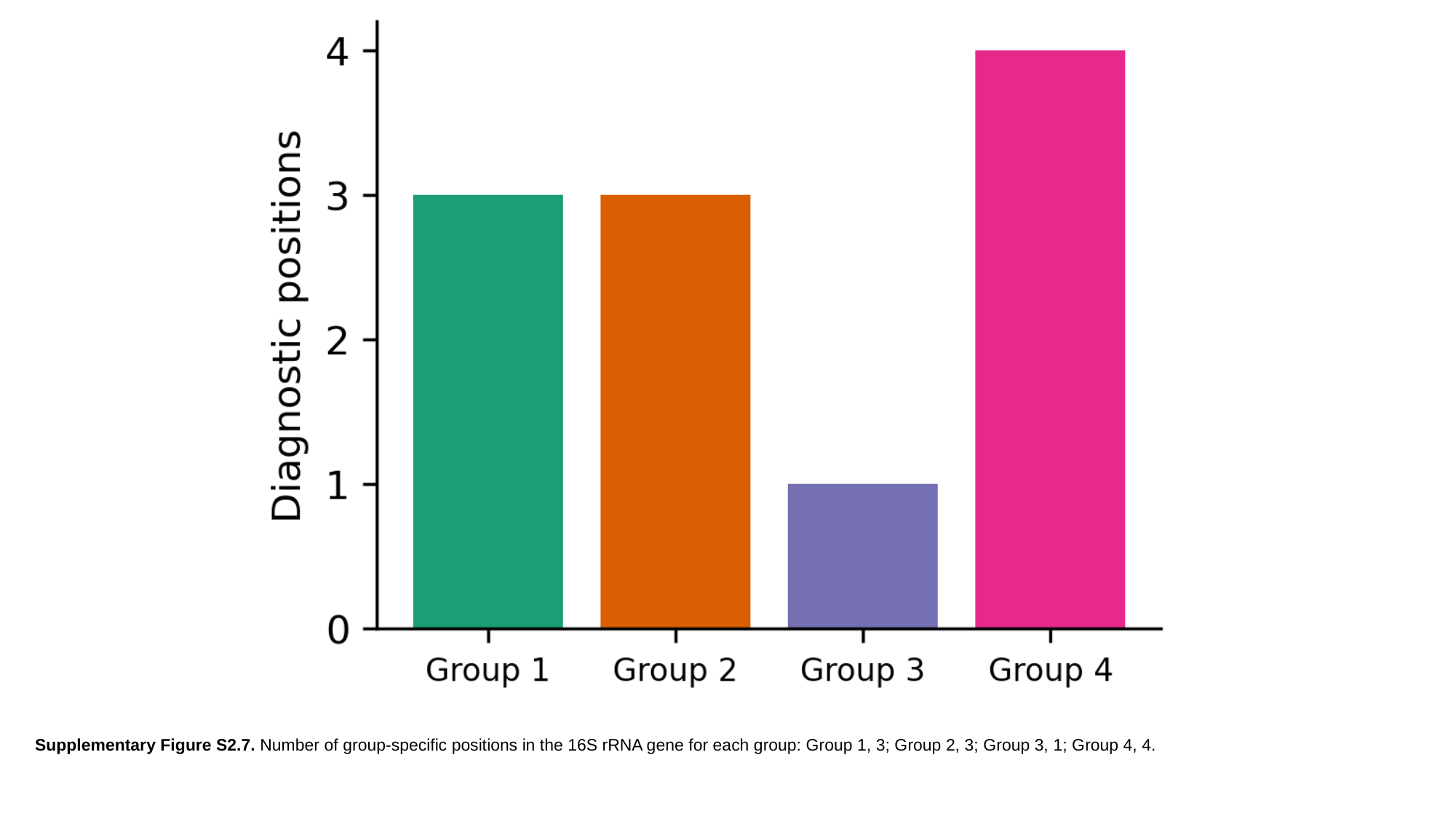

Supplementary Figure S2.7. Number of group-specific positions in the 16S rRNA gene for each group: Group 1, 3; Group 2, 3; Group 3, 1; Group 4, 4.

### Slide 9
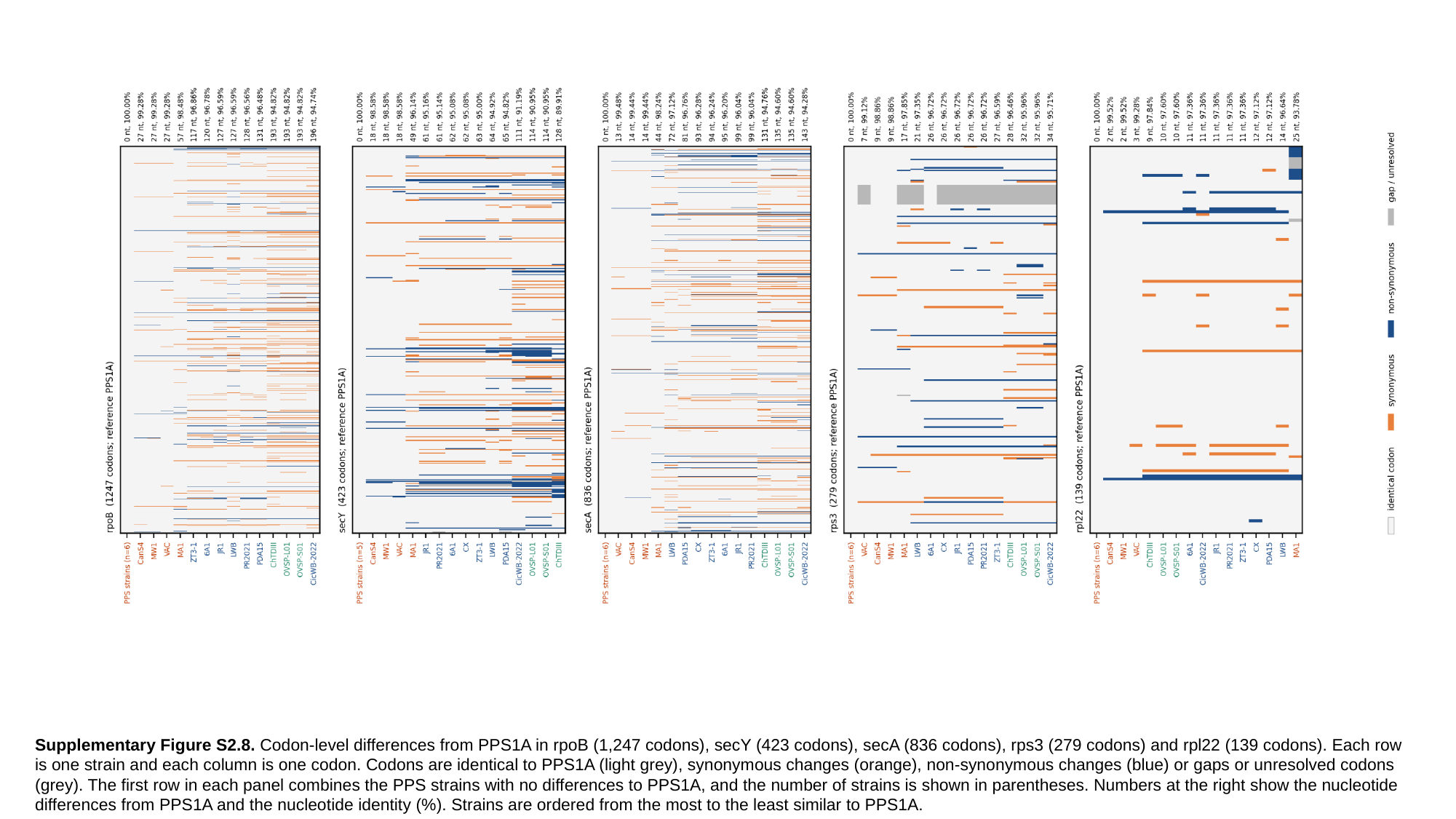

Supplementary Figure S2.8. Codon-level differences from PPS1A in rpoB (1,247 codons), secY (423 codons), secA (836 codons), rps3 (279 codons) and rpl22 (139 codons). Each row is one strain and each column is one codon. Codons are identical to PPS1A (light grey), synonymous changes (orange), non-synonymous changes (blue) or gaps or unresolved codons (grey). The first row in each panel combines the PPS strains with no differences to PPS1A, and the number of strains is shown in parentheses. Numbers at the right show the nucleotide differences from PPS1A and the nucleotide identity (%). Strains are ordered from the most to the least similar to PPS1A.
