## Supplementary material for "Taxogenomic resolution of the 16SrIII (X-disease) group: elevation of ‘*Candidatus* Phytoplasma pruni’-related strains to species rank and proposal of three novel *Candidatus* species ‘*Candidatus* Phytoplasma caryae’ sp. nov., ‘*Candidatus* Phytoplasma conii’ sp. nov. and ‘*Candidatus* Phytoplasma": Rabiu et al., Supplementary figures and Tables: Rabiu_et_al_Supplementary_File_3.pptx

### Slide 1
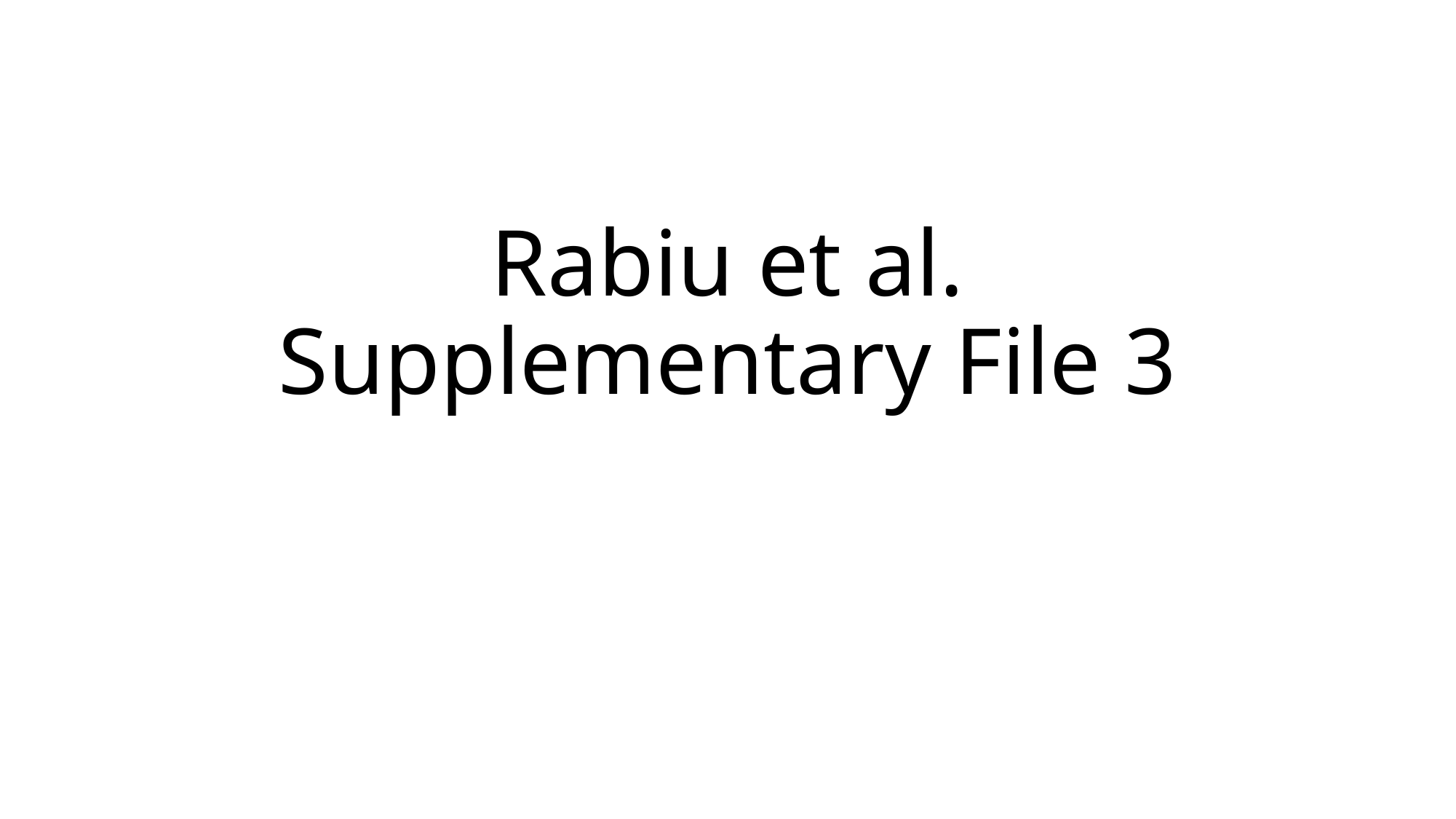

#### Rabiu et al. Supplementary File 3

### Slide 2
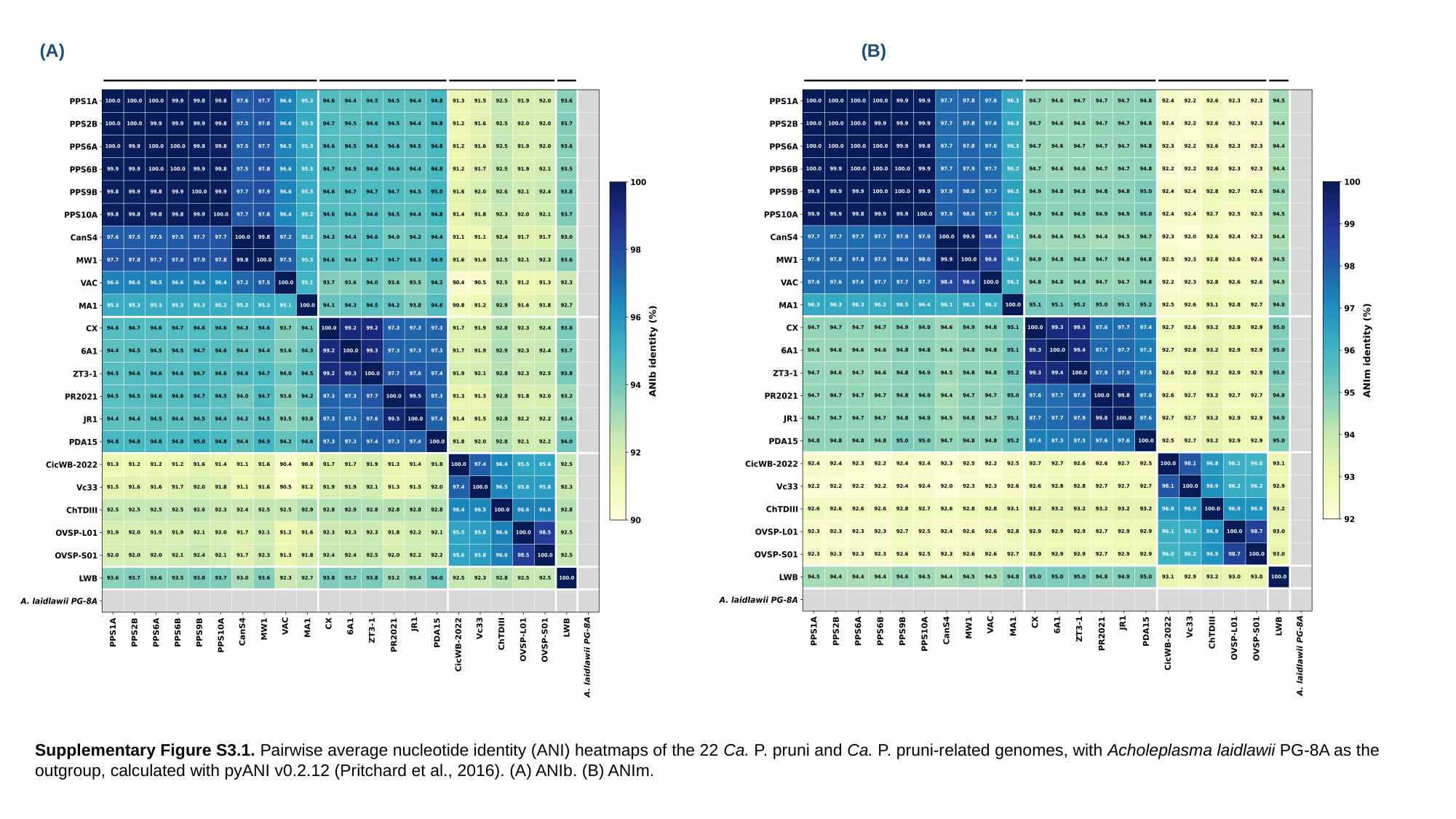

(A)
(B)
Supplementary Figure S3.1. Pairwise average nucleotide identity (ANI) heatmaps of the 22 Ca. P. pruni and Ca. P. pruni-related genomes, with Acholeplasma laidlawii PG-8A as the outgroup, calculated with pyANI v0.2.12 (Pritchard et al., 2016). (A) ANIb. (B) ANIm.

### Slide 3
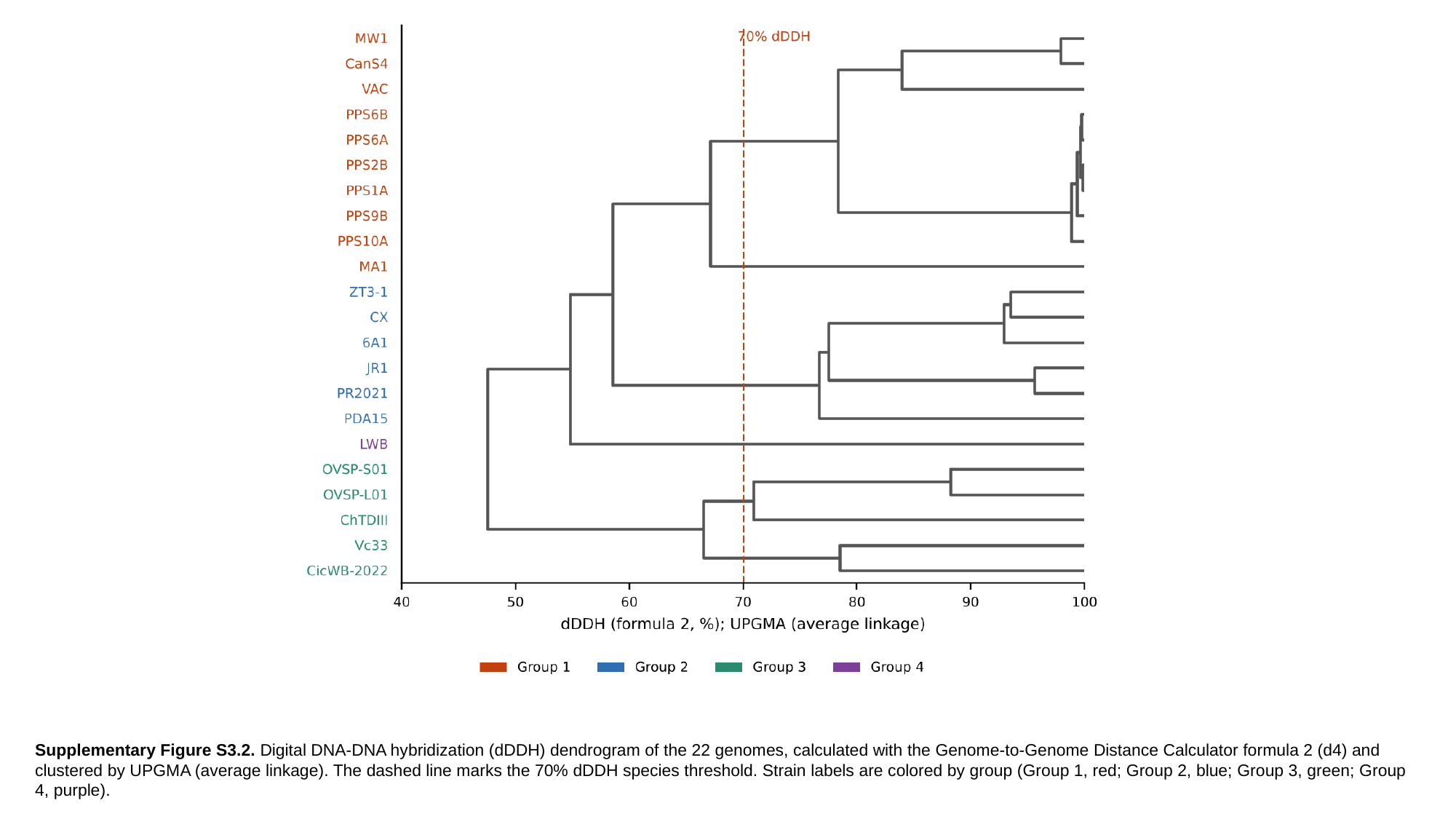

Supplementary Figure S3.2. Digital DNA-DNA hybridization (dDDH) dendrogram of the 22 genomes, calculated with the Genome-to-Genome Distance Calculator formula 2 (d4) and clustered by UPGMA (average linkage). The dashed line marks the 70% dDDH species threshold. Strain labels are colored by group (Group 1, red; Group 2, blue; Group 3, green; Group 4, purple).

### Slide 4
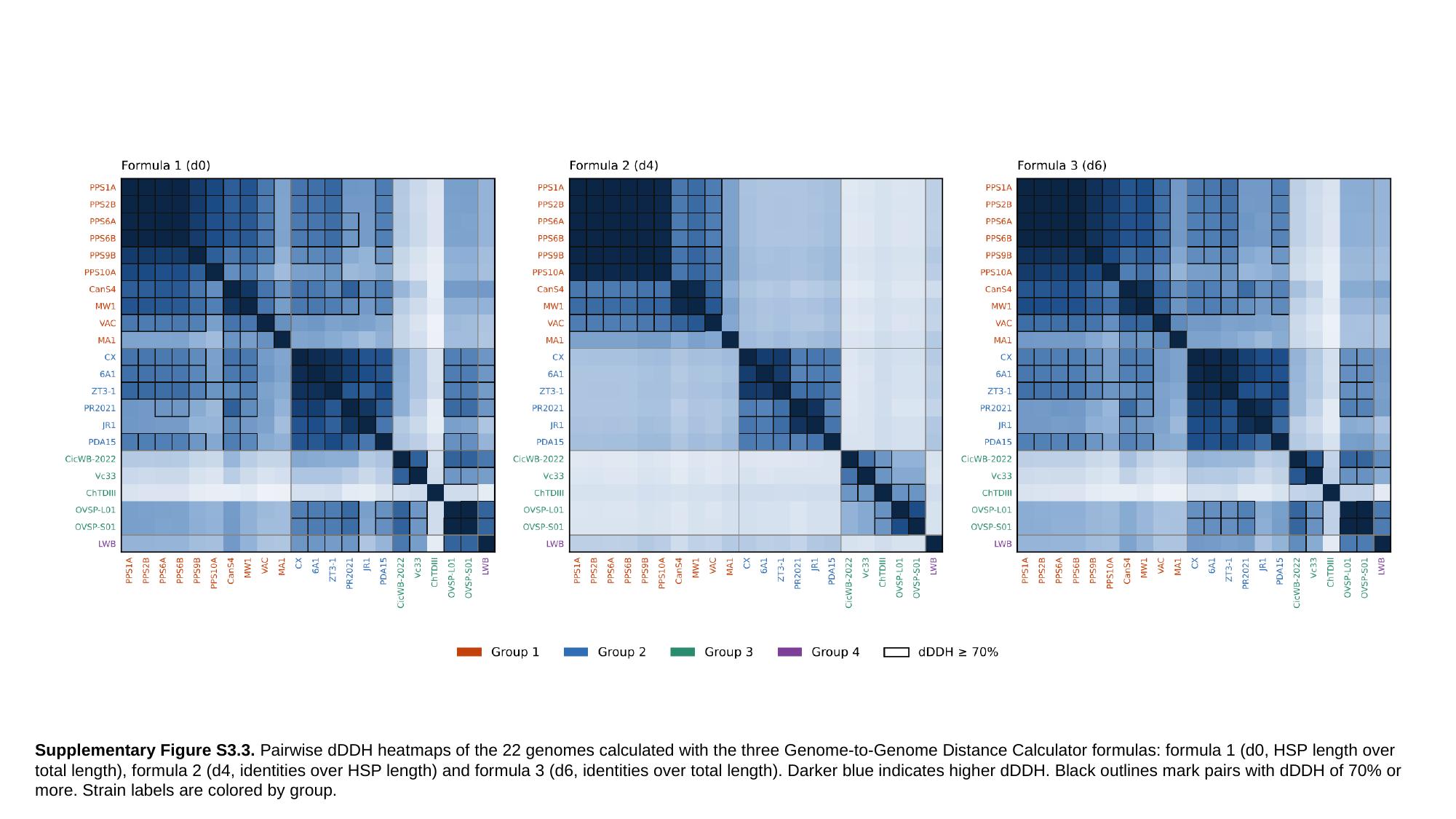

Supplementary Figure S3.3. Pairwise dDDH heatmaps of the 22 genomes calculated with the three Genome-to-Genome Distance Calculator formulas: formula 1 (d0, HSP length over total length), formula 2 (d4, identities over HSP length) and formula 3 (d6, identities over total length). Darker blue indicates higher dDDH. Black outlines mark pairs with dDDH of 70% or more. Strain labels are colored by group.
