## Supplementary material for "Taxogenomic resolution of the 16SrIII (X-disease) group: elevation of ‘*Candidatus* Phytoplasma pruni’-related strains to species rank and proposal of three novel *Candidatus* species ‘*Candidatus* Phytoplasma caryae’ sp. nov., ‘*Candidatus* Phytoplasma conii’ sp. nov. and ‘*Candidatus* Phytoplasma": Rabiu et al., Supplementary figures and Tables: Rabiu_et_al_Supplementary_File_4.pptx

### Slide 1
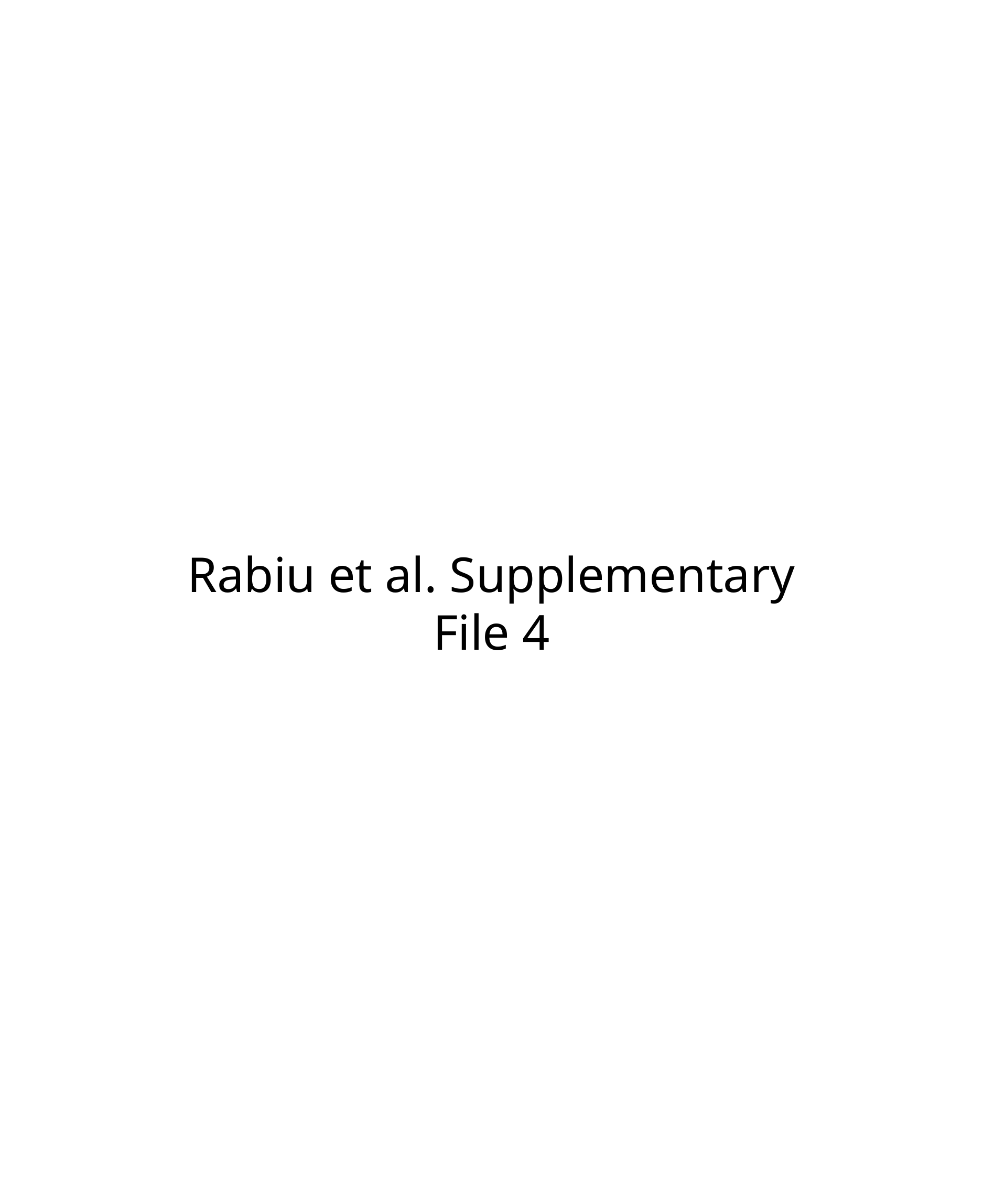

Rabiu et al. Supplementary File 4

### Slide 2
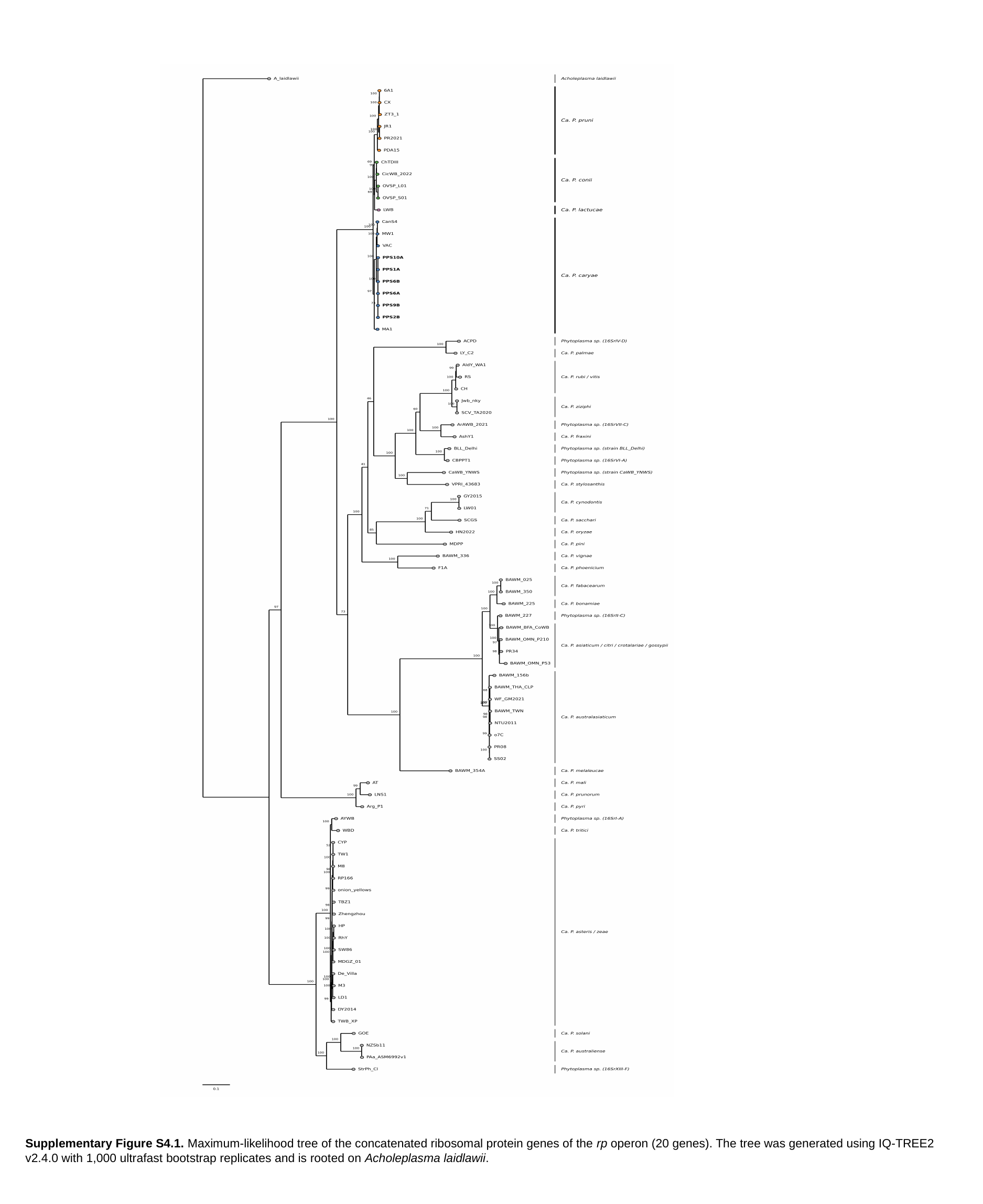

Supplementary Figure S4.1. Maximum-likelihood tree of the concatenated ribosomal protein genes of the rp operon (20 genes). The tree was generated using IQ-TREE2 v2.4.0 with 1,000 ultrafast bootstrap replicates and is rooted on Acholeplasma laidlawii.

### Slide 3
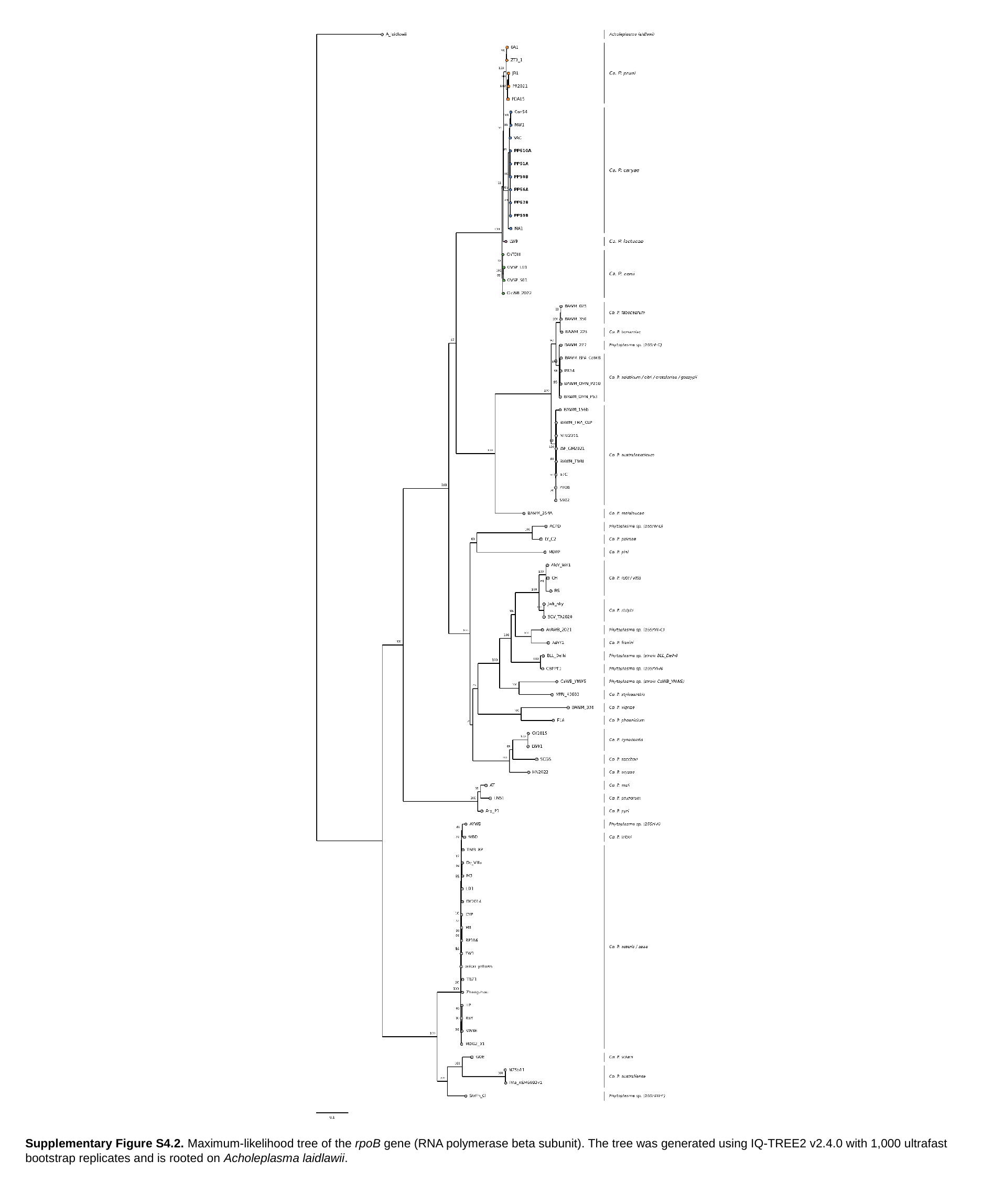

Supplementary Figure S4.2. Maximum-likelihood tree of the rpoB gene (RNA polymerase beta subunit). The tree was generated using IQ-TREE2 v2.4.0 with 1,000 ultrafast bootstrap replicates and is rooted on Acholeplasma laidlawii.

### Slide 4
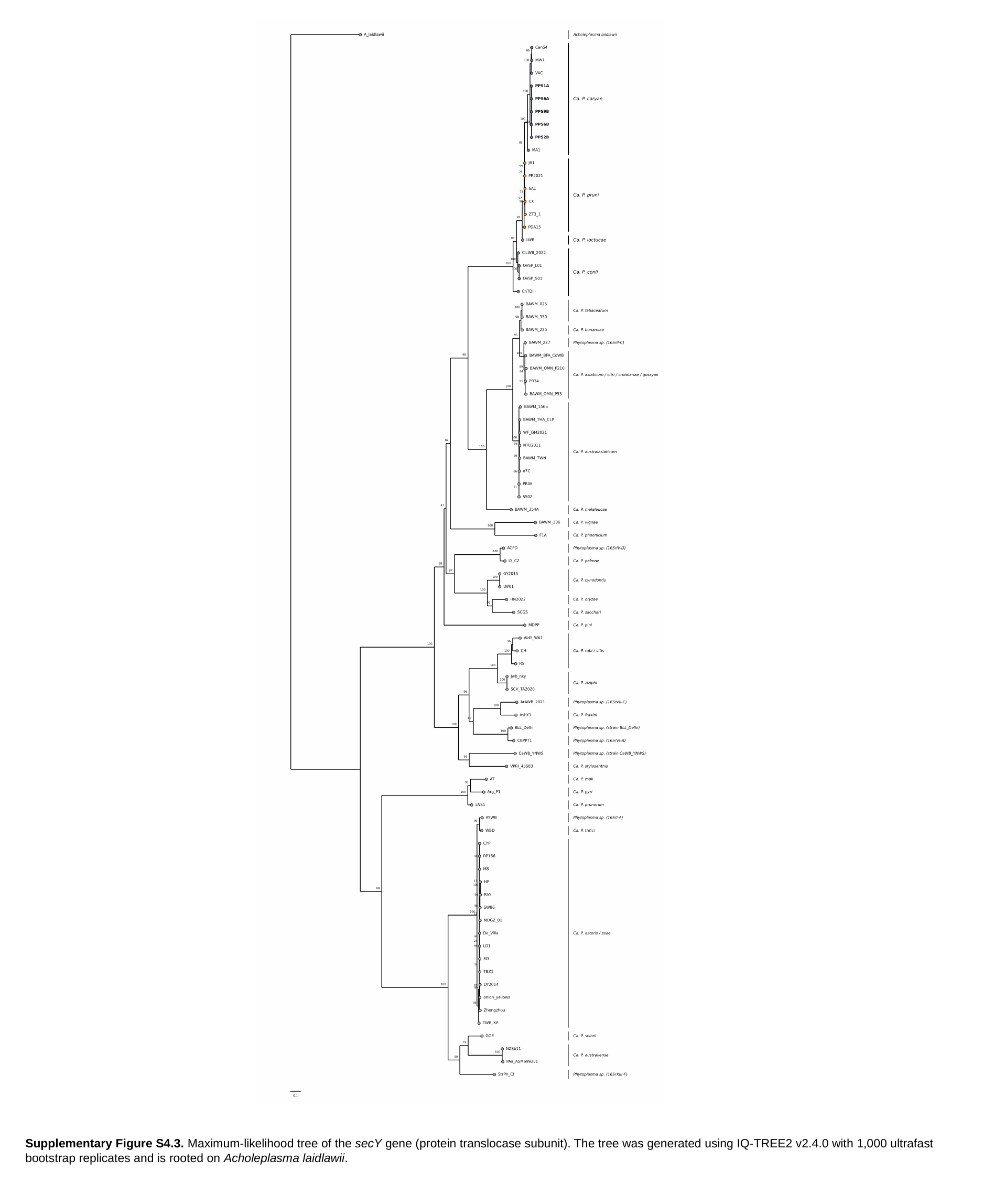

Supplementary Figure S4.3. Maximum-likelihood tree of the secY gene (protein translocase subunit). The tree was generated using IQ-TREE2 v2.4.0 with 1,000 ultrafast bootstrap replicates and is rooted on Acholeplasma laidlawii.

### Slide 5
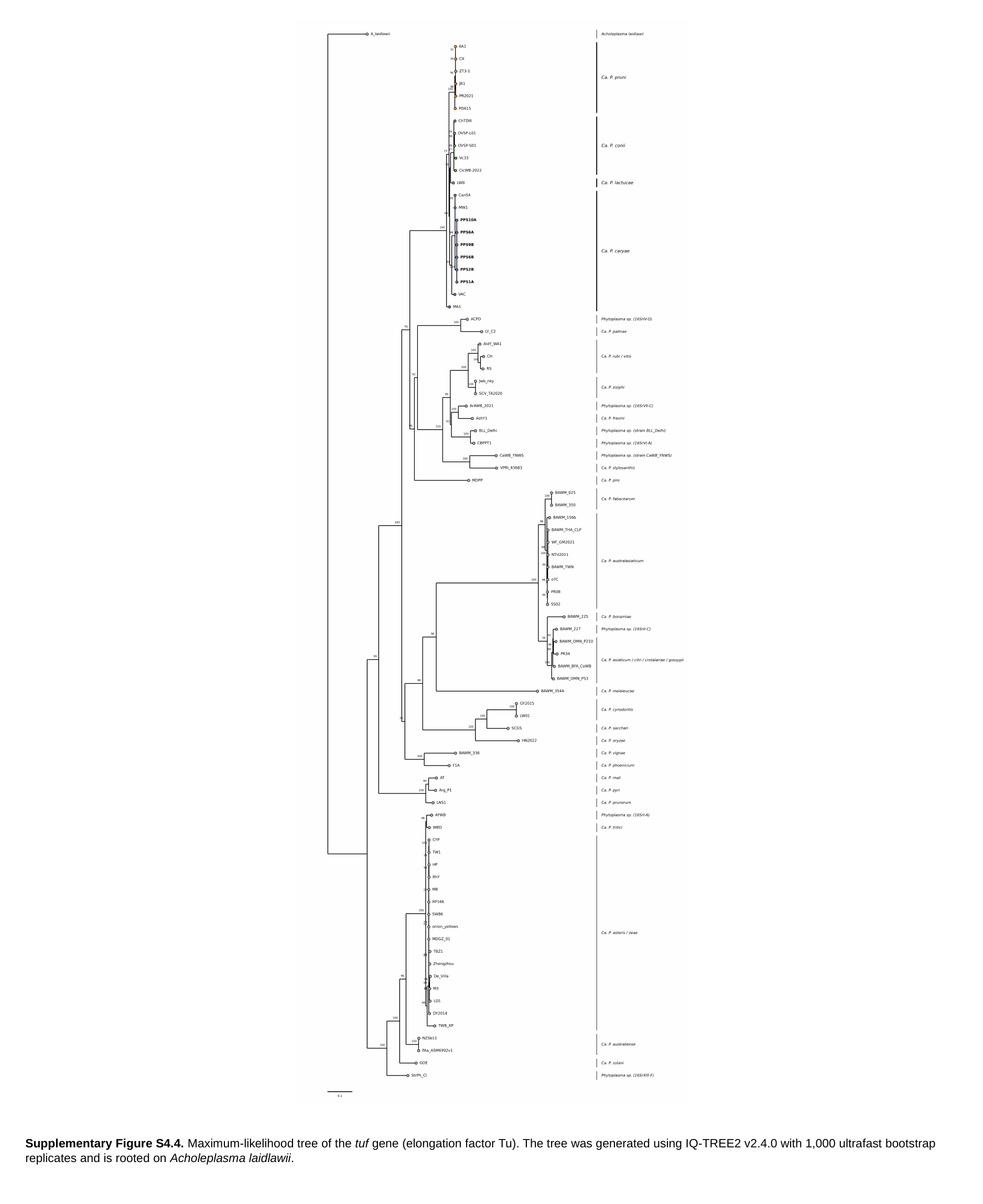

Supplementary Figure S4.4. Maximum-likelihood tree of the tuf gene (elongation factor Tu). The tree was generated using IQ-TREE2 v2.4.0 with 1,000 ultrafast bootstrap replicates and is rooted on Acholeplasma laidlawii.

### Slide 6
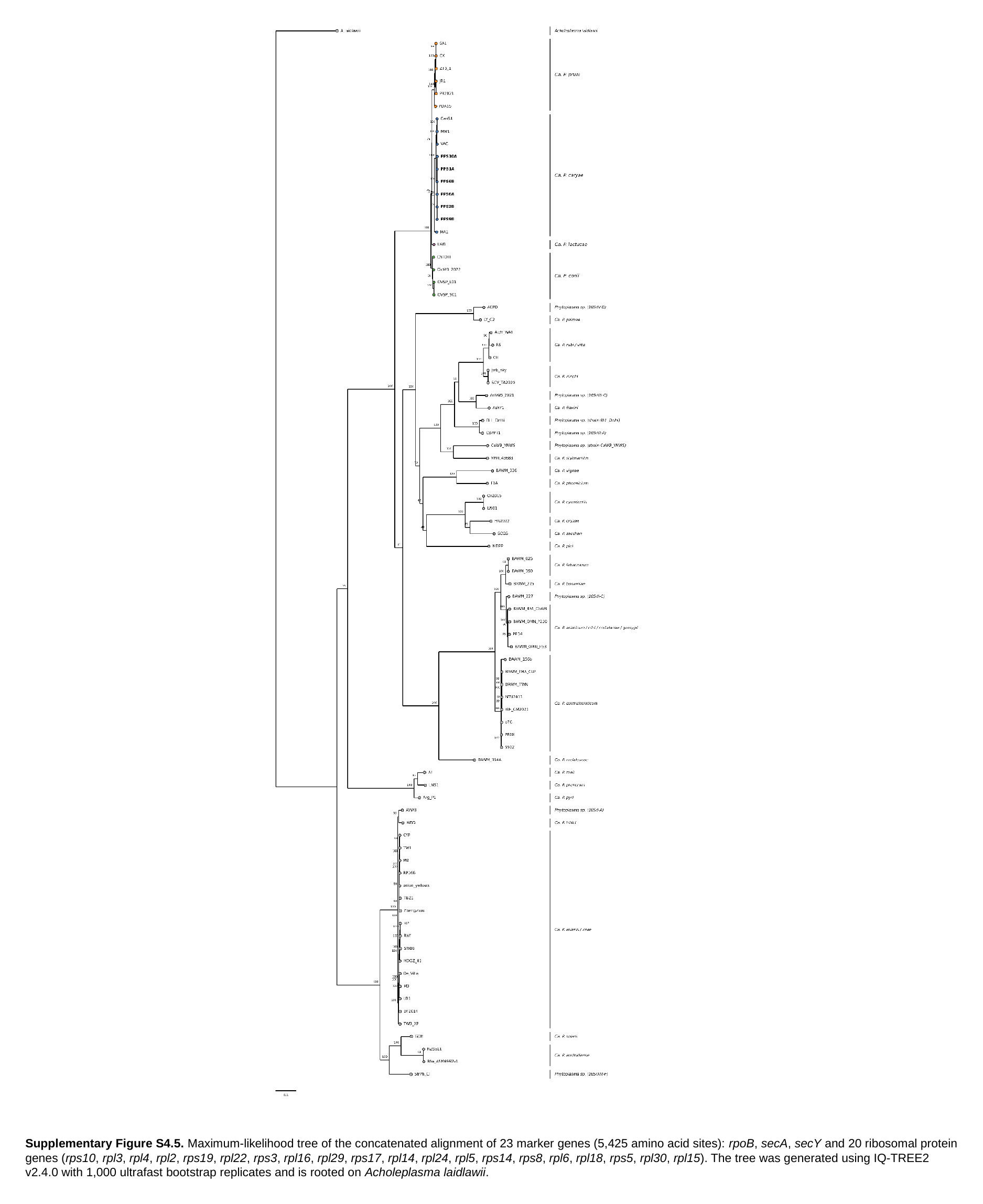

Supplementary Figure S4.5. Maximum-likelihood tree of the concatenated alignment of 23 marker genes (5,425 amino acid sites): rpoB, secA, secY and 20 ribosomal protein genes (rps10, rpl3, rpl4, rpl2, rps19, rpl22, rps3, rpl16, rpl29, rps17, rpl14, rpl24, rpl5, rps14, rps8, rpl6, rpl18, rps5, rpl30, rpl15). The tree was generated using IQ-TREE2 v2.4.0 with 1,000 ultrafast bootstrap replicates and is rooted on Acholeplasma laidlawii.

### Slide 7
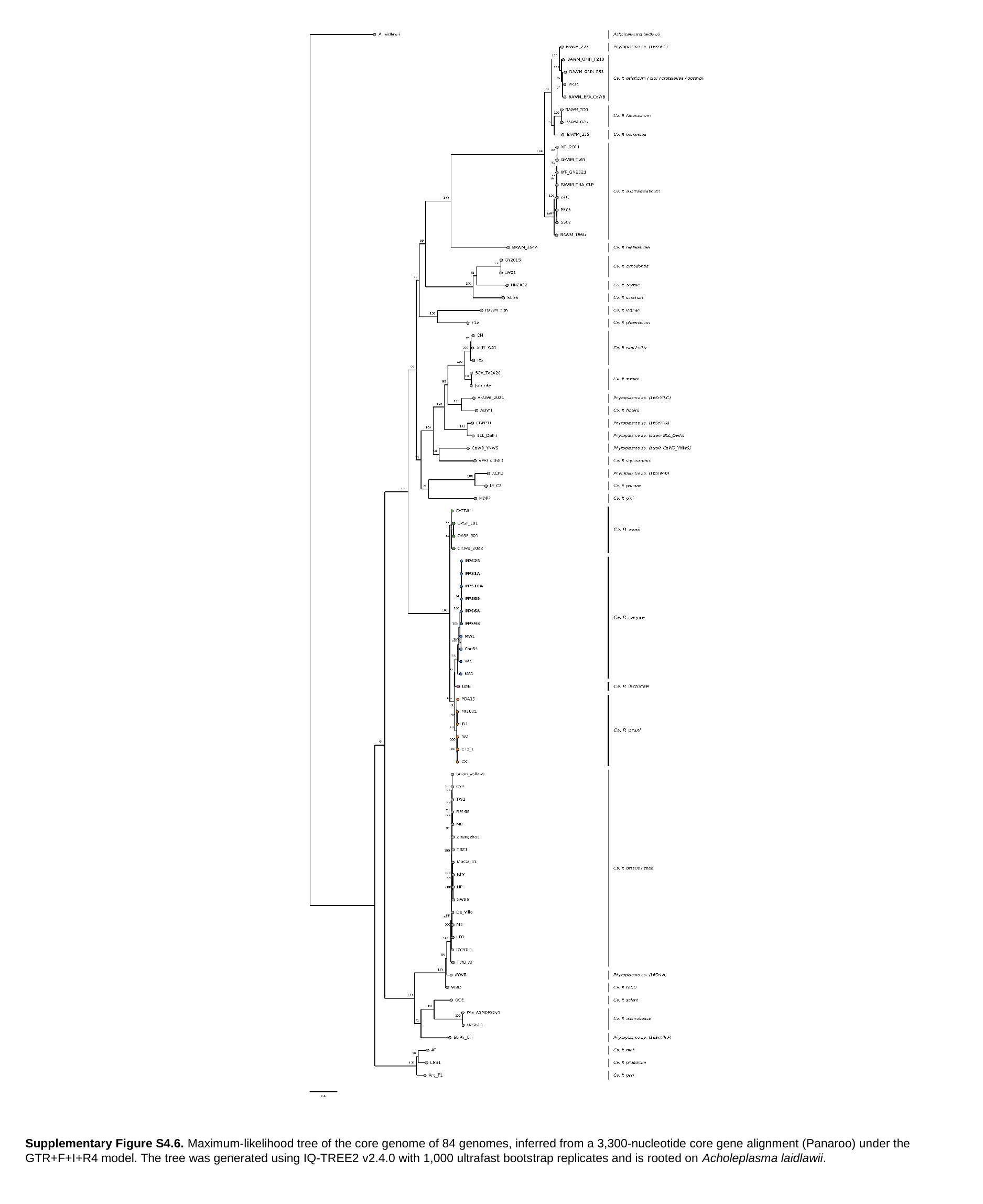

Supplementary Figure S4.6. Maximum-likelihood tree of the core genome of 84 genomes, inferred from a 3,300-nucleotide core gene alignment (Panaroo) under the GTR+F+I+R4 model. The tree was generated using IQ-TREE2 v2.4.0 with 1,000 ultrafast bootstrap replicates and is rooted on Acholeplasma laidlawii.
