## Supplementary material for "Taxogenomic resolution of the 16SrIII (X-disease) group: elevation of ‘*Candidatus* Phytoplasma pruni’-related strains to species rank and proposal of three novel *Candidatus* species ‘*Candidatus* Phytoplasma caryae’ sp. nov., ‘*Candidatus* Phytoplasma conii’ sp. nov. and ‘*Candidatus* Phytoplasma": Rabiu et al., Supplementary figures and Tables: Rabiu_et_al_Supplementary_File_5.pptx

### Slide 1
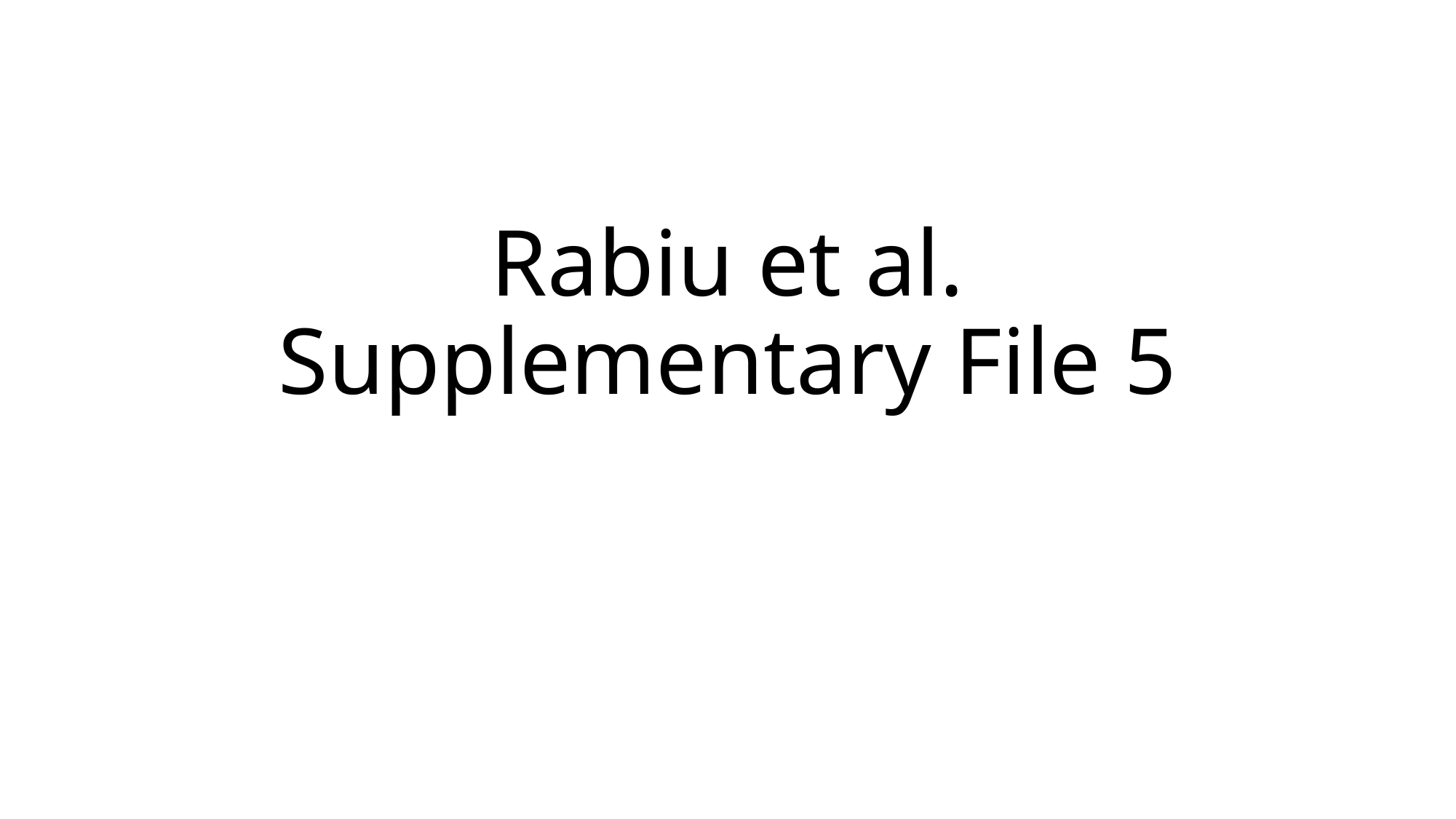

#### Rabiu et al. Supplementary File 5

### Slide 2
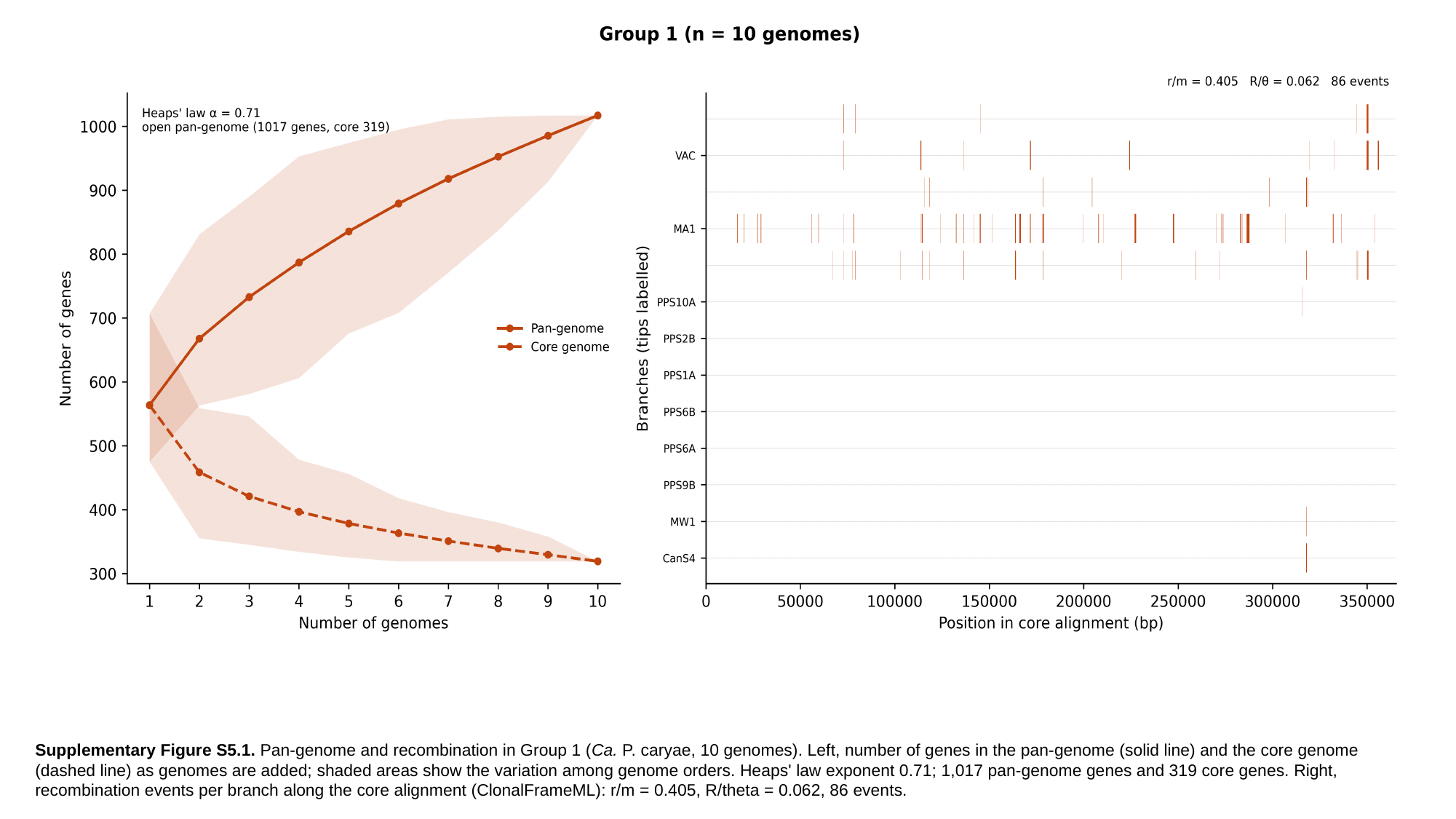

Supplementary Figure S5.1. Pan-genome and recombination in Group 1 (Ca. P. caryae, 10 genomes). Left, number of genes in the pan-genome (solid line) and the core genome (dashed line) as genomes are added; shaded areas show the variation among genome orders. Heaps' law exponent 0.71; 1,017 pan-genome genes and 319 core genes. Right, recombination events per branch along the core alignment (ClonalFrameML): r/m = 0.405, R/theta = 0.062, 86 events.

### Slide 3
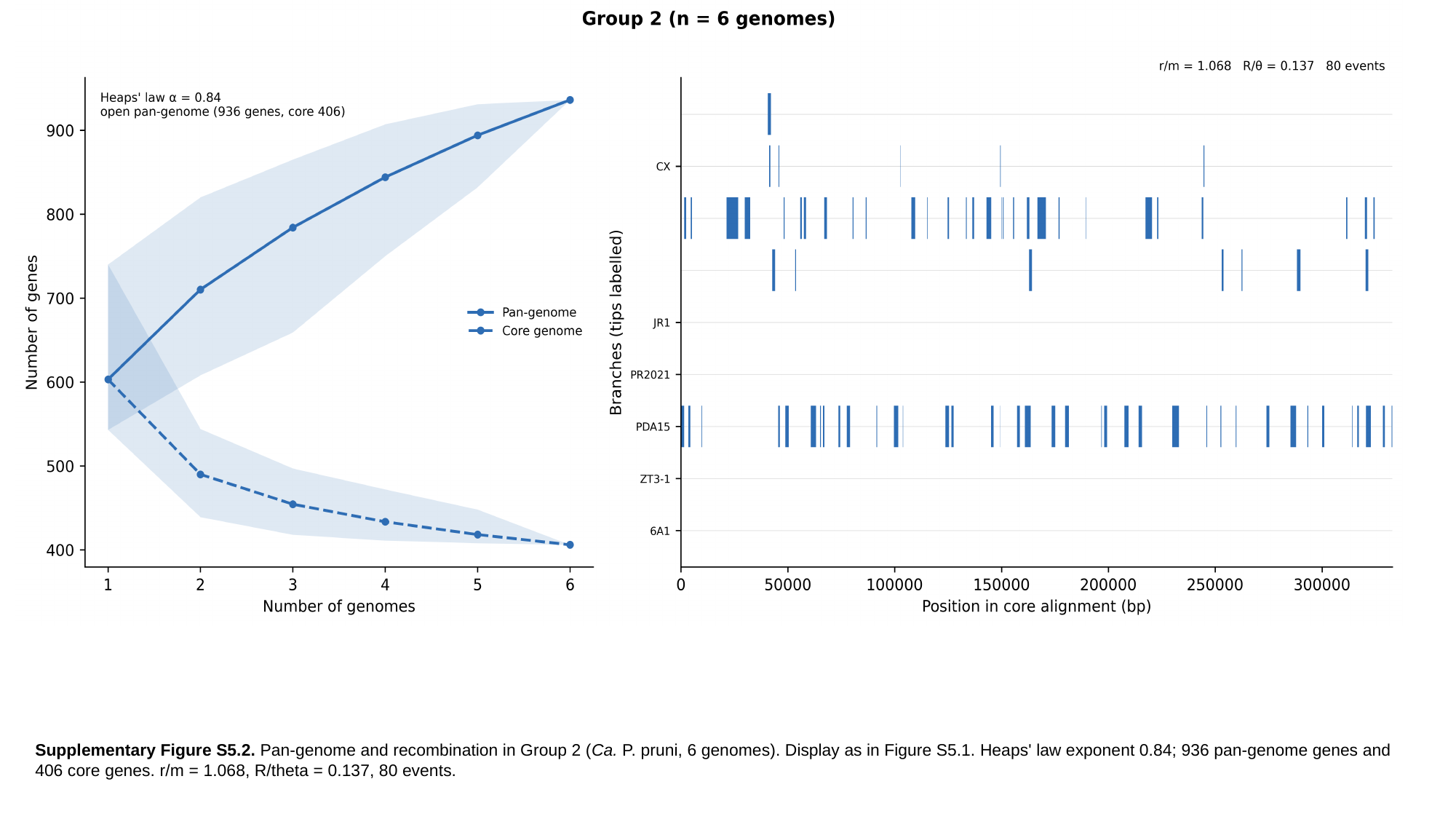

Supplementary Figure S5.2. Pan-genome and recombination in Group 2 (Ca. P. pruni, 6 genomes). Display as in Figure S5.1. Heaps' law exponent 0.84; 936 pan-genome genes and 406 core genes. r/m = 1.068, R/theta = 0.137, 80 events.

### Slide 4
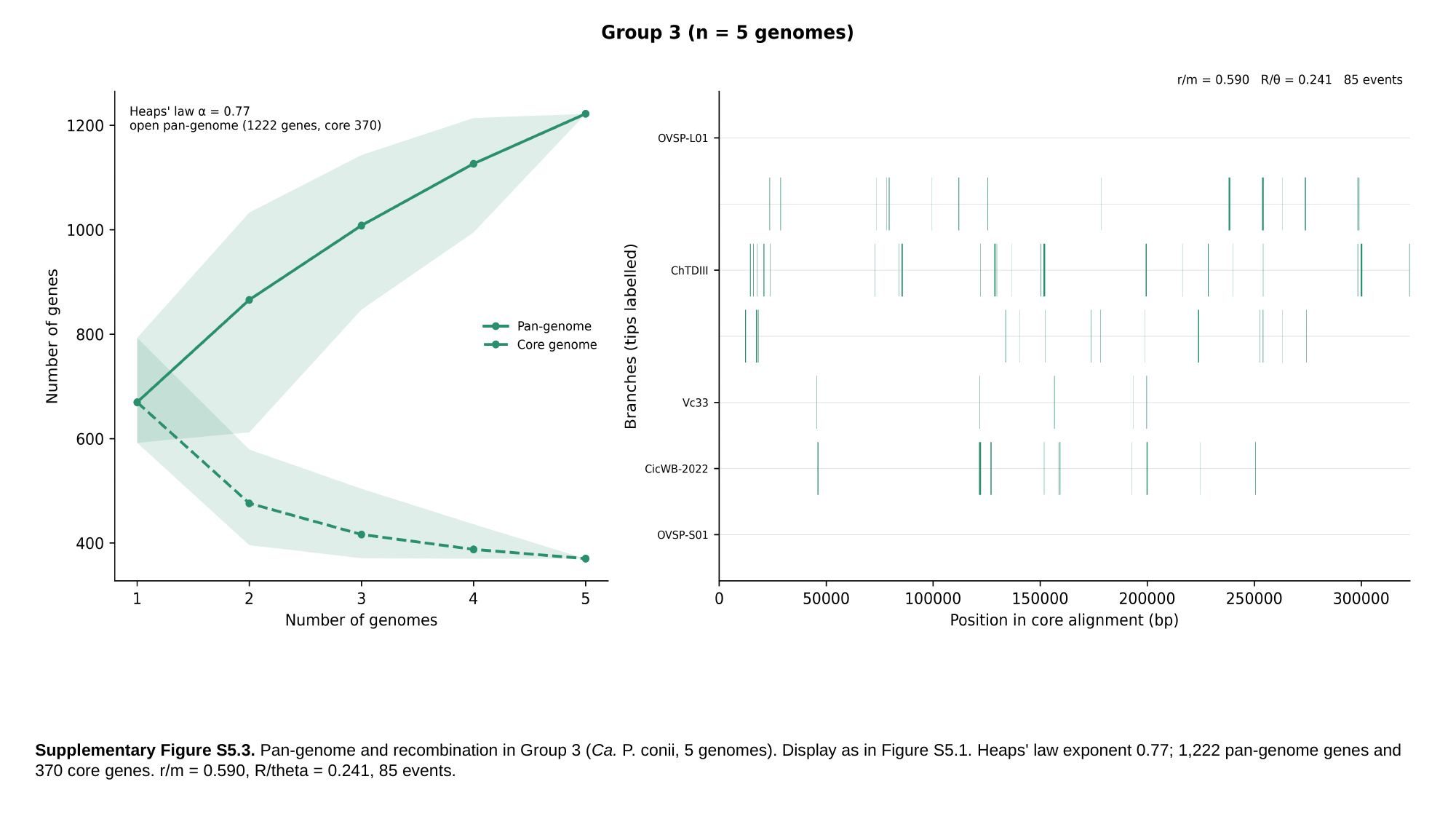

Supplementary Figure S5.3. Pan-genome and recombination in Group 3 (Ca. P. conii, 5 genomes). Display as in Figure S5.1. Heaps' law exponent 0.77; 1,222 pan-genome genes and 370 core genes. r/m = 0.590, R/theta = 0.241, 85 events.

### Slide 5
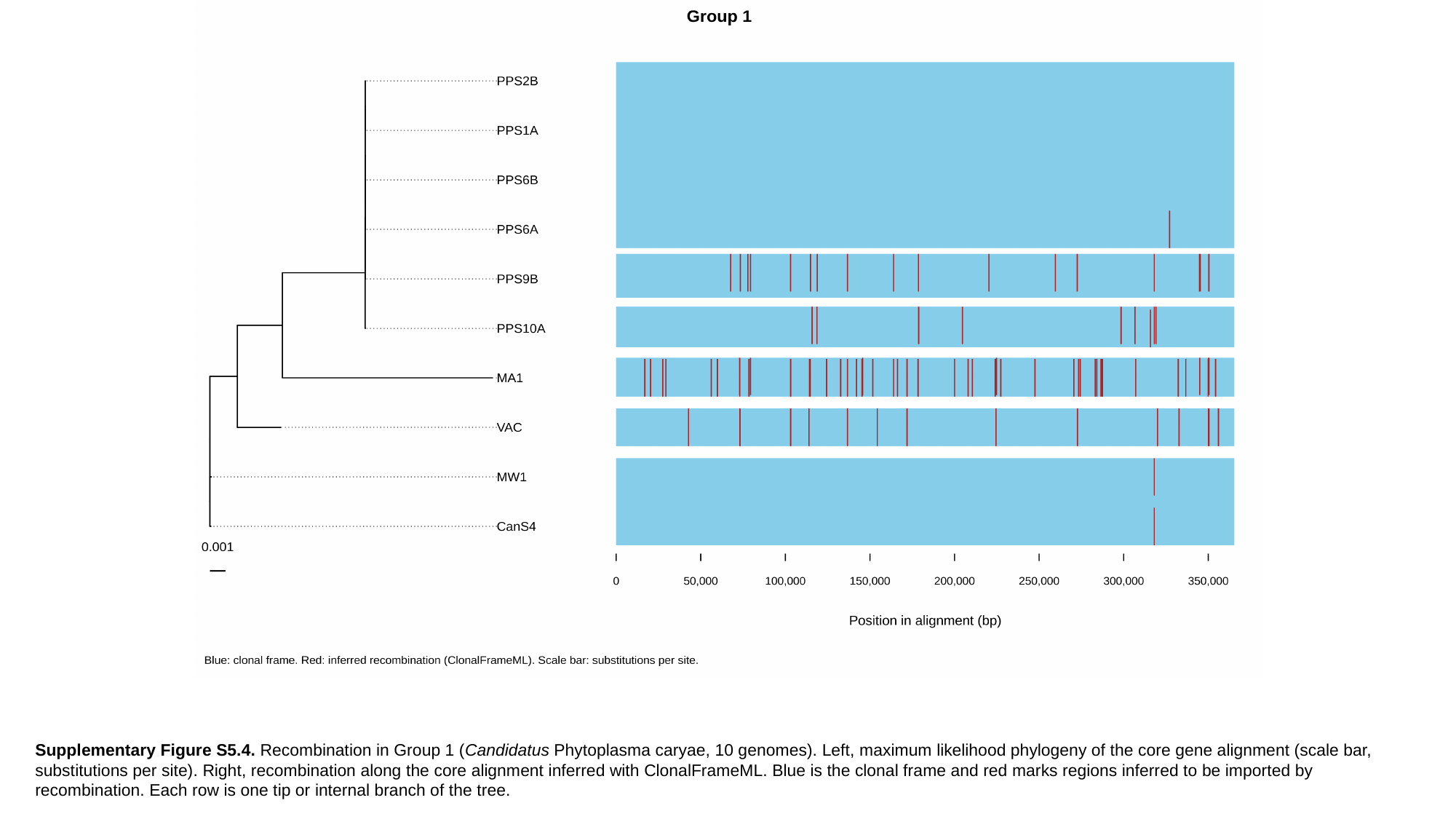

Supplementary Figure S5.4. Recombination in Group 1 (Candidatus Phytoplasma caryae, 10 genomes). Left, maximum likelihood phylogeny of the core gene alignment (scale bar, substitutions per site). Right, recombination along the core alignment inferred with ClonalFrameML. Blue is the clonal frame and red marks regions inferred to be imported by recombination. Each row is one tip or internal branch of the tree.

### Slide 6
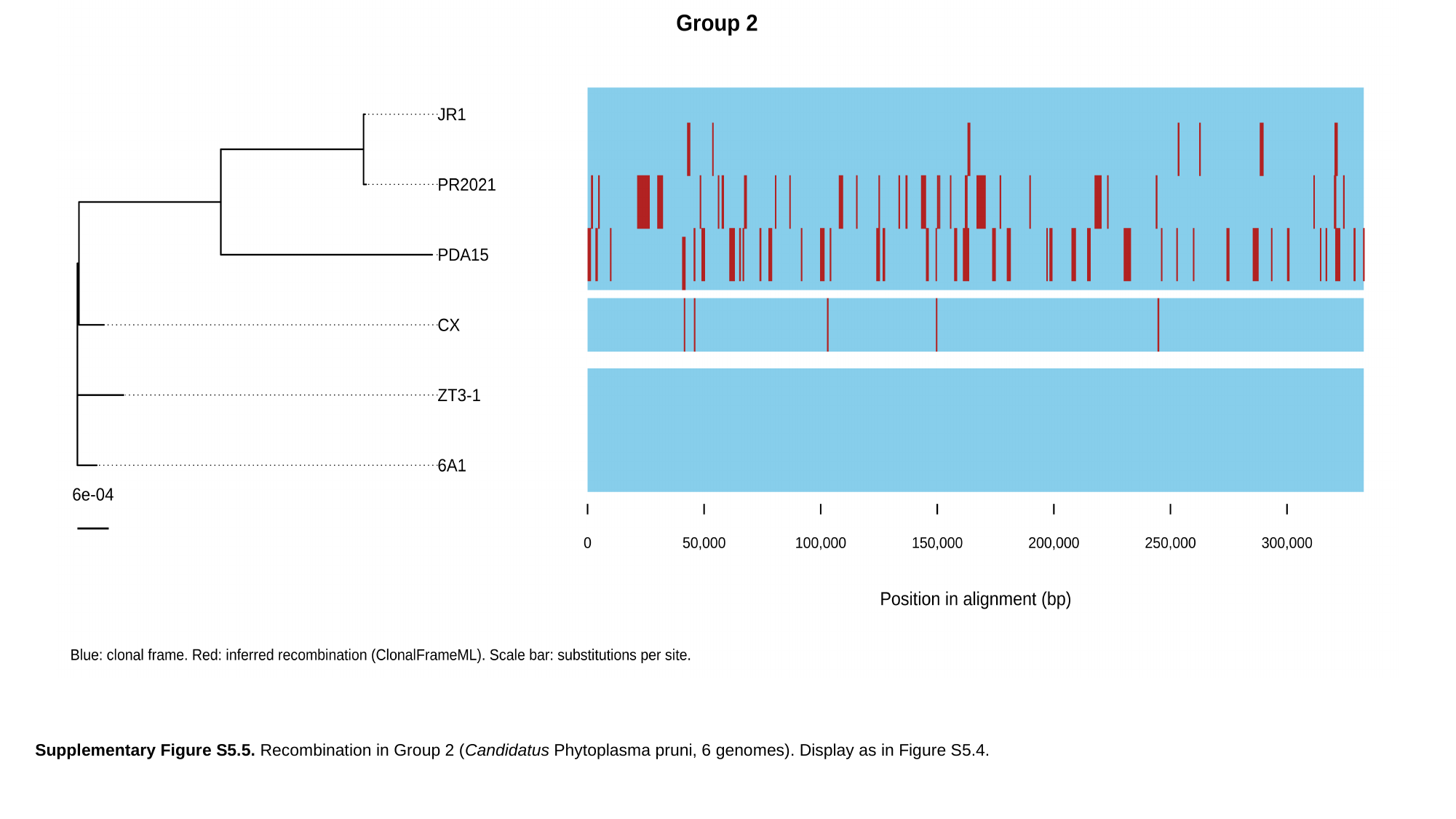

Supplementary Figure S5.5. Recombination in Group 2 (Candidatus Phytoplasma pruni, 6 genomes). Display as in Figure S5.4.

### Slide 7
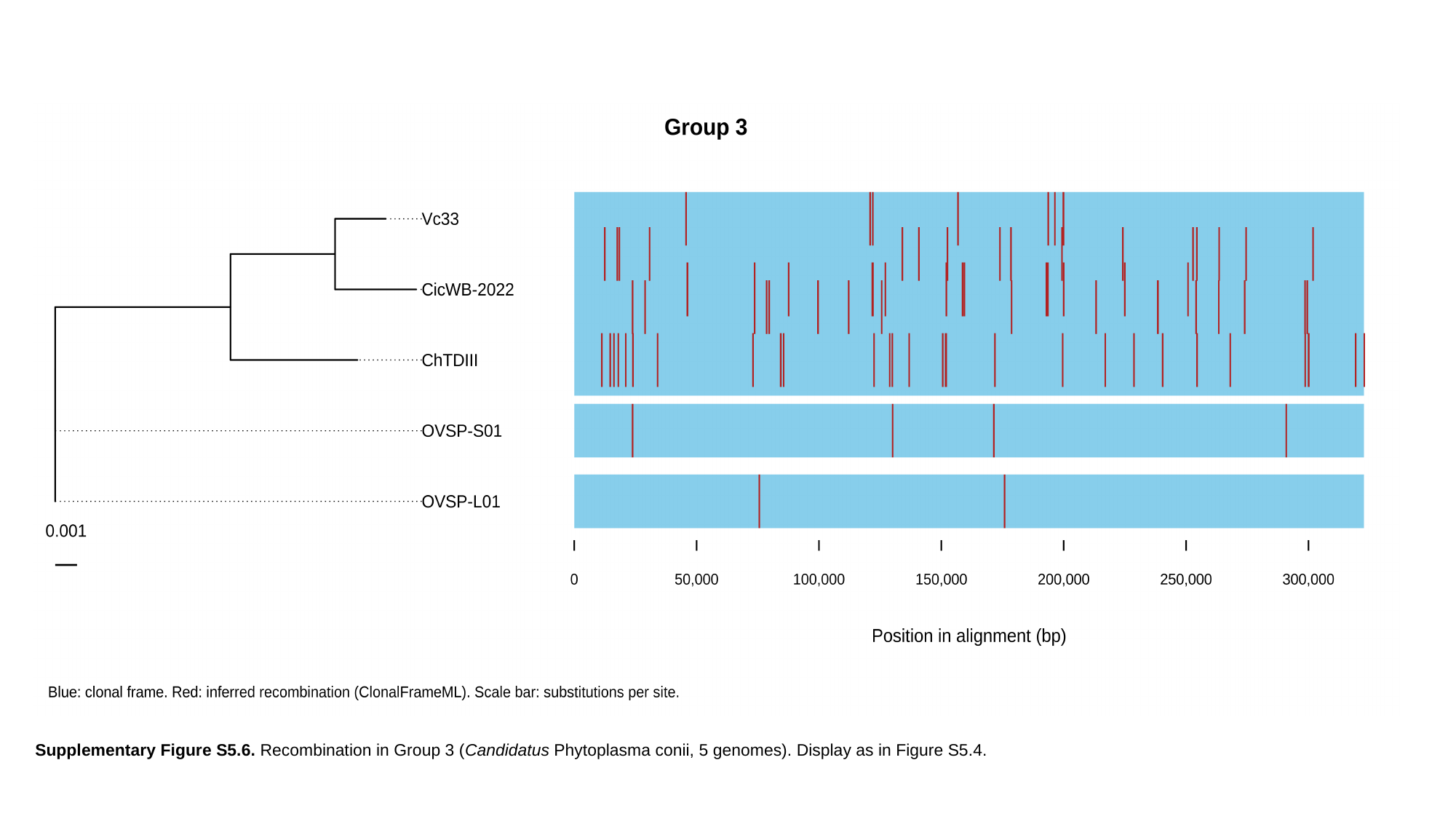

Supplementary Figure S5.6. Recombination in Group 3 (Candidatus Phytoplasma conii, 5 genomes). Display as in Figure S5.4.
