## Supplementary material for "Taxogenomic resolution of the 16SrIII (X-disease) group: elevation of ‘*Candidatus* Phytoplasma pruni’-related strains to species rank and proposal of three novel *Candidatus* species ‘*Candidatus* Phytoplasma caryae’ sp. nov., ‘*Candidatus* Phytoplasma conii’ sp. nov. and ‘*Candidatus* Phytoplasma": Rabiu et al., Supplementary figures and Tables: Rabiu_et_al_Supplementary_File_6.pptx

### Slide 1
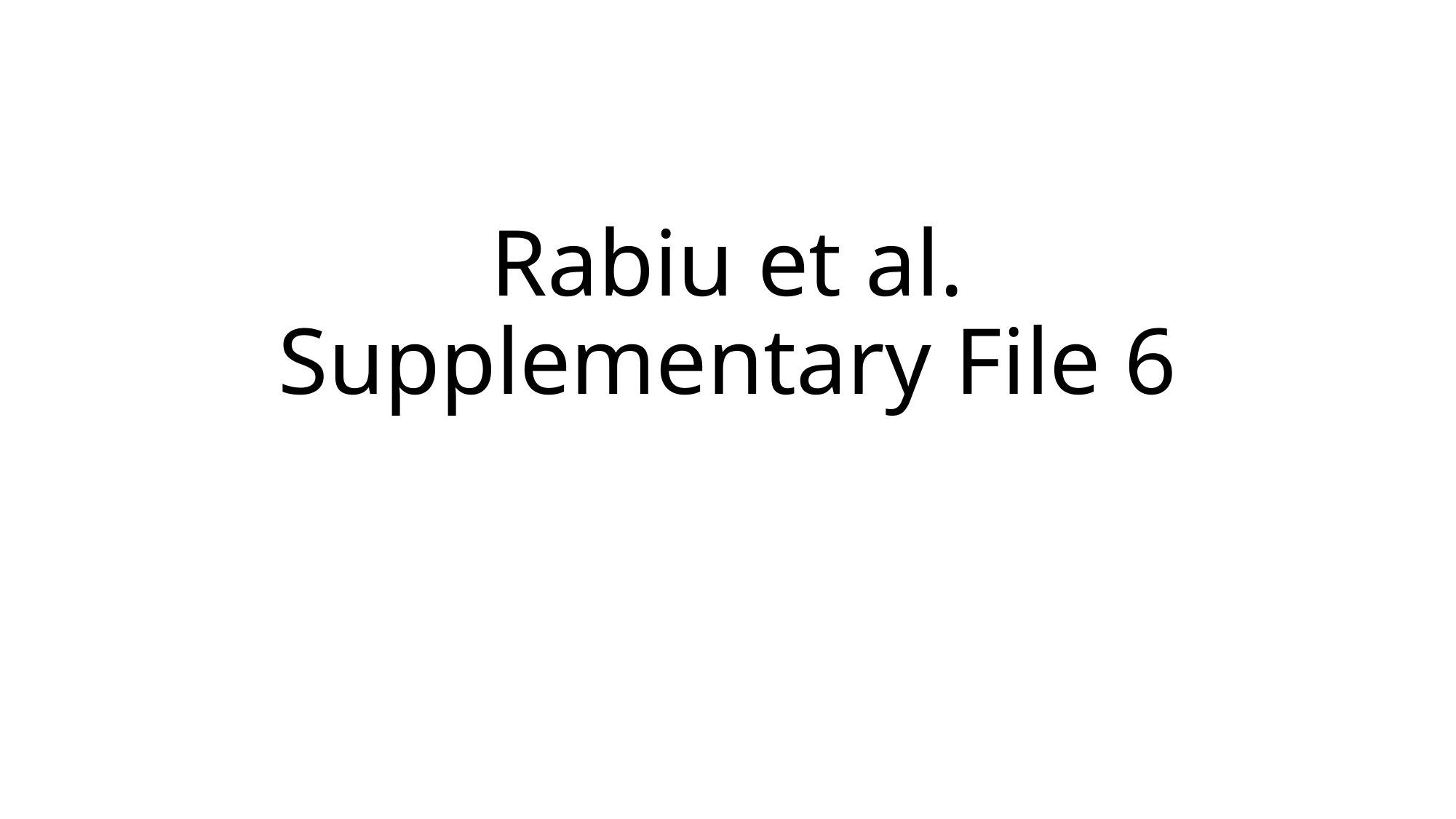

#### Rabiu et al. Supplementary File 6

### Slide 2
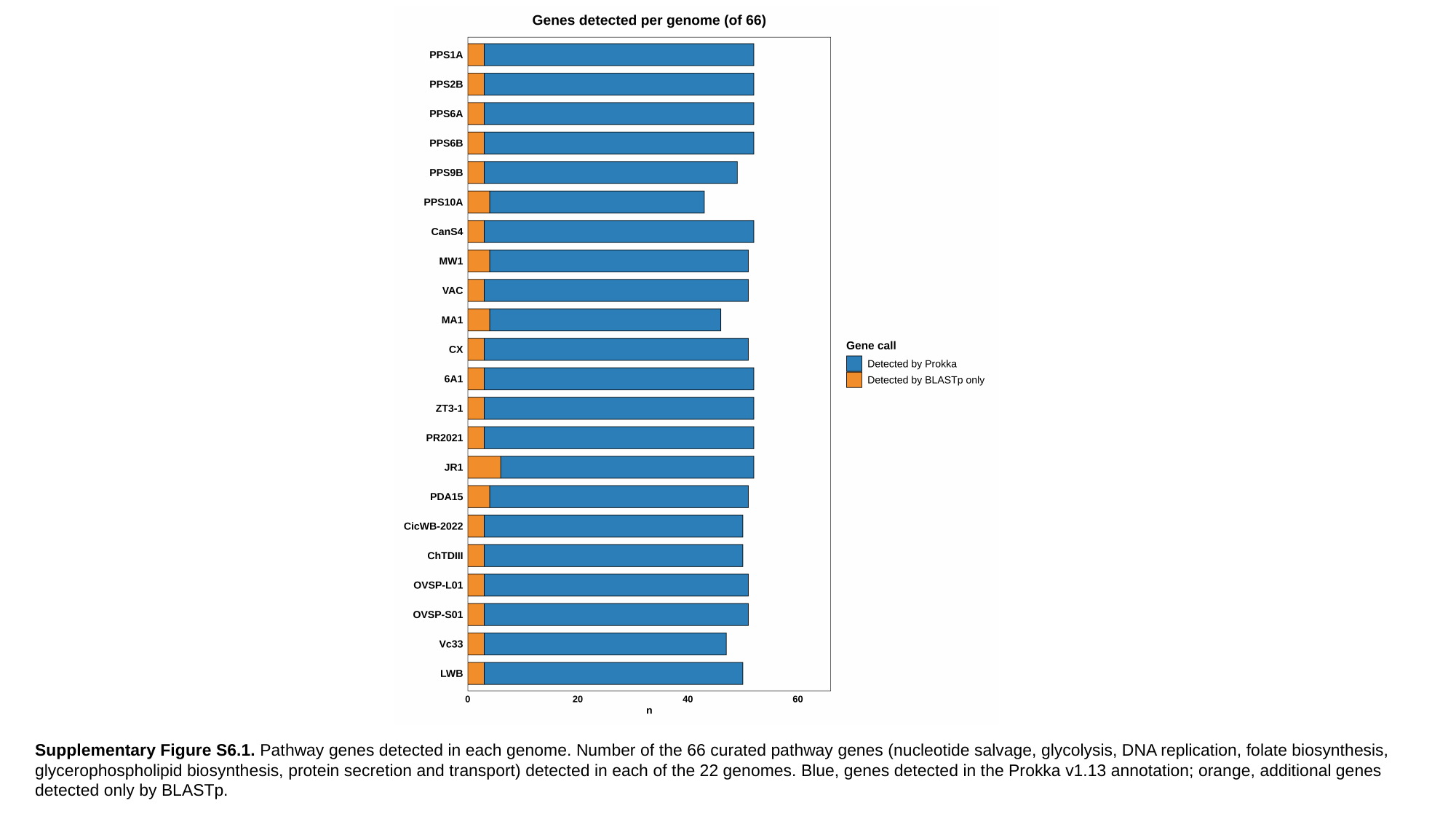

Supplementary Figure S6.1. Pathway genes detected in each genome. Number of the 66 curated pathway genes (nucleotide salvage, glycolysis, DNA replication, folate biosynthesis, glycerophospholipid biosynthesis, protein secretion and transport) detected in each of the 22 genomes. Blue, genes detected in the Prokka v1.13 annotation; orange, additional genes detected only by BLASTp.

### Slide 3
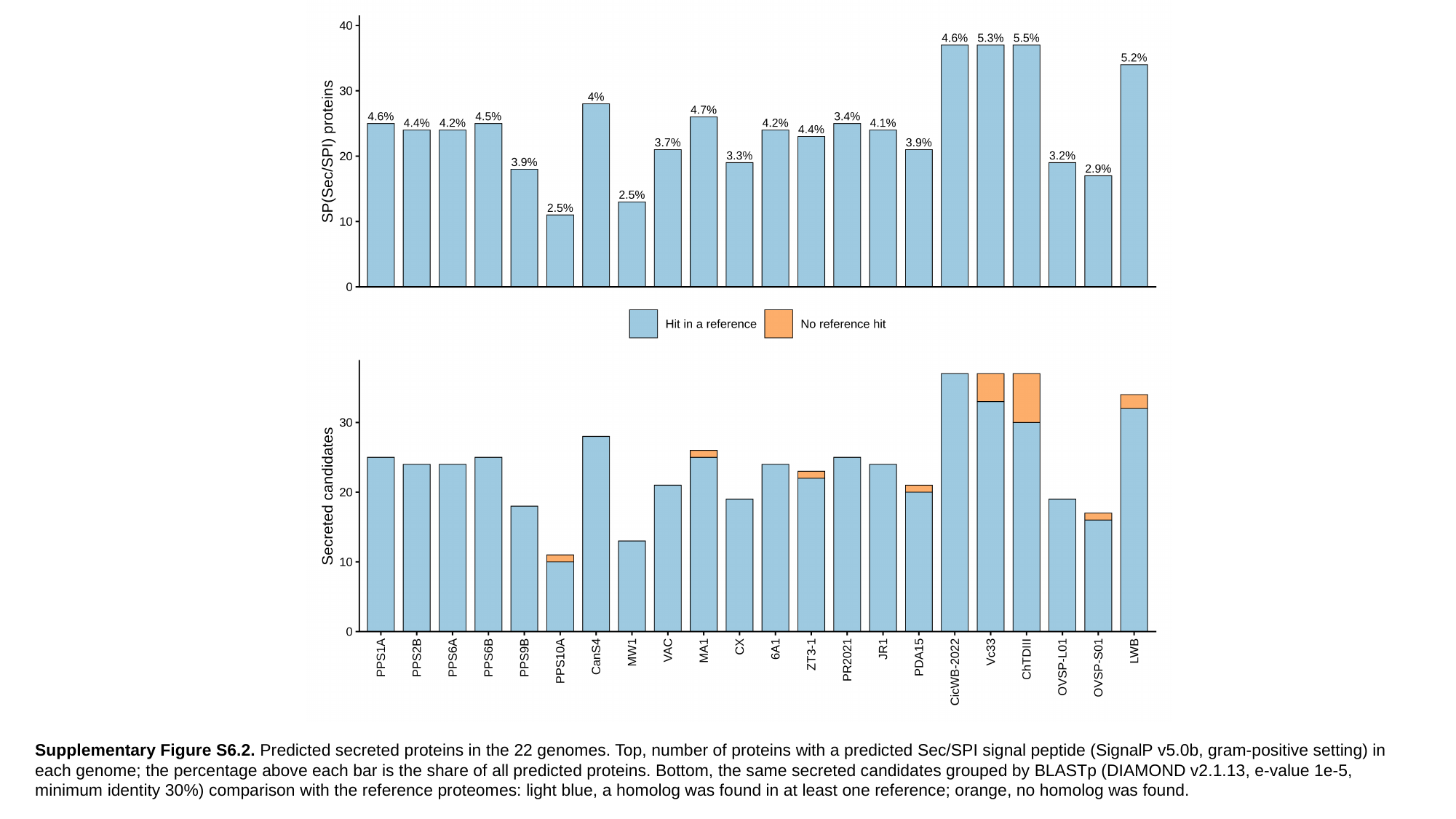

Supplementary Figure S6.2. Predicted secreted proteins in the 22 genomes. Top, number of proteins with a predicted Sec/SPI signal peptide (SignalP v5.0b, gram-positive setting) in each genome; the percentage above each bar is the share of all predicted proteins. Bottom, the same secreted candidates grouped by BLASTp (DIAMOND v2.1.13, e-value 1e-5, minimum identity 30%) comparison with the reference proteomes: light blue, a homolog was found in at least one reference; orange, no homolog was found.

### Slide 4
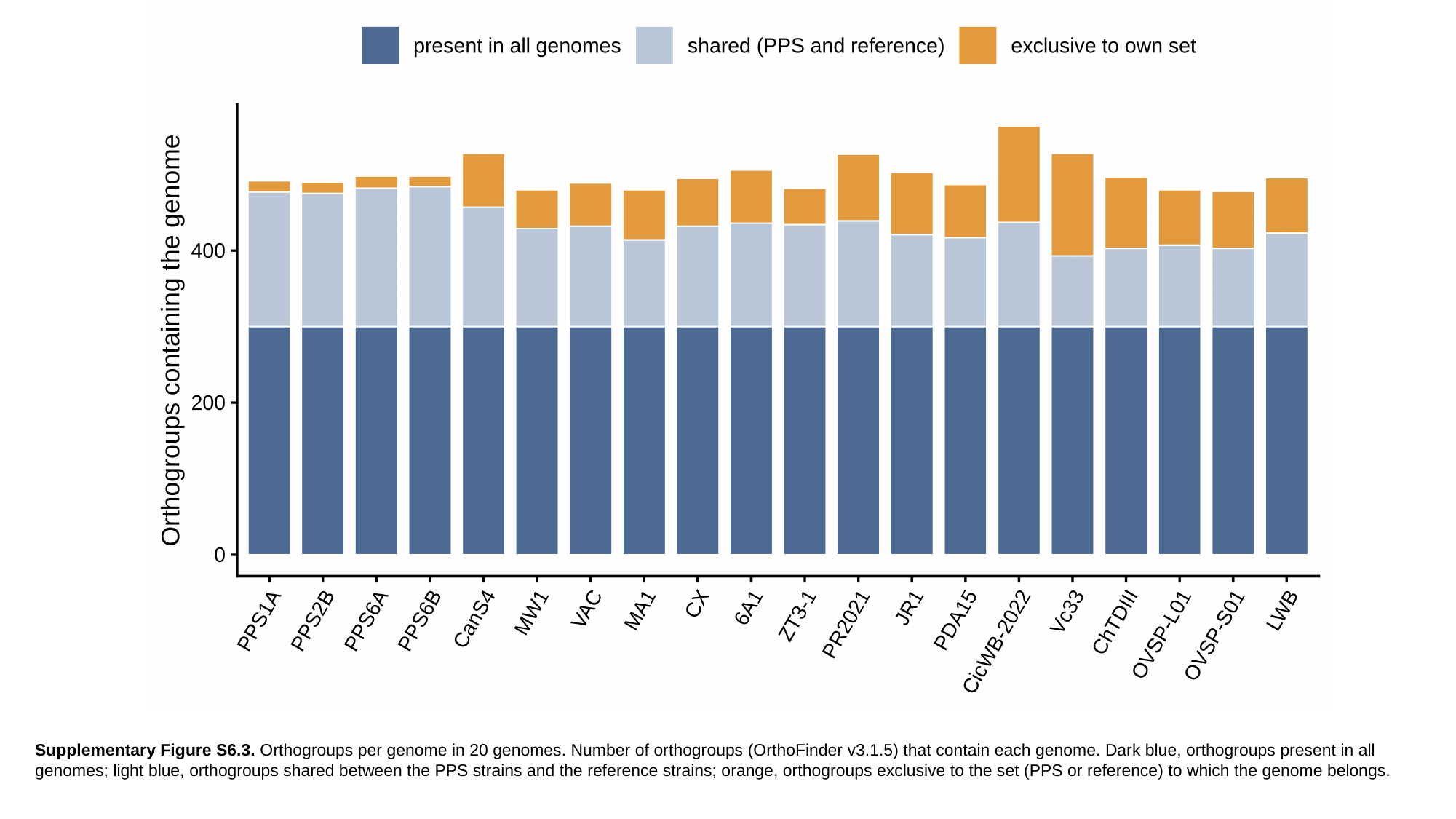

Supplementary Figure S6.3. Orthogroups per genome in 20 genomes. Number of orthogroups (OrthoFinder v3.1.5) that contain each genome. Dark blue, orthogroups present in all genomes; light blue, orthogroups shared between the PPS strains and the reference strains; orange, orthogroups exclusive to the set (PPS or reference) to which the genome belongs.

### Slide 5
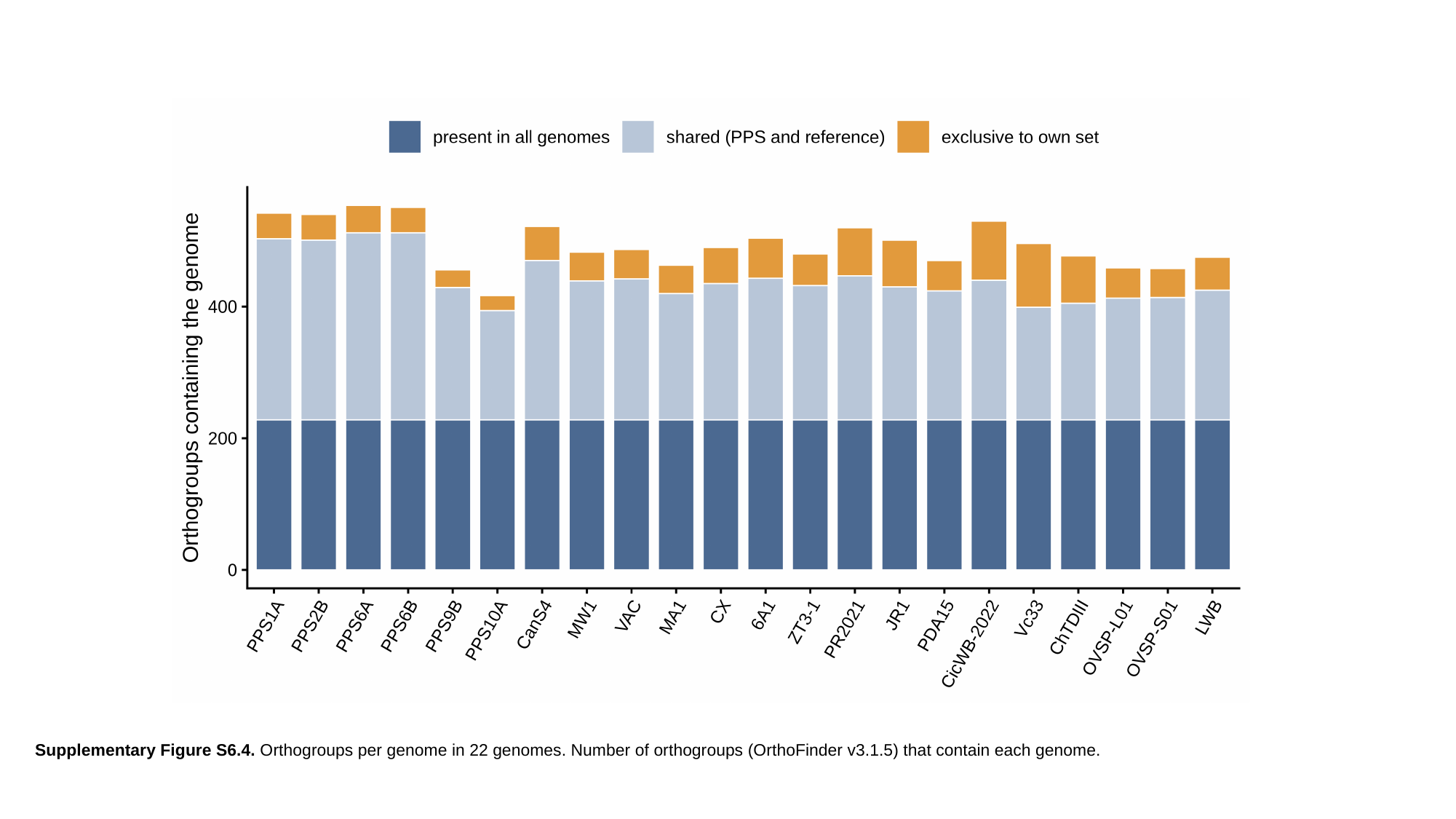

Supplementary Figure S6.4. Orthogroups per genome in 22 genomes. Number of orthogroups (OrthoFinder v3.1.5) that contain each genome.

### Slide 6
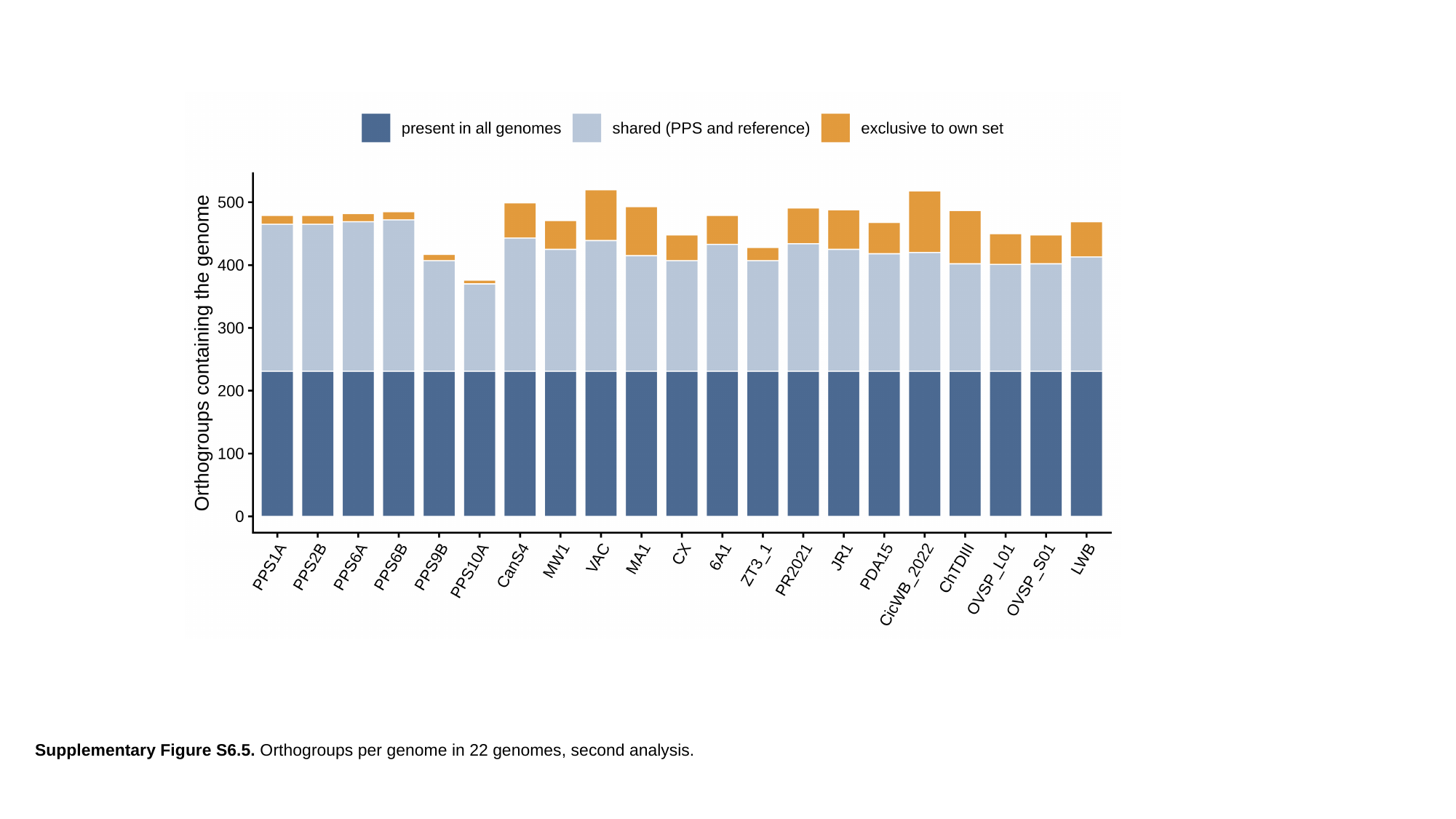

Supplementary Figure S6.5. Orthogroups per genome in 22 genomes, second analysis.
